# Direct and diffuse cross-kingdom interactions in plant microbiome assembly

**DOI:** 10.1101/2025.10.24.684285

**Authors:** Rachel A. Hammer, Marissa R. Lee, Jeffrey A. Kimbrel, Rhona K. Stuart, Christine V. Hawkes

## Abstract

Studies of plant-associated microbial communities consistently indicate a role for classic assembly mechanisms, such as environmental and host filters, but often leave substantial unexplained variation. Biotic interactions within microbial communities may help to fill this gap, specifically cross-kingdom interactions between fungi and bacteria, as these are increasingly found to be important to both assembly and function. We hypothesized that direct interactions between bacteria and fungi are an important driver of composition in low-diversity leaf habitats, where pairwise interactions are more likely. In high-diversity root habitats, we expected diffuse, indirect interactions to be more relevant to composition. To test these hypotheses, we characterized bacterial and fungal communities of switchgrass (*Panicum virgatum* L.) leaves and roots at 14 sites spanning mountain to coastal ecoregions of North Carolina, USA. We analyzed putative direct and diffuse interactions using ecological network inference and partitioned variance explained in microbial community composition by spatial, environmental, and biotic interactions. We found that cross-kingdom biotic interactions contributed to microbial community structure. The largest improvements to variance explained (5-11%) were from direct interactions, except for root fungal communities where diffuse interactions (7.5%) explained more than double that of direct interactions (2.8%). These contributions were comparable to those from environmental and spatial factors. The joint effects of putative biotic interactions and environmental conditions also contributed to the explained variation, highlighting the importance of environmental tracking in microbes. These findings suggest that using network inference for identifying cross-kingdom ecological interactions can improve our fundamental understanding of how plant-associated microbiomes assemble, which is also directly relevant to applied efforts such as the effective development of synthetic communities.

## INTRODUCTION

Studies of plant-associated microbial communities consistently indicate a role for classic assembly mechanisms, such as environmental and host filters, but often leave substantial unexplained variation (Mészárošová et al. 2024; Lee and Hawkes 2021a). Here, we propose that this knowledge gap may be partly filled by considering biotic interactions, particularly between fungi and bacteria. Advances in statistical and mechanistic modeling have expanded the capacity to infer biotic interactions and their role in community assembly from large, high-dimensional ecological datasets (Layeghifard et al. 2017; Muller et al. 2018; Pichler and Hartig 2021). These advances have facilitated our ability to consider biotic interactions and uncover key microbes that respond both to abiotic factors and each other (e.g., Agler et al. 2016).

To date, the vast majority of experimental microbiome studies address biotic interactions pairwise, which assumes that interactions are direct and independent of other taxa in the community. Yet emergent properties arising from indirect interactions, which are generally unpredictable from pairwise effects, are common in both plant and animal communities (Lai et al. 2022; Werner and Peacor 2003) and their occurrence in microbial communities is supported by both empirical evidence and models (Sanchez-Gorostiaga et al. 2019). Indirect effects occur when the interaction between two species is mediated via a third species by, for example, interaction chains, modification of environmental conditions, or changes in resource availability (Wootton 1994). Indirect interactions in multispecies communities may be diffuse if the effect of one species is spread over many other species (Moen 1989). In microbial communities, such diffuse effects might occur through production of extracellular enzymes, signaling molecules, and other metabolites (Weiland-Bräuer 2021). In plant-associated microbial communities, diffuse effects might also include modifications to the host environment (Seybold et al. 2020).

Cross-kingdom interactions may be as important as within-kingdom interactions in microbial communities (Lee et al. 2022), including both direct and indirect effects. Bacteria and fungi often live in close physical associations that can facilitate interactions, such as in mixed biofilms within soil micro-niches and the rhizosphere, where both competition and facilitation can occur. For instance, mycorrhizal hyphae are known to harbor bacteria internally, on the hyphal surface, and in the hyphosphere, where recent studies point to carbon-nutrient exchange (Duan et al. 2023) and facilitation of stress tolerance (Hestrin et al. 2022). Indirect interactions are less studied, but a clear example was reported by Sharma et al. (2021), in which a nitrogen-fixing bacterium modified the interaction between a fungus and a fungus-feeding bacterium. Despite evidence supporting cross-kingdom interactions (Yuan et al. 2021), these are rarely incorporated into assessments of community assembly and our understanding of microbial interaction effects on assembly therefore remain limited.

To examine the role of both direct and indirect (diffuse) cross-kingdom interactions in microbiome community assembly, we characterized bacterial and fungal communities associated with switchgrass (*Panicum virgatum* L.) at 14 stands spanning mountain to coastal ecoregions of North Carolina, USA. We inferred putative cross-kingdom direct and diffuse interactions using ecological networks. Using variance partitioning analysis, we examined how direct and diffuse cross-kingdom interactions obtained from the networks contributed to community assembly relative to spatial and environmental factors. We hypothesized that direct interactions between bacterial and fungal taxa would be an important driver of leaf microbiome composition, because these are oligotrophic, low-diversity habitats and microbes tend to aggregate in more favorable microsites (Chaudhry et al. 2020), making pairwise interactions more likely. In roots, however, we expected diffuse, indirect interactions to be more relevant to composition, because these are exudate-rich habitats with high microbial diversity, density, and activity (Huang et al. 2014).

## METHODS

### Experimental design and study sites

Switchgrass (*Panicum virgatum* L.) was sampled at 14 sites across a 462-km range in North Carolina, USA, in September and October 2018, as described in Lee and Hawkes (Lee and Hawkes 2021a). Briefly, we sampled 8 plants per site in a stratified random design. At each plant, we collected 1) five healthy, undamaged leaves spaced throughout the plant canopy, and 2) roots and soils to 15-cm depth immediately adjacent to the north and south side of the plant. Samples were stored on ice for transport to the lab, where fine roots < 2 mm were separated from soils and washed. Subsamples of leaves, roots, and soils were used for biogeochemical analyses or frozen at −80 °C until DNA extraction.

### Plant and soil properties

Plant measurements included height, basal width and length, and GPS location. In the lab, we measured soil pH, microbial biomass carbon (MBC), moisture, and soil texture as described in Lee and Hawkes (2021b). Total soil C and N were measured by combustion at the North Carolina State University Environmental and Agricultural Testing Service. Other soil nutrients (P, K) were measured at the NC Department of Agriculture and Consumer Services. Climate variables included mean annual precipitation (MAP) and mean annual temperature (MAT) based on 30-year normals from 1981-2010 (PRISM Climate Group 2014).

### Microbial community composition

We extracted DNA from 40-50 mg of leaves or roots using the Synergy 2.0 Plant DNA Extraction Kit (OPS Diagnostics, Lebanon, New Jersey, USA). Two leaf samples were excluded because they were senesced (brown) at collection. Amplicons were generated for bacteria in the 16S rRNA region using the primers 515F and 806R (Caporaso et al. 2011) and for fungi in the 5.8S and ITS2 rRNA region with the primers 5.8S_FUN and ITS4_FUN (Taylor et al. 2016), modified with Illumina adapters. PCR reactions included 1× KAPA Taq Ready Mix (Roche, Pleasanton, CA, USA), 0.2 µM of each primer, 10 ng µL^−1^ of DNA template, and water up to 25 µL. For 16S, we also added 0.25 µM each of universal peptide nucleic acid (PNA) blockers for host mitochondria and plastids (Lundberg et al. 2013). For ITS, we added 0.5 mg mL^−1^ bovine serum albumin and 15 µM of a custom PNA designed for *P. virgatum* (Lee and Hawkes 2021a). Amplicon fragments <50 bp in size were removed using Agencourt AMPure XP beads in a ratio of 1:1.8 (Beckman Coulter, Indianapolis, IN, USA). Negative controls included one pooled blank from DNA extractions and two PCR blanks. Libraries were prepared using the Nextera XT Index Kit (Illumina, San Diego, CA, USA) and submitted sequencing (2 x 250PE) on Illumina MiSeq v2 in the Genomic Sciences Laboratory at North Carolina State University (Raleigh, NC, USA). Raw sequences are available on the NCBI Short Read Archive under BioProject PRJNA648664.

Primers were removed with *cutadapt* v3.3 (Martin 2011). Amplicon sequence variants were identified using *dada2* v1.10.1, (Callahan et al. 2016) with maxEE=c(2,5) and otherwise default settings. Seven leaf samples and one root sample were omitted from the dataset due to insufficient 16S or ITS reads (< 150). The resulting ASVs were curated using *lulu* v0.1.0 (Frøslev et al. 2017) with default settings. Fourteen contaminants were removed from the 16S dataset based on analysis of negative controls with *decontam* v1.10.0 (Davis et al. 2018), which led to the removal of 9 samples with < 150 reads. Initial taxonomic identification of 16S ASVs relied on SILVA v132 (Glöckner et al. 2017) and ITS were based on UNITE (Nilsson et al. 2019). We then removed singleton ASVs and any ASVs that were not identified as bacteria or fungi in the 16S and ITS data, respectively. To improve ASV identification and generate phylogenies, we used T-BAS v2.3 to place both 16S and ITS sequences on reference trees (Carbone et al. 2019). The resulting tables included 16,296 bacterial ASVs from 323 samples and 932 fungal ASVs from 332 samples (Appendix S1: Figure S1).

### Network Analyses

We analyzed community assembly of leaf and root compartments separately because they had very little overlap in previous analyses (Lee and Hawkes, 2021a), which we confirmed here for both fungi and bacteria (see Appendix S1: Methods/Results; Table S1; Figure S2).

In order to incorporate biotic interactions in community analyses, we first inferred cross-kingdom ecological association networks from which direct and diffuse interactions could be defined for the leaf and root communities (*SpiecEasi* v1.1.2 and *NetCoMi* v1.1.0; Kurtz et al. 2015; Peschel et al. 2021). Sparse inverse covariance estimation for ecological association inference (SPIEC-EASI) improves upon correlation methods, such as co-occurrence or joint species distribution models, without relying on pairwise interactions. Instead, SPIEC-EASI acknowledges the conditional states of all nodes such that the resulting inverse covariance matrix contains only values for conditionally dependent (direct) interactions, while the conditionally independent (indirect) interactions are near zero (Kurtz et al. 2015).

Prior to network construction, we filtered samples to only those present in both fungal and bacterial datasets and removed rare and low abundance taxa. Specifically, we 1) filtered leaf ASVs to include only those present in ≥10% of samples, 2) filtered root ASVs to only those present in either ≥10% of samples for fungi or ≥25% for bacteria, and 3) retained only those ASVs with >1% relative abundance. Filtering by prevalence and abundance resulted in 235 out of 1,002 bacterial ASVs and 114 out of 215 fungal ASVs in leaf communities and 388 out of 11,167 bacterial ASVs and 75 out of 730 fungal ASVs in root communities. The filtering did not remove any sites from the analysis, and overall microbial abundance patterns across sites were comparable before and after filtering. Networks were created for each site and filtered to include only high-confidence edges, where the edge-wise variability estimate was >0.8, indicating that the edge appeared in 80% of subsampled datasets. We performed a union of the site-level ASV networks and their putative interactions and inferred final networks for roots and leaves (Figure 1, Appendix S1: Table S2; *Cytoscape* v3.10.1, Shannon et al. 2003).

**Figure 1.**
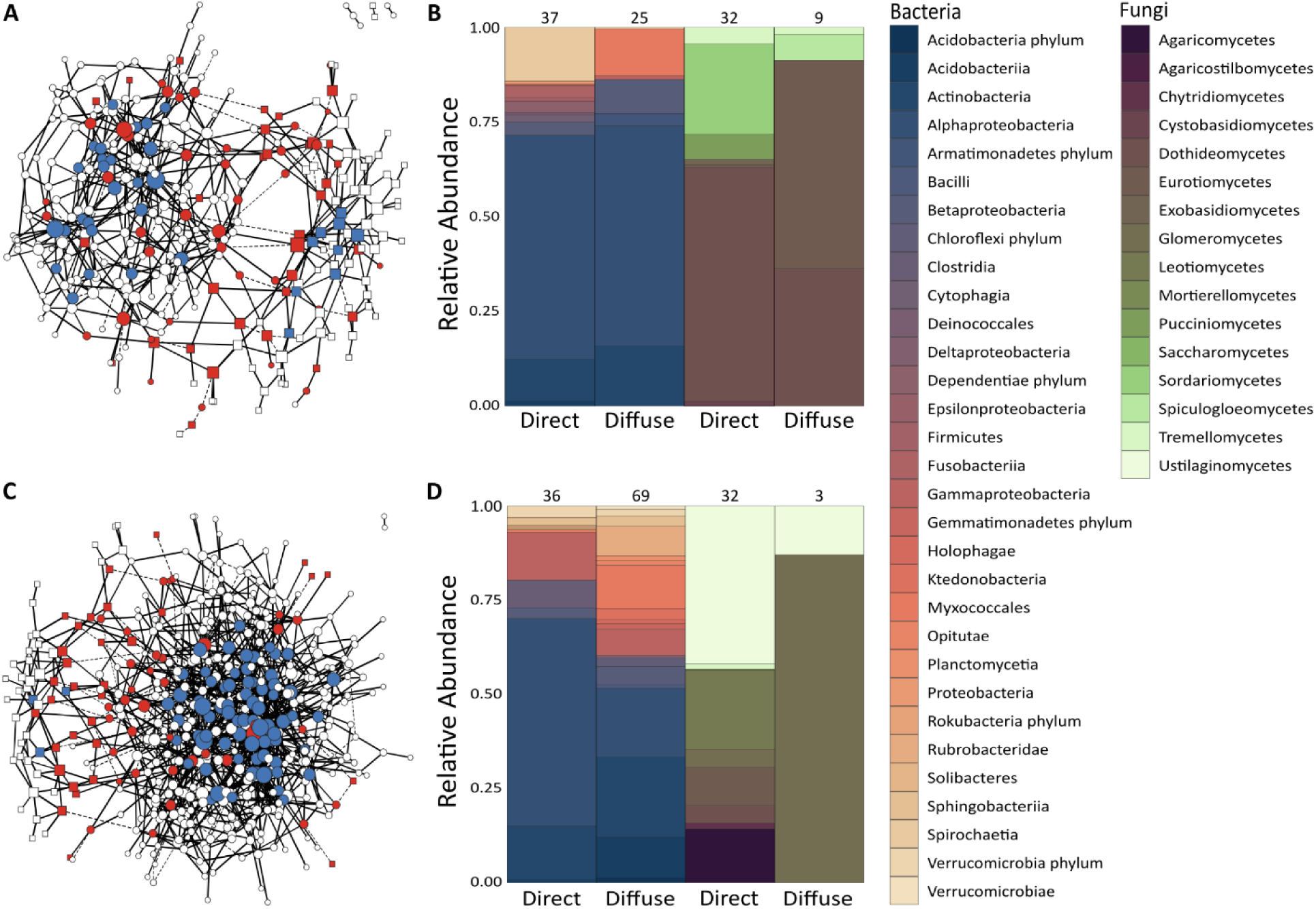
Ecological association networks for (A) leaves and (C) roots used to define cross-kingdom interactions. Direct interactors are red nodes, keystone hub taxa are blue nodes, and taxa with only within-kingdom interactions are white nodes. Edge line type indicates the direction of association (negative are dashed and positive are solid). The relative abundance of taxa defined in each interaction type are shown for (B) leaves and (D) roots, with the total number of taxa at the top of each bar and color representing taxonomic order.

We defined two categories of cross-kingdom interactions: direct and diffuse. Directly interacting taxa were defined by network edges connecting bacterial and fungal nodes (Appendix S1: Tables S3-4). Diffuse interactions were represented in two ways, 1) by keystone hub taxa (Appendix S1: Tables S5-6) and 2) by alpha diversity. Keystone hub taxa—defined as network nodes in the 75th percentile of degree and eigenvector centrality within each kingdom—represent those that are most influential to the network and may be necessary to community assembly without directly interacting with the other kingdom (Berry and Widder 2014). As an alternative to the keystone model, alpha diversity—defined as the richness of either bacteria or fungi in each sample—may better represent the set of all possible interactions between the kingdoms (Fiore-Donno et al. 2024). However, because alpha diversity never explained more than 1.5% of the variation in community composition, these results are only presented in the supplement (Appendix S1: Methods/Results, Figure S3). To better understand the bacterial and fungal taxa identified as direct or diffuse interactors, we tested for phylogenetic conservation of interaction type with Pagel’s λ (*geiger* v2.0.10, *ape* v5.6-2, *phytools* v1.0-1; Pennell et al. 2014; Paradis and Schliep 2019; Revell 2012). We also conducted literature searches to identify putative ecological roles of all direct and diffuse ASVs (Appendix S1: Table S7).

### Statistics

To address our hypotheses, we examined whether 1) interactions with bacteria (direct or diffuse) improved our ability to explain fungal community assembly and 2) whether interactions with fungi (direct or diffuse) improved our ability to explain bacterial community assembly. To do this, we used variance partitioning analysis (*vegan* v2.6-4; Oksanen et al. 2022) with either bacterial or fungal community composition as the dependent variable, represented as the Hellinger-transformed ASV count matrix (Peres-Neto et al. 2006). Independent predictors were matrices of environmental variables, spatial variables, and a set of biotic interactions (either direct, diffuse keystone hubs, or diffuse alpha diversity). Each variance partitioning analysis included the same environmental and spatial variables (Appendix S1: Table S8). Environmental variables included climate (MAP, MAT), plant size (height), microbial biomass, soil macronutrients (C, N, P, K), and soil properties (pH, texture, moisture). Spatial variables were stand area and sampling location (latitude, longitude). All environmental and spatial variables were standardized by scaling to zero mean and unit variance using the *decostand* function from the *vegan* R package (v2.6-4; Oksanen et al. 2022).

Each variance partitioning analysis was run four times: 1) without biotic interactions, then with biotic interactions represented by a secondary ASV matrix from the opposite kingdom including either 2) the directly interacting ASVs, 3) the diffusely interacting ASVs represented by keystone taxa, or 4) the diffusely interacting ASVs represented by alpha diversity (Appendix S1: Table S9). The sensitivity of these results to different network definitions is provided in Appendix S1: Tables S10-15. All testable fractions from the variance partitioning analyses were analyzed for significance with redundancy analysis (RDA) models and *P*-values were calculated based on a distribution of *R*^2^ values from permuted data (*vegan* v2.6-4; Oksanen et al., 2001). Shared variation fractions between environmental factors and biotic interactions were further investigated using permanova (*RRPP* v1.2.1; Collyer and Adams 2018) to determine environmental variables significantly associated (*P* < 0.05) with the ASVs. A subset of the significant environmental variables with the highest *R*^2^ values were further examined using Threshold Indicator Taxa Analysis (*TITAN2* v2.4.3; Baker and King 2010) to characterize how occurrence and relative abundance of each ASV shifts along the environmental variable ranges.

## RESULTS

### Identification of direct and diffuse interactions

*Direct interactions.—*Putative cross-kingdom direct interactions identified by ecological association networks included 69 taxa in leaves (37 bacteria and 32 fungi) and 68 taxa in roots (36 bacteria and 32 fungi), representing 22% and 15% of all taxa included in the networks, respectively (Figure 1). These directly interacting taxa were phylogenetically diverse, representing 26 bacterial orders and 19 fungal orders (Figure 1). In both the leaf and root networks, a similar number of direct edges were found with positive association estimates indicating co-occurrence between fungal and bacterial taxa (leaf: 24, root: 18) and negative associations indicating inverse relationships between taxa (leaf: 19, root: 20, respectively). The distribution of ASV abundances across sites indicated that both bacteria and fungi with direct cross-kingdom interactions ranged from rare (2-3 sites) to prevalent (8-14 sites) (Appendix S1: Figure S4). Most taxa involved in cross-kingdom direct interactions were only connected to one other ASV (78%), but 21% had multiple direct links (Appendix S1: Table S16). For example, in the leaf network, an Alpha-Proteobacteria ASV in the genus *Bosea*, which is known for plant growth promotion, shared edges with ASVs in three different fungal genera: two basidiomycetous yeasts (*Fibulobasidium*, *Bulleribasidium*) and one agaric fungus (*Chrysomycena*) (Appendix S1: Tables S3, S7). Similarly, in the root network, a bacterial ASV in the genus *Streptomyces*, also known for promoting plant growth, shared edges with an ascomycetous yeast (*Citeromyces* sp.) and a widespread basidiomycetous fungus (*Hydnum* sp.) (Appendix S1: Tables S3, S7).

*Diffuse interactions.—*Keystone taxa representing putative diffuse interactions included 25 bacteria and 9 fungi identified for the leaf community and 69 bacteria and 3 fungi for the root community (Figure 1). Keystone ASVs were generally prevalent, with bacterial keystone taxa found in 7 to 14 sites and fungal keystones found in 4 to 14 sites (Appendix S1: Figure S4). At the genus-level, the most frequently occurring keystone bacteria in leaves were Actinomycetes represented by three *Nocardiodes* ASVs that are potentially associated with plant growth promotion (Appendix S1: Tables S7, S16). The most common keystone bacteria in the root were ASVs in the Alpha-Proteobacteria (*Filomicrobium* sp.), Acidobacteria (*Koribacter* sp.), and Actinomycetes (*Mycobacterium* sp.; a putative plant growth promoter) (Appendix S1: Tables S7, S16). Of the 9 keystone fungal ASVs in the leaf network, 7 were Ascomycota (5 Dothideomycetes, 2 Eurotiomycetes) and 2 were Basidiomycota yeasts (Tremellomycetes, Spiculogloeomycetes). In roots, 2 of the 3 keystone fungi were Glomeromycetes—arbuscular mycorrhizas in the genus *Paraglomus*—and the third ASV was a basidiomycetous pathogen from the genus *Shivasia* (Ustilaginomycetes) (Appendix S1: Tables S7, S16).

*Diversity and phylogenetic relationships*.—As described above, both direct and diffuse interactors *within* each kingdom included phylogenetically diverse taxa with a range of putative ecological roles (Figure 1, Appendix S1: Tables S3-7). No phylogenetic signals were detected for interaction type (Appendix S1: Table S17), so further analyses did not account for phylogeny. This is supported in the observed composition patterns, where neither bacteria nor fungi had much taxonomic overlap below the level of order. For example, Dothideomycetes fungal ASVs were found across three out of four conditions (direct leaf, diffuse leaf, and direct root), but included 13 different genera among which *Sphaerellopsis* and *Cumuliphoma* (both potentially pathogenic) were dominant in leaves and *Poaceascoma* (potentially endophytic) was dominant in roots based on relative abundance.

### Contributions of cross-kingdom biotic interactions to community assembly

In the absence of biotic interactions, environmental variables explained 27.5% and 18.6% of fungal and 22.2% and 21.9% of bacterial leaf and root communities, respectively (Figure 2, Appendix S1: Table S9). When biotic interactions were considered, standalone environmental contributions to community composition declined by ∼15-20% in leaves and ∼8-10% in roots (Figure 2, Appendix S1: Table S9). In all analyses with and without biotic interactions, spatial variables and their intersections explained less than ∼10% of the variation.

**Figure 2.**
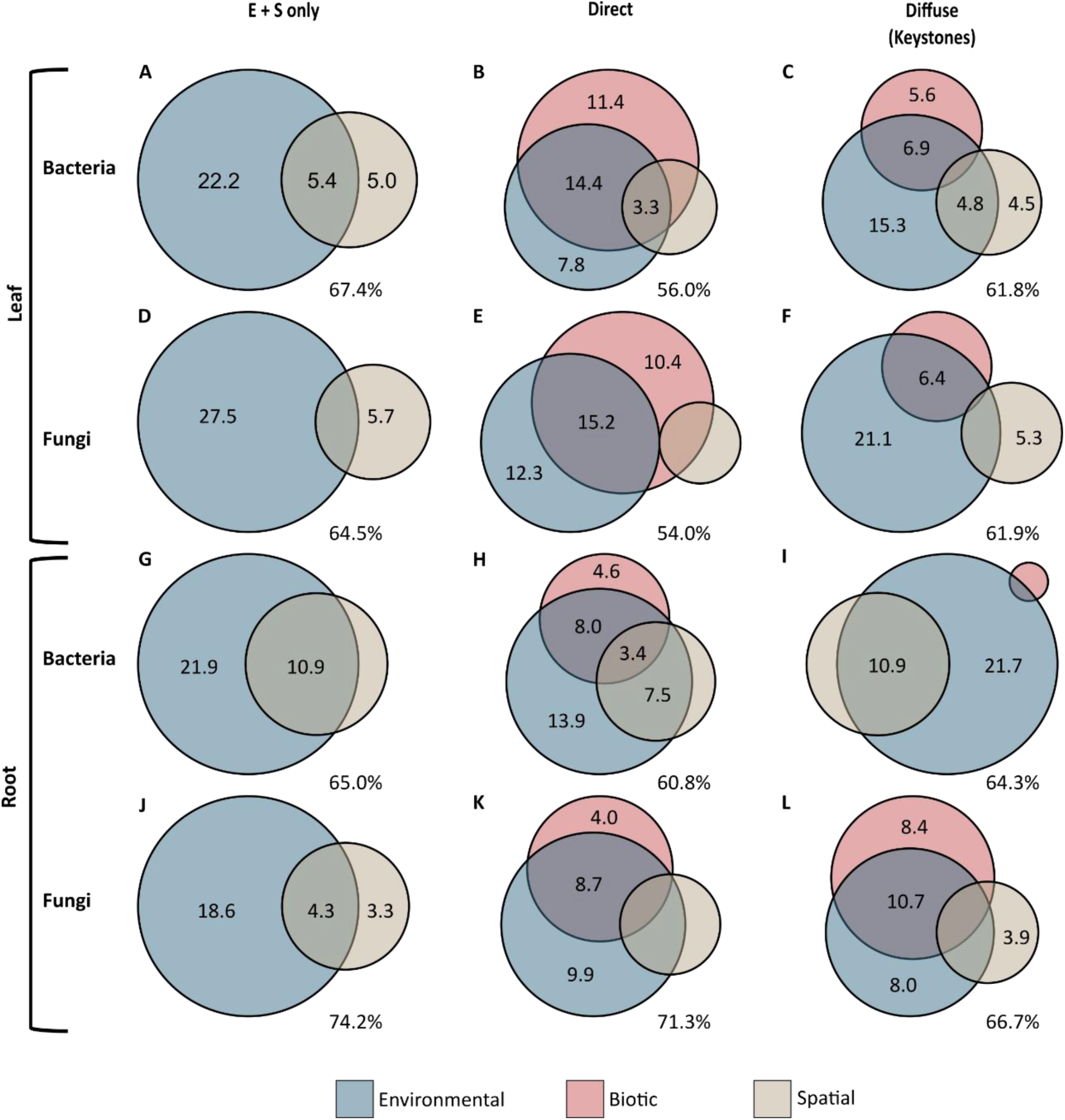
Variance partitioning of microbial community composition when including only environmental and spatial (E+S) factors, or additionally including direct or diffuse biotic interactions. Rows are the bacterial communities in the (A-C) leaf and (G-I) root or the fungal communities in the (D-F) leaf and (J-L) root, with each column indicating the subset of taxa used to represent the cross-kingdom biotic interactions. The size of each circle is relative to all other values in each set. Residuals are reported adjacent to each Venn diagram. Only values > 3% are displayed.

Putative direct cross-kingdom interactions were better predictors than diffuse interactions in most cases, explaining 2 to 6 times more variation in community composition of foliar fungi (10.4% direct vs. 2.6% diffuse), foliar bacteria (11.4% direct vs. 5.6% diffuse), and root bacteria (4.6% direct vs. 0.7% diffuse; Figure 2, Appendix S1: Table S9). The opposite pattern was observed only for root fungi, where diffuse interactions with bacteria explained more than twice as much of the variation in fungal community composition (8.4%) compared to direct interactions (4.0%).

The joint effects of environment and biotic interactions also helped to explain community patterns. These joint effects were dominant in leaves when considering direct interactions for both fungi (15.2%) and bacteria (14.4%) and in root fungal communities when considering either direct (8.7%) or diffuse (10.7%) cross-kingdom interactions (Appendix S1: Table S9). In all other cases, the joint effects of environment and cross-kingdom interactions explained less than environment alone (Appendix S1: Table S9).

The shared effects of environment and biotic interactions largely reflect correlations between interacting ASVs and environmental variation (Appendix S1: Table S18). In leaves, community-level TITAN analyses revealed that soil texture (% sand) structured bacterial and fungal assemblages across interaction types. Taxa that increased in abundance with soil sand content were most often found above ∼60% sand, whereas those taxa that decreased with soil sand content were most often found at ∼30% sand (Figure 3A-D). In roots, the strongest responses were in bacterial communities, with the highest summed z-scores and narrow peaks along soil pH and soil carbon gradients (Figure 3E-H). Root-associated bacterial direct interactors exhibited pH-associated increases, with the strongest change point at pH 5.2 (Figure 3E). Root-associated direct and diffuse bacteria primarily responded negatively to soil carbon content, with peak compositional change at ∼1.0% carbon (Figure 3F-G). Root fungal ASVs characterized as direct interactors also responded to soil nitrogen, with a change point of ∼0.04% for taxa responding negatively and ∼0.2% for taxa responding positively (Figure 3H). Responses of individual taxa are provided in the supplement (Appendix S1: Figure S6, Tables S19-20).

**Figure 3.**
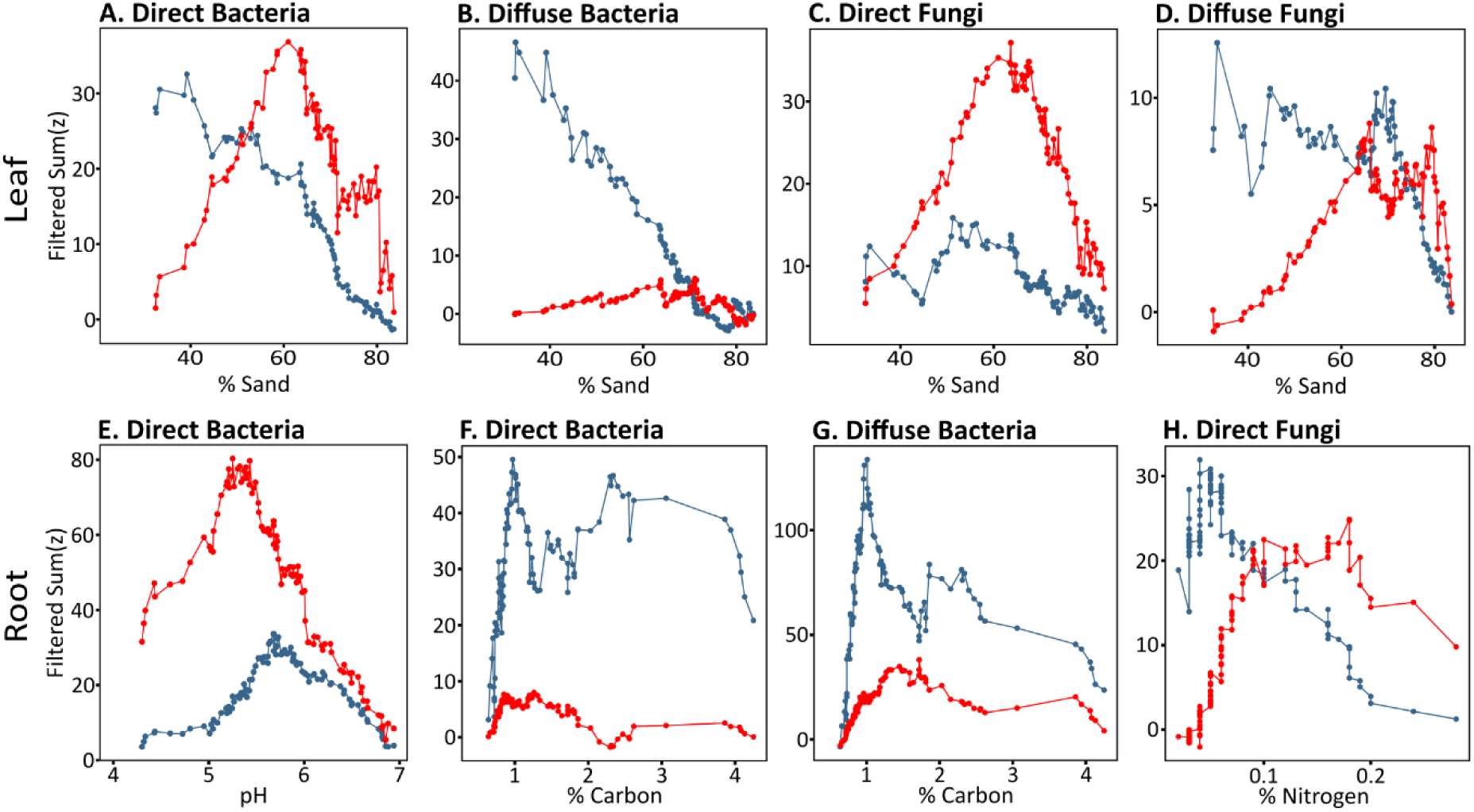
Community-level TITAN summed z-score responses of (A-D) leaf and (E-H) root-associated microbial interactors along selected environmental gradients. Positive (red) and negative (blue) curves represent taxa increasing or decreasing in abundance along each gradient, respectively. Peaks in summed z-scores indicate community change points where coordinated shifts among taxa are strongest. Full TITAN2 results are reported in Appendix S1: Tables S19-S20.

## DISCUSSION

Cross-kingdom interactions between bacteria and fungi represented almost a quarter of all links in their networks and were an important driver of microbial community composition similar in importance to environmental and spatial predictors. Their contributions increased even further when considering the joint effects of environment and biotic interactions, which highlights the complexity of factors influencing microbial community composition and the need to expand the classic approach to describing community assembly. Our findings add to the growing pool of research studies that have found biotic interactions to be important in explaining community structure (Fiore-Donno et al. 2024; Li et al. 2023), while offering a method for better defining those biotic interactions in ecological terms.

Our original hypothesis, that cross-kingdom direct interactions would be more important than diffuse interactions in lower diversity leaf environments, was largely supported. Microorganisms tend to cluster in microhabitats on leaves, enabling direct competition for space and resources or other antagonistic or mutualistic interactions (Chaudhry et al. 2020). For example, several studies have shown the importance of bacteria-bacteria interactions in *Arabidopsis* leaves—these interactions tend to be negative, supported by antimicrobial compound production and shifts in community composition that can lead to damaged plant tissue (Chen et al. 2020; Schäfer et al. 2022). Other studies have identified cross-kingdom interactions in plants where bacteria may have an antagonistic effect on fungal pathogens. In one case, Kamensky et al. (2003) demonstrated that a gram-negative bacterial strain was able to protect the plant host from multiple necrotrophic fungi and identified production of several anti-fungal compounds as the most likely mechanism. In the current study, the reported ecological roles of the taxa involved in putative direct interactions suggest the potential for either competition (negative edges) or facilitation (positive edges) between bacteria and plant pathogenic fungi. Putative direct interactions identified from ecological association networks may point to a key subset of ecologically important taxa that can be prioritized for further study.

In roots, we originally expected cross-kingdom diffuse interactions to be better predictors than direct interactions given the typically high diversity of root microbial communities. However, this was only the case for root fungi and their diffuse interactions with bacteria. This outcome aligns with their observed alpha diversities—bacterial richness in roots was six times higher than in leaves, but fungal richness was comparatively lower and did not change between the plant compartments. The higher bacterial richness we found in roots may increase the potential for diffuse interactions or modifications to the environment. Supporting this, selective reductions in soil microbial diversity result in expansions of previously rare taxa and modifications of specific biogeochemical processes, potentially altering local environmental conditions (Romdhane et al. 2022). Alternatively, the larger number of bacteria may have allowed for more robust identification of keystone taxa compared to the fungi.

Keystone taxa in microbial networks tend to be important for community structure, ecosystem functioning, and interactions with abiotic and biotic factors (Yang et al. 2021; Agler et al. 2016). We found keystone taxa to be a good representation of diffuse interactions, likely because defining keystone ASVs as network nodes that were highly connected (degree) and had a strong influence (eigenvector centrality) partly captured their role in determining overall community structure (Yang et al. 2021). In support of our approach, Berry and Widder (2014) demonstrated that high degree, closeness centrality, and transitivity corresponded to “keystoneness.” In contrast, we found that alpha diversity had poor explanatory power for community assembly, perhaps because using overall species richness to represent diffuse interactions can dilute the effects of keystone taxa. This differs from other studies that reported modest effects of biotic interactions defined as species richness on community assembly. For example, Fiore-Donno et. al. (2024) observed that species richness explained composition in some seasons, indicating that biotic interactions may be linked to environmental conditions, as observed in our study.

Up to 15% of the variation in microbial community structure was present in the joint effects of environment and biotic interactions, likely caused by both direct and diffuse ASVs that tracked specific environmental conditions. Although there were no universal patterns across taxa, there were clear trends that suggest potential ecological processes. For example, 70-75% of leaf ASVs were either positively (direct interactors) or negatively (diffuse interactors) associated with soil sand content, suggesting that soil texture may influence source pools, local dispersal, or recruitment to the plant host. Sand content affects multiple soil properties, including nutrient and water availability, for which it may serve as a proxy, but we also found some taxa that were correlated with soil nutrients. In roots, diffusely and directly interacting bacteria and directly interacting fungi exhibited large declines in relative abundance at distinct thresholds of carbon or nitrogen, consistent with these serving as key ecological filters. In most cases, environmental factors are likely driving ASV distributions, which can affect the structure of microbial co-occurrence networks by increasing or reducing potential interactions (Lekberg et al. 2021). Alternatively, biotic interactions can lead to changes in environmental factors, such as altering available nutrients via the formation and degradation of soil organic matter, that are important to both the plant host and microbial community (Sokol et al. 2022). Understanding the relationships between abiotic factors and microbial species distributions can offer insight into what drives the correlations underlying joint effects.

Inferring biotic interactions from ecological association networks based on amplicon sequencing of rDNA necessarily means that these interactions are putative. Although this approach is not purely correlational, it relies on distributional co-occurrence patterns captured at a single time point rather than explicit tests of interaction. Similarly, we identified putative ecological roles of taxa from trait databases and cases reported in the literature, which we note are particularly problematic at the genus level when both conspecifics and congeners can exhibit multiple or context-dependent trophic strategies. The identification of putative interactions and ecological roles might be improved by using methods that characterize the community over time to identify how consistent these putative interactions might be (Fiore-Donno et al. 2024; Yuan et al. 2021), or by employing multi-omics networks that model metabolic interactions (Muller et al. 2018). Alternatively, stable isotope probing could identify active taxa that use the same substrates and likely compete (Prosser et al. 2006), taxa turnover using labeled oxygen (Schwartz et al. 2016), or taxa that are part of a trophic web (Lueders et al., 2004).

Nevertheless, our results were robust to the changes in network parameters and keystone threshold. Variance partitioning results at the different edge-wise variability estimates were all within a 10% range of total variation explained. The largest difference observed with increasing stringency of the edge-wise variability estimate was in network connectedness, evidenced by the dramatic decrease in the number of edges especially those that were cross-kingdom. In the current study, the majority of keystone taxa were retained across different edge-wise variability estimates, though there were unique keystones identified in each due to the change in nodes and edges maintained at increasing stringency. The minimal sensitivity of overall outcomes to these changes suggests that putative direct and diffuse interactions identified from networks are largely internally consistent and can improve our assessment of community drivers.

The incorporation of biotic interactions into investigating microbial community assembly is a relatively recent and growing trend that is supported by our results. We took this a step further by defining different types of interactions—direct and diffuse—that proved to be important sources of variation in microbial community composition aligned with expectations from ecological theory. However, it is important to note that there was still 54-74% residual variation that was unexplained by our models. These efforts are hampered by reliance on putative interactions and difficulty in identifying true interactions in complex microbial communities. To partly overcome this, others have successfully used both taxonomic and functional characteristics of taxa to better capture interactions in well-characterized systems (Li et al., 2023). Moving forward, community assembly models can be improved by a greater understanding of how we can better estimate and represent cross-kingdom biotic interactions, which we expect will also facilitate better implementation of applied development of synthetic microbial communities.

## ACKNOWLEDGEMENTS

For assistance with lab and field work, we thank N. Yang, B. Whitaker, I. McBryde, J. Czel, E. Hardy, and M. Alvarez. Site access was provided by NCSU, North Carolina Department of Agriculture & Consumer Services, BASF Corp., Weyerhaeuser Co., E. Deal, and C. Wilson. The manuscript was improved by comments from M. Cubeta. This research was supported by a DOE Genomic Sciences Program SFA (SCW1039) subaward, by the Research Capacity Fund (HATCH), project award no. 7005451, from the U.S. Department of Agriculture’s National Institute of Food and Agriculture, and by startup funding from North Carolina State University to CVH.

## AUTHOR CONTRIBUTIONS

The experiment was designed by CVH. MRL led field and lab work and carried out the bioinformatics of amplicon sequencing data. RAH analyzed the data. RAH and CVH wrote the manuscript. All authors edited the manuscript.

