## Supplemental Material for "Direct and diffuse cross-kingdom interactions in plant microbiome assembly"


| Item | Description | Page |
|---|---|---|
| Methods/Results | | |
| | Alpha diversity | 2 |
| | Diffuse interactions – alpha diversity | 2 |
| | Network sensitivity analysis | 2 |
| Figures | | |
| Fig. S1 | Sample rarefaction curves | 4 |
| Fig. S2 | Sample ordinations | 5 |
| Fig. S3 | Alpha diversity | 6 |
| Fig. S4 | Occupancy-abundance plots | 7 |
| Fig. S5 | Network sensitivity | 8 |
| Fig. S6 | Environmental gradient analysis (TITAN2) results | 9 |
| Tables | | |
| Table S1 | PERMANOVA results of beta diversity on plant compartment | 10 |
| Table S2 | Network statistics across parameters to investigate sensitivity | 11 |
| Table S3 | Bacterial taxa - putative direct cross-kingdom interactions | 12 |
| Table S4 | Fungal taxa - putative direct cross-kingdom interactions | 15 |
| Table S5 | Bacterial taxa - putative diffuse (keystone) interactions | 17 |
| Table S6 | Fungal taxa - putative diffuse (keystone) interactions | 21 |
| Table S7 | Putative ecological roles for taxa identified as interactors | 22 |
| Table S8 | Independent variables for variance partitioning analysis (VPA) | 27 |
| Table S9 | Main variance partitioning analysis results | 28 |
| Table S10 | VPA sensitivity results - leaf, direct interactions | 29 |
| Table S11 | VPA sensitivity results - leaf, diffuse (keystone) interactions | 30 |
| Table S12 | VPA sensitivity results - leaf, diffuse (alpha) interactions | 31 |
| Table S13 | VPA sensitivity results - root, direct interactions | 32 |
| Table S14 | VPA sensitivity results - root, diffuse (keystone) interactions | 33 |
| Table S15 | VPA sensitivity results - root, diffuse (alpha) interactions | 34 |
| Table S16 | Frequency tables of direct and diffuse interactors | 35 |
| Table S17 | Pagel's lambda phylogenetic signal results | 36 |
| Table S18 | PERMANOVA results of interactors on environmental variables | 37 |
| Table S19 | Significant ASVs from TITAN2 analysis - leaf | 39 |
| Table S20 | Significant ASVs from TITAN2 analysis - root | 43 |
| References | | 47 |

### Supplemental Methods/Results

#### Diversity

We compared the alpha diversity of leaf and root fungal and bacterial communities using observed richness (Figure S3). Observed richness of bacteria ranged from 93.58 ± 4.55 in leaves to 570.15 ± 27.36 in roots (Figure S3). In contrast, fungal richness was lower than bacteria and was similar between leaves (38.75 ± 0.99) and roots (34.75 ± 1.36, Figure S3). Linear model evaluation with randomized residuals in a permutation procedure showed that both bacterial and fungal community alpha diversity significantly differed by compartment ($P < 0.05$, Table S1; R, rrpp v1.2.1; Collyer and Adams 2018).

Additionally, we evaluated the similarity of the leaf and root microbial communities by calculating the Euclidean distance on the centered log-ratio transformed bacterial and fungal ASV matrices to obtain Aitchison's distance, which is appropriate for compositional data (Gloor et al. 2017).We then tested how plant compartment affected fungal and bacterial community composition using residual randomized permutation procedures (RRPP v.1.2.1; Collyer and Adams 2018) and found that the leaf and root microbiota had minimal overlap ($P = 0.001$, Table S1; Figures S2).

#### Diffuse Interactions – Alpha Diversity

As an alternative metric for overall diffuse interactions, we used observed richness to represent alpha diversity. When used to represent diffuse biotic interactions in variance partitioning analyses, alpha diversity of the other kingdom explained only 0-1.5% of variation in community composition (Table S10 and S15).

#### Sensitivity Analysis

We looked at the sensitivity of key SPIEC-EASI parameters and keystone definitions to determine the robustness of our criteria decisions (Table S2, Figure S5) and how these definitions influenced variance partitioning analysis results (Tables S9-15). The edge-wise variability indicates the proportion of subsampled network matrices an edge appears in, thus allowing for filtering out edges with low confidence and reducing noise. We chose an edge-wise variability estimate of 0.8 as recommended by the creators, and we compared this to the results of no threshold, as well as two more stringent thresholds, 0.9, and 0.95. To define keystone nodes representing diffuse interactions, we used the 75$^{th}$, 80$^{th}$, or 90$^{th}$ percentile of both degree and eigenvector centrality within each kingdom, which represents increasing connectedness and influence (Berry and Widder 2014). The keystone criteria range was selected to balance identification of nodes highest in those metrics relative to the number and range of values in each group to include both bacteria and fungi at each threshold. Common practice involves using a method to determine the cutoff value, such as fitting a log-normal distribution or above an empirical percentile (Agler et al. 2016; Peschel et al. 2021).

Increasing the edge-wise variability estimate did not have a strong effect on the number of nodes retained in the networks, which declined by only 3-9% between 0 and 0.8 (Figure S5A). However, increasing the edge-wise variability cutoff from 0 to 0.8 did dramatically reduce the total number of edges (91-93%) and direct cross-kingdom edges (98-99%), but minimal reductions were seen with further increasing stringency (< 76%). This analysis demonstrates why a threshold is needed to remove poorly supported edges that are unlikely to be indicative of true interactions (Figure S5A).

At each keystone inclusion threshold, many keystone taxa (29-100%) were retained across different edge-wise variability estimates, though there were unique keystones identified in each due to the change in nodes and edges included at increasing stringency (Figure S5B). Within a network, increasing the threshold led to 11-64% fewer keystones (Figure S5B). In the full networks (i.e., no edge-wise variability estimate restriction), identifying keystone nodes with our definition was near impossible since 99% of the taxa had a direct cross-kingdom edge.

Based on these analyses, we chose an edge-wise variability estimate of 0.8 to reduce noise without being overly conservative. We also chose the 75$^{th}$ percentile for keystones because this retained at least one taxon from each kingdom at each level while still being representative of the most connected and influential taxa in the networks.

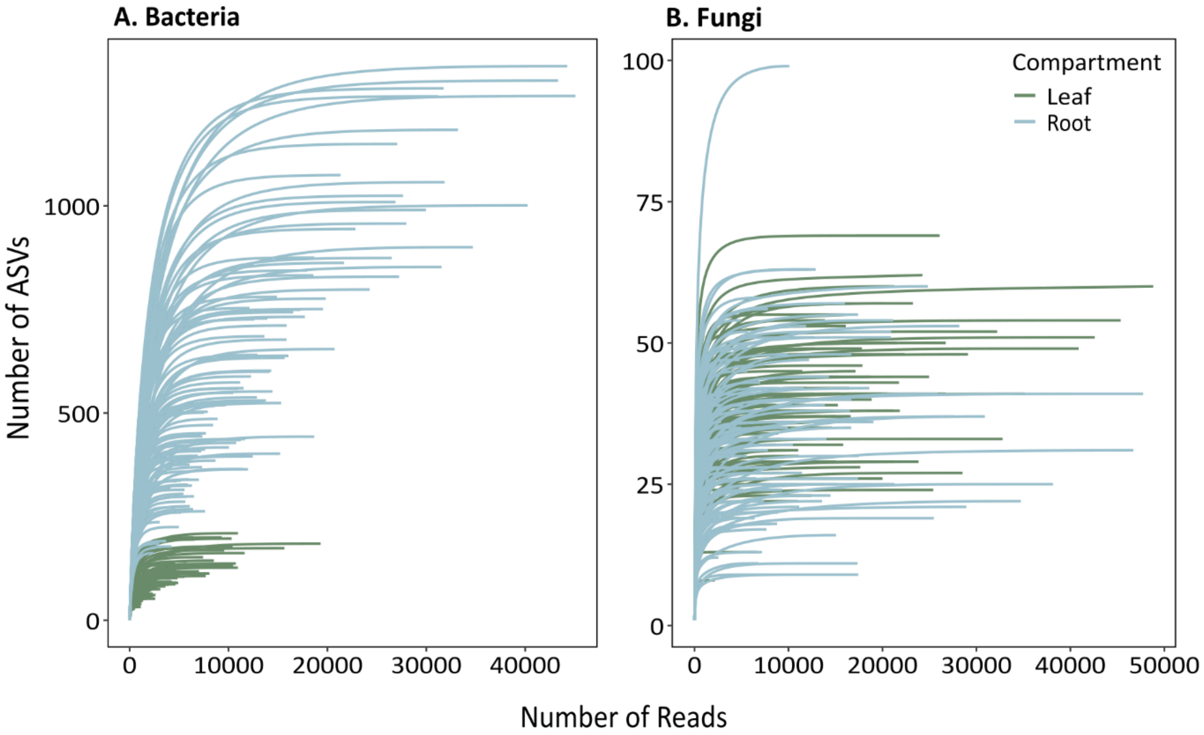


**Figure S1**. Plot of rarefaction curves for (A) bacteria and (B) fungi samples, showing the relationship between the number of reads per sample and cumulative number of ASVs detected. Samples are color coded by type (leaf, root, soil).

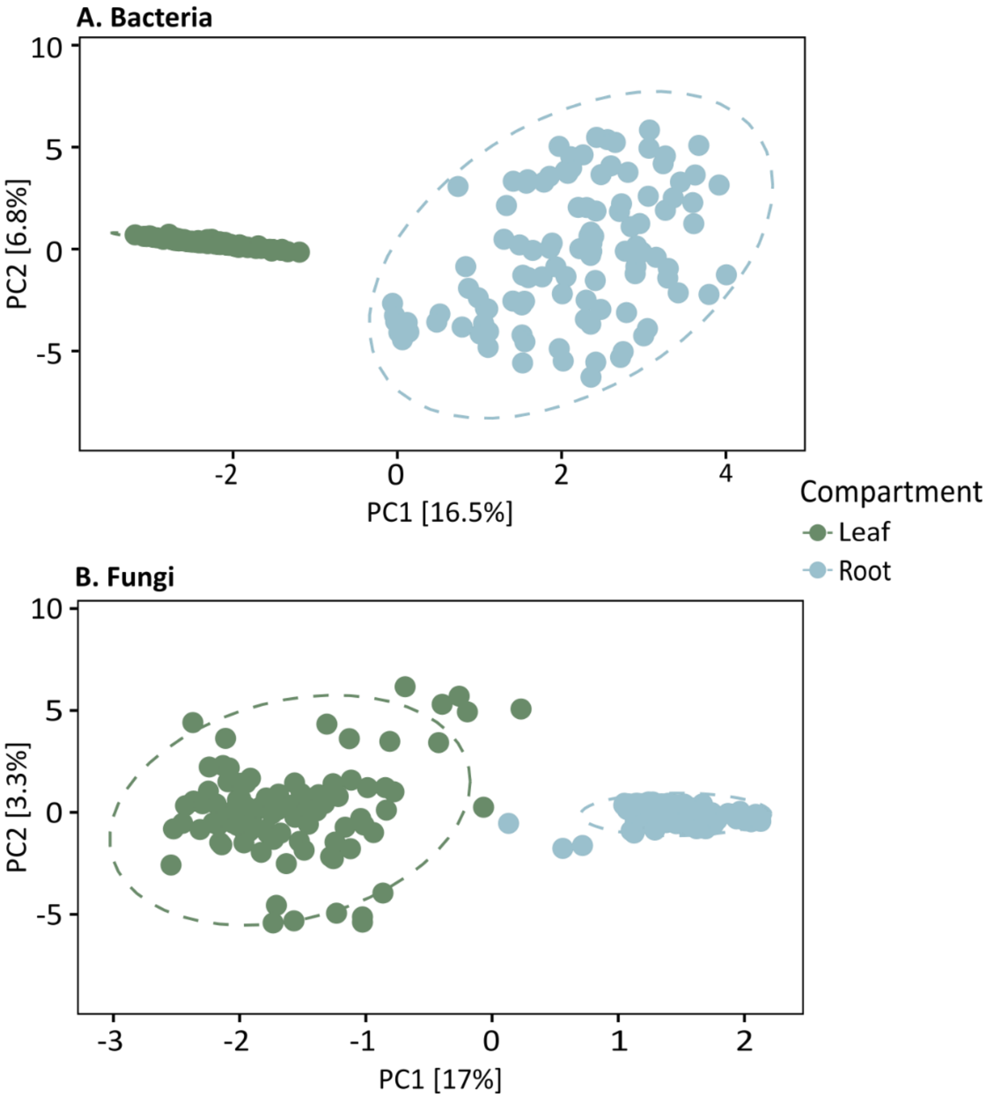


**Figure S2**. Results of redundancy analysis of CLR-transformed ASV matrices for (A) bacteria and (B) fungi. Samples clustered by compartment (leaf vs. root). For both bacteria and fungi, beta diversity was significantly different between compartments based on permANOVA ($P < 0.001$, Table S1).

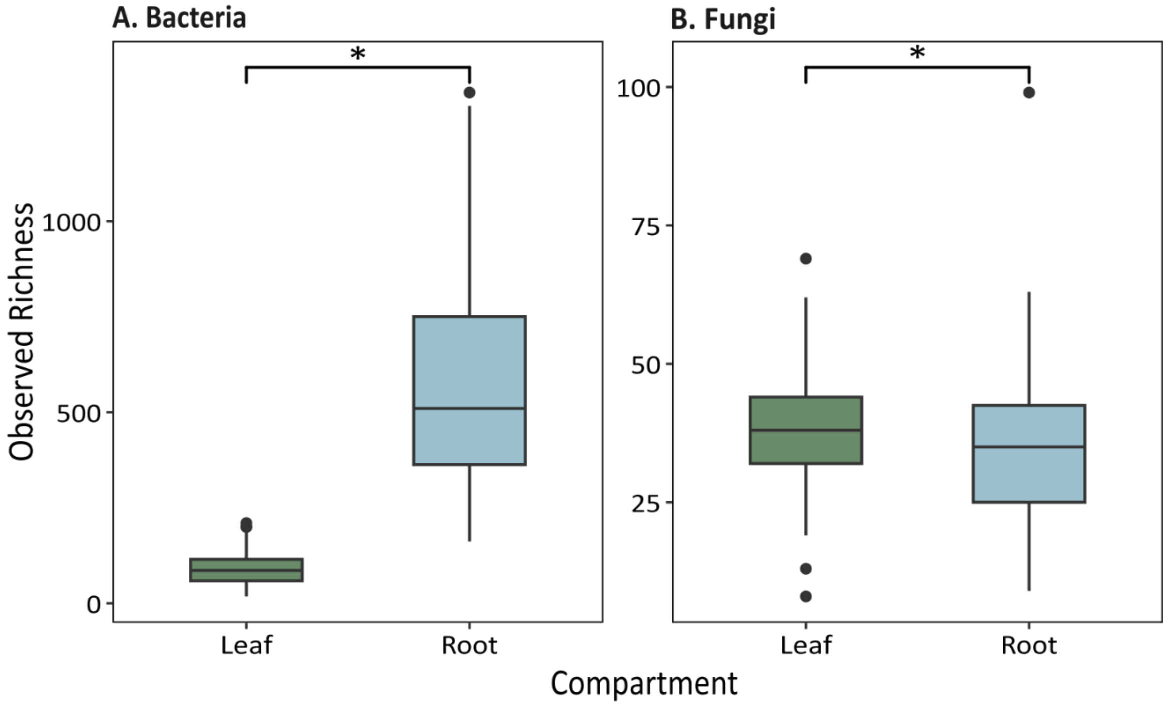


**Figure S3**. Alpha diversity of (A) bacteria and (B) fungi across leaf and root compartments. Box plots show median observed richness (horizontal line), interquartile ranges (box), and outliers (points). Asterisks indicate significant differences between compartments, alpha diversity of bacteria is lower in leaves than in roots (P = 0.001) while the opposite is true for fungi (P = 0.03, Table S1). Note the difference in magnitude on y-axis scales between panels.

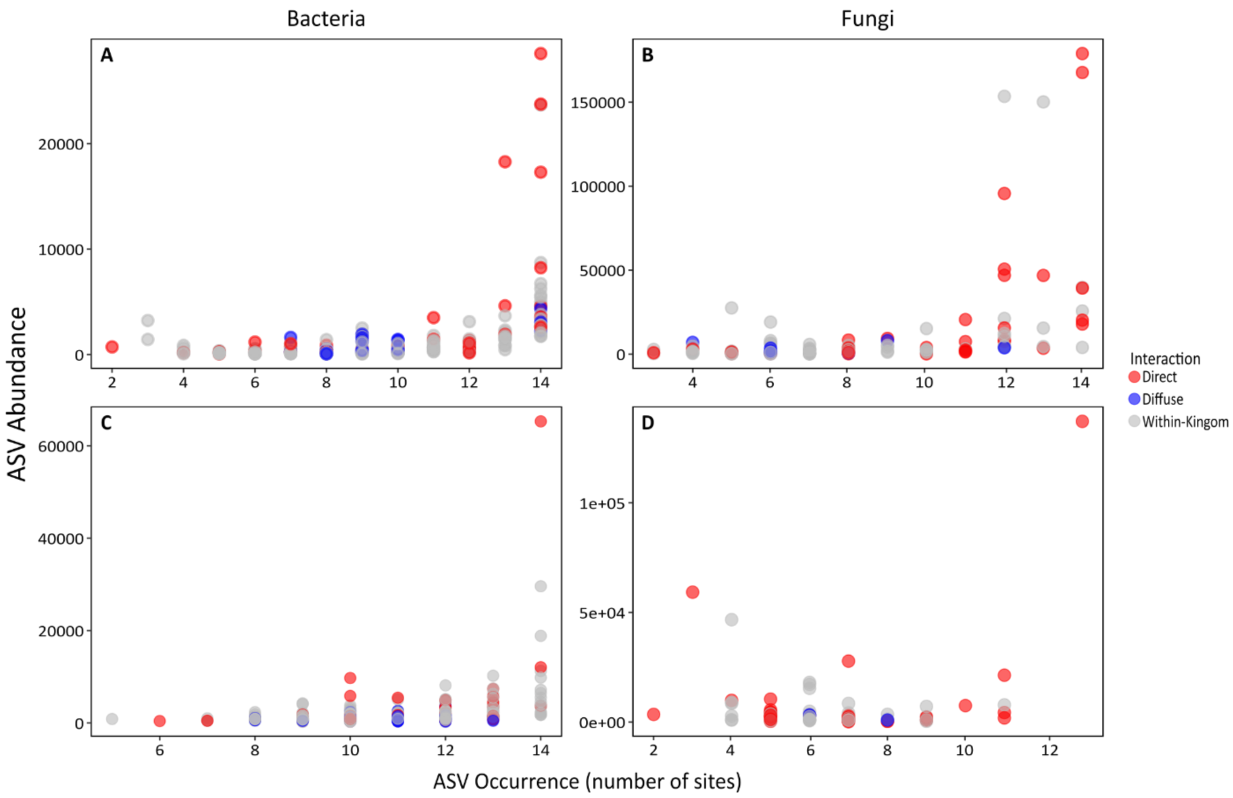


**Figure S4**. Occupancy-abundance plots of bacteria (A, C) and fungi (B, D) in leaves (A, B) and roots (C, D). Distribution of ASV frequency was similar across interaction types.

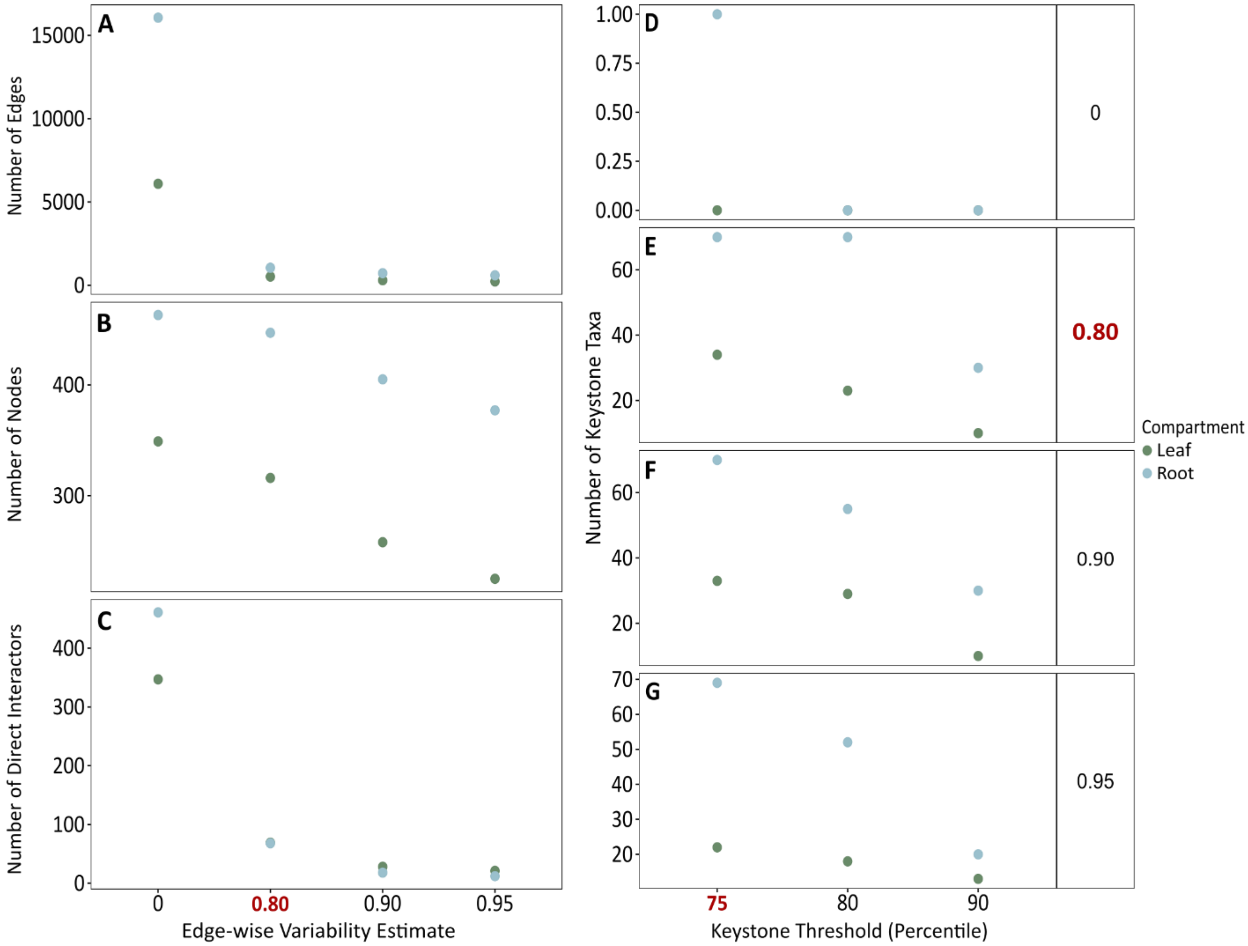


**Figure S5**. Sensitivity of network parameters to the (A-C) edge-wise variability estimate and (D-G) keystone inclusion criteria for each edge-wise variability threshold. Network conditions for the final analysis (edge-wise variability = 0.80, keystone threshold of 75$^{th}$ percentile) were chosen to balance confidence of included edges and keystones while still maintaining feasible networks and are indicated by bolded red text.

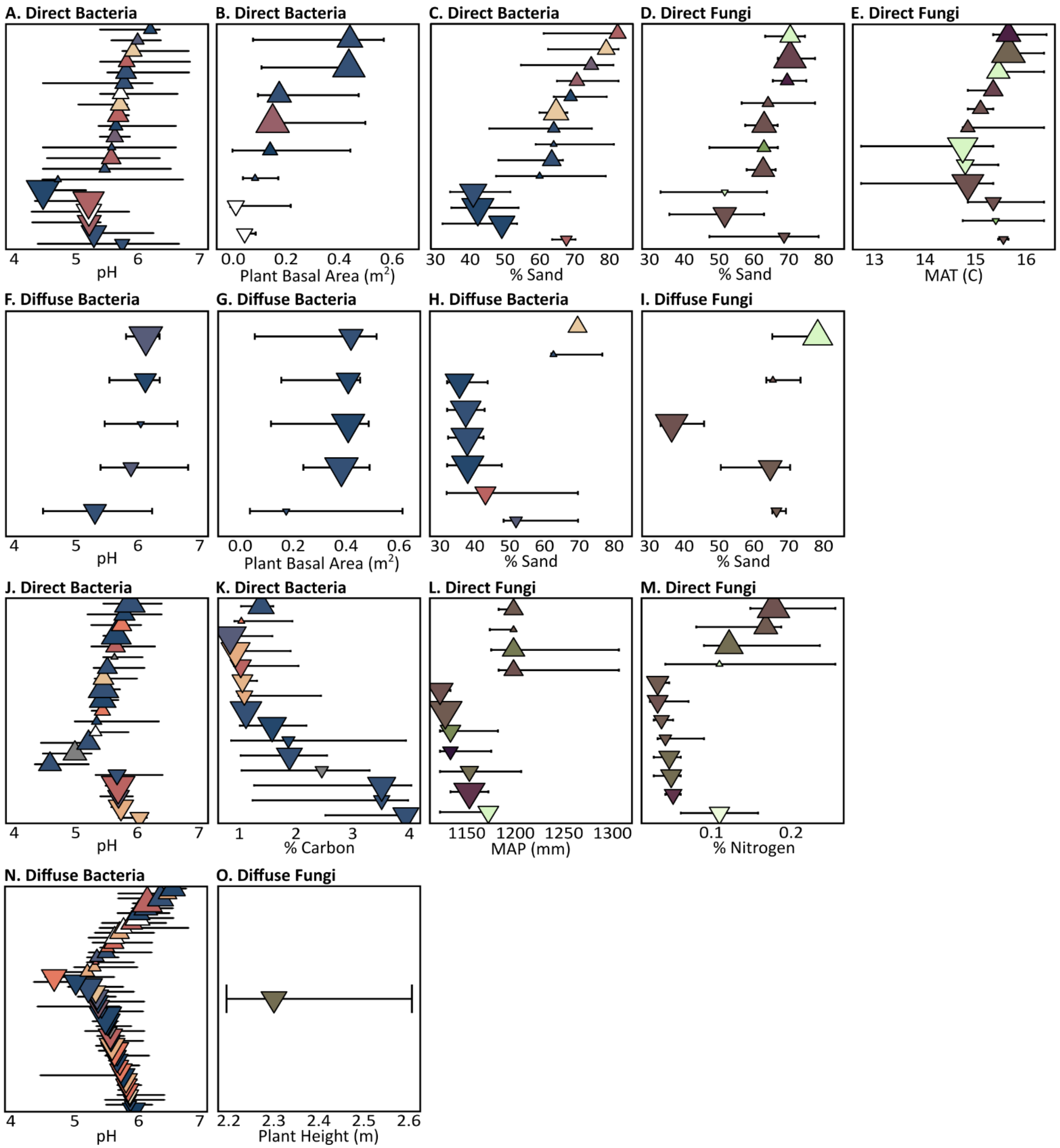


**Figure S6**. Environmental gradient analysis (TITAN2) results for (A-I) leaf or (J-O) root ASVs that significantly tracked selected environmental variables. Triangle symbols are the median change points of the ASV probability density function across the environmental gradient. Symbol size represents the strength of the median z-score, symbol direction indicates increases (up triangle) or decreases (down triangle) in relative abundance along the gradient, and symbol color represents taxonomic order as in Fig. 1. The horizontal lines around each symbol show the 5-95% quantiles of bootstrapped change point distribution.

**Table S1.** Results of permANOVA tests of microbial community (A-B) beta and (C-D) alpha diversity on plant compartments (leaf and root)

**A. Bacterial beta diversity by plant compartment**

| | df | SS | MS | $R^2$ | *F* | *Z* | *P* |
|---|---|---|---|---|---|---|---|
| Compartment | 1 | 201816 | 201816 | 0.1499 | 36.679 | 2.900 | 0.001 |
| Residuals | 208 | 1144450 | 5502 | 0.8501 | | | |
| Total | 209 | 1346265 | | | | | |

**B. Fungal beta diversity by plant compartment**

| | df | SS | MS | $R^2$ | *F* | *Z* | *P* |
|---|---|---|---|---|---|---|---|
| Compartment | 1 | 44395 | 44395 | 0.1568 | 38.679 | 3.404 | 0.001 |
| Residuals | 208 | 238737 | 1148 | 0.8432 | | | |
| Total | 209 | 283132 | | | | | |

**C. Bacterial alpha diversity by plant compartment**

| | df | SS | MS | $R^2$ | *F* | *Z* | *P* |
|---|---|---|---|---|---|---|---|
| Compartment | 1 | 12322071 | 12322071 | 0.5687 | 278.180 | 7.392 | 0.001 |
| Residuals | 211 | 9346286 | 44295 | 0.4313 | | | |
| Total | 212 | 21668357 | | | | | |

**D. Fungal alpha diversity by plant compartment**

| | df | SS | MS | $R^2$ | *F* | *Z* | *P* |
|---|---|---|---|---|---|---|---|
| Compartment | 1 | 756 | 756 | 0.0218 | 4.850 | 1.771 | 0.029 |
| Residuals | 218 | 33977 | 156 | 0.9782 | | | |
| Total | 219 | 34733 | | | | | |

**Table S2.** Statistics for each network from changing edge-wise variability estimate inclusion criteria and keystone percentile cut-offs to evaluate the sensitivity of results to changing thresholds. Value in the parentheses represents the number of taxa excluded from being keystones due to being involved in a direct cross-kingdom interaction (defined a fungal and bacterial node connected by an edge).

| | Leaf | | | | Root | | | |
|---|---|---|---|---|---|---|---|---|
| **Edge-wise Variability Estimate Cut-off** | **0.95** | **0.01** | **0.80** | **None** | **0.95** | **0.90** | **0.80** | **None** |
| Number nodes | 225 | 258 | 316 | 349 | 377 | 405 | 447 | 463 |
| Number edges | 230 | 301 | 522 | 6092 | 597 | 723 | 1051 | 16064 |
| Average number neighbors | 2.236 | 2.430 | 3.353 | 34.911 | 3.283 | 3.640 | 4.716 | 69.391 |
| Network diameter | 22 | 24 | 12 | 3 | 19 | 15 | 10 | 3 |
| Network radius | 11 | 12 | 7 | 2 | 10 | 8 | 6 | 2 |
| Characteristic path length | 9.827 | 7.963 | 5.253 | 1.978 | 6.307 | 5.590 | 4.392 | 1.852 |
| Clustering coefficient | 0.035 | 0.025 | 0.036 | 0.149 | 0.154 | 0.138 | 0.104 | 0.178 |
| Network density | 0.013 | 0.010 | 0.011 | 0.100 | 0.009 | 0.009 | 0.011 | 0.150 |
| Network heterogeneity | 0.581 | 0.577 | 0.552 | 0.405 | 0.653 | 0.665 | 0.614 | 0.253 |
| Network centralization | 0.027 | 0.024 | 0.028 | 0.099 | 0.025 | 0.024 | 0.026 | 0.095 |
| Connected components | 17 | 9 | 4 | 1 | 10 | 6 | 2 | 1 |
| Number of direct interactors | 21 | 28 | 69 | 347 | 12 | 18 | 68 | 461 |
| Number of direct bacteria | 11 | 15 | 37 | 233 | 6 | 9 | 36 | 386 |
| Number of direct fungi | 10 | 13 | 32 | 114 | 6 | 9 | 32 | 75 |
| % Taxa that interact cross-kingdom | 9.3 | 10.9 | 21.8 | 99.4 | 3.2 | 4.4 | 15.2 | 99.6 |
| Number of direct edges | 11 | 15 | 43 | 2062 | 6 | 9 | 38 | 2936 |
| % Edges that interact cross-kingdom | 4.8 | 5.0 | 8.2 | 33.8 | 1.0 | 1.2 | 3.6 | 18.3 |
| Number fungi | 86 | 91 | 104 | 114 | 59 | 63 | 71 | 75 |
| Number bacteria | 139 | 167 | 212 | 235 | 318 | 342 | 376 | 388 |
| Number keystone 90th | 13 (3) | 10 (3) | 10 (6) | 0 (31) | 20 (2) | 30 (4) | 23 (4) | 0 (41) |
| Keystone 90th - bacteria | 8 (1) | 6 (1) | 10 (1) | 0 (19) | 19 (0) | 29 (1) | 22 (1) | 0 (34) |
| Keystone 90th - fungi | 5 (2) | 4 (2) | 0 (5) | 0 (12) | 1 (2) | 1 (3) | 1 (3) | 0 (7) |
| Number keystone 80th | 18 (4) | 29 (7) | 23 (12) | 0 (62) | 52 (4) | 55 (5) | 64 (12) | 0 (81) |
| Keystone 80th - bacteria | 10 (2) | 19 (3) | 21 (5) | 0 (42) | 46 (2) | 52 (2) | 62 (4) | 0 (68) |
| Keystone 80th - fungi | 8 (2) | 10 (4) | 2 (7) | 0 (20) | 6 (2) | 3 (3) | 2 (8) | 0 (13) |
| Number keystone 75th | 22 (4) | 33 (8) | 34 (17) | 0 (84) | 69 (5) | 70 (5) | 72 (14) | 1 (102) |
| Keystone 75th - bacteria | 12 (2) | 21 (3) | 25 (6) | 0 (57) | 63 (2) | 63 (2) | 69 (6) | 1 (84) |
| Keystone 75th - fungi | 10 (2) | 13 (5) | 9 (11) | 0 (27) | 6 (3) | 7 (3) | 3 (8) | 0 (18) |

**Table S3.** Bacterial taxa involved in putative direct cross-kingdom interactions identified in the network using an edge-wise variability estimate of 0.8. Assignments result from placing unknown sequences on T-BAS phylogenetic reference tree.

| Compartment | ASV | Phylum | Class | Order | Family | Genus | Connections (fungal) |
|---|---|---|---|---|---|---|---|
| Leaf | ASV_1145 | Deinococcus-Thermus | Deinococcales | Deinococcales | Deinococcaceae | Deinococcus | ASV_138 |
| Leaf | ASV_1151 | Bacteroidota | Cytophagia | Cytophagales | Hymenobacteraceae | Siccationidurans | ASV_856 |
| Leaf | ASV_11551 | Proteobacteria | Gammaproteobacteria | Legionellales | Legionellaceae | Legionella | ASV_24, ASV_18 |
| Leaf | ASV_116 | Actinobacteria | Actinobacteria | Actinomycetales | Microbacteriaceae | Microbacterium | ASV_7 |
| Leaf | ASV_12 | Spirochaetes | Spirochaetia | Spirochaetales | Treponemataceae | Treponema | ASV_531, ASV_4 |
| Leaf | ASV_121 | Pseudomonadota | Betaproteobacteria | Burkholderiales | Oxalobacteraceae | Massilia | ASV_53 |
| Leaf | ASV_1246 | Proteobacteria | Alphaproteobacteria | Rhizobiales | Aurantimonadaceae | Aurantimonas | ASV_144 |
| Leaf | ASV_13 | Proteobacteria | Alphaproteobacteria | Sphingomonadales | Sphingomonadaceae | Sphingopyxis | ASV_256 |
| Leaf | ASV_1306 | Pseudomonadota | Alphaproteobacteria | Sphingomonadales | Sphingomonadaceae | Sphingomonas | ASV_115 |
| Leaf | ASV_1454 | Actinobacteria | Actinobacteria | Actinomycetales | Intrasporangiaceae | Kineococcus | ASV_102 |
| Leaf | ASV_166 | Dependentiae | unclassified | unclassified | unclassified | unclassified | ASV_66 |
| Leaf | ASV_1729 | Pseudomonadota | Alphaproteobacteria | Rhodospirillales | Azospirillaceae | Skermanella | ASV_2 |
| Leaf | ASV_1751 | Pseudomonadota | Alphaproteobacteria | Sphingomonadales | Sphingomonadaceae | Sphingomonas | ASV_138 |
| Leaf | ASV_189 | Fusobacteria | Fusobacteriia | Fusobacteriales | Leptotrichiaceae | Sebaldella | ASV_259 |
| Leaf | ASV_2248 | Pseudomonadota | Alphaproteobacteria | Sphingomonadales | Sphingomonadaceae | Sphingomonas | ASV_473 |
| Leaf | ASV_2996 | Fusobacteria | Fusobacteriia | Fusobacteriales | Fusobacteriaceae | Psychrilyobacter | ASV_916 |
| Leaf | ASV_3132 | Proteobacteria | Gammaproteobacteria | Xanthomonadales | Xanthomonadaceae | Xanthomonas | ASV_225 |
| Leaf | ASV_320 | Pseudomonadota | Alphaproteobacteria | Sphingomonadales | Sphingomonadaceae | Sphingomonas | ASV_2 |
| Leaf | ASV_3230 | Spirochaetes | Spirochaetia | Spirochaetales | Spirochaetaceae | Thiospirochaeta | ASV_177 |
| Leaf | ASV_3885 | Dependentiae | unclassified | unclassified | unclassified | unclassified | ASV_130 |
| Leaf | ASV_4017 | Fusobacteria | Fusobacteriia | Fusobacteriales | Leptotrichiaceae | Streptobacillus | ASV_58 |
| Leaf | ASV_4236 | Actinobacteria | Actinobacteria | Actinomycetales | Actinospicaceae | Actinospica | ASV_7 |
| Leaf | ASV_443 | Actinomycetota | Actinobacteria | Corynebacteriales | Mycobacteriaceae | Mycobacterium | ASV_1962, ASV_473 |
| Leaf | ASV_45 | Actinobacteria | Actinobacteria | Actinomycetales | Microbacteriaceae | Curtobacterium | ASV_20 |
| Leaf | ASV_4693 | Actinobacteria | Actinobacteria | Actinomycetales | Microbacteriaceae | Herbiconiux | ASV_4 |
| Leaf | ASV_474 | Acidobacteria | Acidobacteriia | Acidobacteriales | Acidobacteriaceae | Terriglobus | ASV_2 |
| Leaf | ASV_4860 | Actinomycetota | Actinobacteria | Pseudonocardiales | Pseudonocardiaceae | Saccharothrix | ASV_55 |
| Leaf | ASV_4880 | Bacteroidota | Cytophagia | Cytophagales | Hymenobacteraceae | Hymenobacter | ASV_1044 |
| Leaf | ASV_6 | Proteobacteria | Alphaproteobacteria | Rhizobiales | Bradyrhizobiaceae | Bosea | ASV_1833, ASV_458, ASV_130 |
| Leaf | ASV_626 | Bacteroidetes | Cytophagia | Cytophagales | Cytophagaceae | Spirosoma | ASV_53, ASV_144 |
| Leaf | ASV_6310 | Pseudomonadota | Epsilonproteobacteria | Campylobacterales | Campylobacteraceae | Campylobacter | ASV_66 |
| Leaf | ASV_6358 | Pseudomonadota | Myxococcales | Myxococcales | Myxococcaceae | Anaeromyxobacter | ASV_2659 |

**Table S3** (continued)

| Compartment | ASV | Phylum | Class | Order | Family | Genus | Connections (fungal) |
|---|---|---|---|---|---|---|---|
| Leaf | ASV_747 | Proteobacteria | Alphaproteobacteria | Hyphomicrobiales | Methylobacteriaceae | Methylobacterium | ASV_17 |
| Leaf | ASV_842 | Bacteria | Firmicutes | Clostridia | Clostridiales | Lachnospiraceae | ASV_58 |
| Leaf | ASV_9 | Proteobacteria | Alphaproteobacteria | Hyphomicrobiales | Rhabdaerophilaceae | Rhabdaerophilum | ASV_3373 |
| Leaf | ASV_929 | Bacteria | Proteobacteria | unclassified | Alphaproteobacteria | Micavibrio | ASV_744 |
| Leaf | ASV_993 | Proteobacteria | Alphaproteobacteria | Rhizobiales | Methylocystaceae | Methylosinus | ASV_406 |
| Root | ASV_103 | Verrucomicrobia | unclassified | Terrimicrobiales | Terrimicrobiaceae | Chthoniobacter | ASV_282 |
| Root | ASV_1334 | Pseudomonadota | Alphaproteobacteria | Hyphomicrobiales | Xanthobacteraceae | Labrys | ASV_150 |
| Root | ASV_143 | Pseudomonadota | Gammaproteobacteria | Chromatiales | unclassified | Thiohalobacter | ASV_443 |
| Root | ASV_165 | Bacteroidetes | Sphingobacteriia | Sphingobacteriales | Chitinophagaceae | Niastella | ASV_319 |
| Root | ASV_2 | Proteobacteria | Alphaproteobacteria | Rhizobiales | Bradyrhizobiaceae | Afipia | ASV_370 |
| Root | ASV_205 | Actinobacteria | Actinobacteria | Actinomycetales | Frankiaceae | Frankia | ASV_375 |
| Root | ASV_21 | Bacillota | Clostridia | Eubacteriales | Lachnospiraceae | [Clostridium] | ASV_291 |
| Root | ASV_215 | Pseudomonadota | Myxococcales | Myxococcales | Polyangiaceae | Aetherobacter | ASV_416 |
| Root | ASV_24 | Verrucomicrobia | unclassified | Terrimicrobiales | Terrimicrobiaceae | Chthoniobacter | ASV_107 |
| Root | ASV_264 | Proteobacteria | Alphaproteobacteria | Rhizobiales | unclassified | Agrobacterium | ASV_240 |
| Root | ASV_28 | Proteobacteria | Alphaproteobacteria | Sphingomonadales | Erythrobacteraceae | Porphyrobacter | ASV_26 |
| Root | ASV_31 | Proteobacteria | Alphaproteobacteria | Rhizobiales | Bradyrhizobiaceae | Afipia | ASV_231 |
| Root | ASV_322 | Acidobacteria | Solibacteres | Solibacterales | Solibacteraceae | Solibacter | ASV_790 |
| Root | ASV_35 | Proteobacteria | Gammaproteobacteria | Enterobacteriales | Enterobacteriaceae | Pseudomonas | ASV_38 |
| Root | ASV_371 | Pseudomonadota | Alphaproteobacteria | Sphingomonadales | Sphingomonadaceae | Hankyongella | ASV_56 |
| Root | ASV_38 | Actinobacteria | Actinobacteria | Actinomycetales | Streptomycetaceae | Streptomyces | ASV_1787, ASV_334 |
| Root | ASV_397 | Proteobacteria | Gammaproteobacteria | Xanthomonadales | Xanthomonadaceae | Frateuria | ASV_1787 |
| Root | ASV_402 | Proteobacteria | Alphaproteobacteria | Rhizobiales | unclassified | Vasilyevaea | ASV_12 |
| Root | ASV_41 | Proteobacteria | Alphaproteobacteria | Rhodospirillales | Rhodospirillaceae | Dongia | ASV_231 |
| Root | ASV_442 | Proteobacteria | Alphaproteobacteria | Rhodospirillales | Rhodospirillaceae | Dongia | ASV_179 |
| Root | ASV_503 | Proteobacteria | Betaproteobacteria | Burkholderiales | Burkholderiaceae | Glomeribacter | ASV_113, ASV_161 |
| Root | ASV_512 | Actinobacteria | Actinobacteria | Actinomycetales | Thermomonosporaceae | Actinomadura | ASV_169 |
| Root | ASV_539 | Proteobacteria | Betaproteobacteria | Burkholderiales | Alcaligenaceae | Azohydromonas | ASV_150 |
| Root | ASV_595 | Deltaproteobacteria | Myxococcales | Myxococcales | Nannocystaceae | Nannocystis | ASV_827 |
| Root | ASV_61 | Actinobacteria | Actinobacteria | Actinomycetales | Micromonosporaceae | Longispora | ASV_38 |
| Root | ASV_625 | Actinomycetota | Rubrobacteridae | Gaiellales | Gaiellaceae | Gaiella | ASV_248 |
| Root | ASV_630 | Deltaproteobacteria | Myxococcales | Nannocystineae | Kofleriaceae | Haliangium | ASV_254 |
| Root | ASV_663 | Actinomycetota | Rubrobacteridae | Gaiellales | Gaiellaceae | Gaiella | ASV_107 |
| Root | ASV_69 | Pseudomonadota | Alphaproteobacteria | Hyphomicrobiales | Hyphomicrobiaceae | Rhodoplanes | ASV_567 |

**Table S3** (continued)

| Compartment | ASV | Phylum | Class | Order | Family | Genus | Connections (fungal) |
|---|---|---|---|---|---|---|---|
| Root | ASV_699 | Proteobacteria | Betaproteobacteria | Rhodocyclales | Rhodocyclaceae | Propionivibrio | ASV_140 |
| Root | ASV_83 | Acidobacteria | Acidobacteriia | Acidobacteriales | Acidobacteriaceae | Koribacter | ASV_231 |
| Root | ASV_84 | Pseudomonadota | Gammaproteobacteria | unclassified | unclassified | Acidibacter | ASV_209 |
| Root | ASV_887 | Pseudomonadota | Alphaproteobacteria | Sphingomonadales | Sphingomonadaceae | Sphingomonas | ASV_134 |
| Root | ASV_89 | Actinobacteria | Actinobacteria | Actinomycetales | Streptomycetaceae | Streptomyces | ASV_5 |
| Root | ASV_933 | Proteobacteria | Alphaproteobacteria | Sphingomonadales | Sphingomonadaceae | Blastomonas | ASV_654 |
| Root | ASV_96 | Rokubacteria | unclassified | unclassified | unclassified | unclassified | ASV_57 |

**Table S4.** Fungal taxa involved in putative direct cross-kingdom interactions identified in the network using an edge-wise variability estimate of 0.8. Assignments result from placing unknown sequences on T-BAS phylogenetic reference tree.

| Compartment | ASV | Phylum | Class | Order | Family | Genus | Connections (bacterial) |
|---|---|---|---|---|---|---|---|
| Leaf | ASV_102 | Ascomycota | Dothideomycetes | Pleosporales | Pyrenochaetopsidaceae | Pyrenochaetopsis | ASV_1454 |
| Leaf | ASV_1044 | Basidiomycota | Cystobasidiomycetes | Erythrobasidiales | Erythrobasidiaceae | Bannoa | ASV_4880 |
| Leaf | ASV_115 | Basidiomycota | Exobasidiomycetes | Exobasidiales | Brachybasidiaceae | Meira | ASV_1306 |
| Leaf | ASV_130 | Basidiomycota | Tremellomycetes | Tremellales | Sirobasidiaceae | Fibulobasidium | ASV_6, ASV_3885 |
| Leaf | ASV_138 | Basidiomycota | Cystobasidiomycetes | Erythrobasidiales | Erythrobasidiaceae | Microsporomyces | ASV_1751, ASV_1145 |
| Leaf | ASV_144 | Ascomycota | Dothideomycetes | Pleosporales | Coniothyriaceae | Neoconiothyrium | ASV_1246, ASV_626 |
| Leaf | ASV_17 | Ascomycota | Dothideomycetes | Pleosporales | Didymellaceae | Neodidymelliopsis | ASV_747 |
| Leaf | ASV_177 | Ascomycota | Sordariomycetes | Xylariales | Apiosporaceae | Nigrospora | ASV_3230 |
| Leaf | ASV_18 | Basidiomycota | Pucciniomycetes | Pucciniales | Raveneliaceae | Endoraecium | ASV_11551 |
| Leaf | ASV_1833 | Basidiomycota | Agaricomycetes | Agaricales | Porotheleaceae | Chrysomycena | ASV_6 |
| Leaf | ASV_1962 | Basidiomycota | Exobasidiomycetes | Exobasidiales | Brachybasidiaceae | Meira | ASV_443 |
| Leaf | ASV_2 | Ascomycota | Sordariomycetes | Xylariales | Microdochiaceae | Microdochium | ASV_474, ASV_1729, ASV_320 |
| Leaf | ASV_20 | Ascomycota | Dothideomycetes | Pleosporales | Phaeosphaeriaceae | Poaceicola | ASV_45 |
| Leaf | ASV_225 | Ascomycota | Dothideomycetes | Pleosporales | Phaeosphaeriaceae | Pseudostaurosphaeria | ASV_3132 |
| Leaf | ASV_24 | Ascomycota | Dothideomycetes | Pleosporales | Phaeosphaeriaceae | Setophoma | ASV_11551 |
| Leaf | ASV_256 | Ascomycota | Dothideomycetes | Pleosporales | Phaeosphaeriaceae | Phaeosphaeria | ASV_13 |
| Leaf | ASV_259 | Ascomycota | Eurotiomycetes | Chaetothyriales | Herpotrichiellaceae | Phialophora | ASV_189 |
| Leaf | ASV_2659 | Basidiomycota | Agaricostilbomycetes | Agaricostilbales | Chionosphaeraceae | Cystobasidiopsis | ASV_6358 |
| Leaf | ASV_3373 | Basidiomycota | Agaricostilbomycetes | Agaricostilbales | Chionosphaeraceae | Sterigmatomyces | ASV_9 |
| Leaf | ASV_4 | Ascomycota | Dothideomycetes | Mycosphaerellales | Dissoconiaceae | Paradissoconium | ASV_4693, ASV_12 |
| Leaf | ASV_406 | Basidiomycota | Tremellomycetes | Tremellales | Sirobasidiaceae | Fibulobasidium | ASV_993 |
| Leaf | ASV_458 | Basidiomycota | Tremellomycetes | Tremellales | Bulleribasidiaceae | Bulleribasidium | ASV_6 |
| Leaf | ASV_473 | Ascomycota | Sordariomycetes | Xylariales | Apiosporaceae | Nigrospora | ASV_443, ASV_2248 |
| Leaf | ASV_53 | Ascomycota | Dothideomycetes | Cladosporiales | Cladosporiaceae | Melanodothis | ASV_121, ASV_626 |
| Leaf | ASV_531 | Ascomycota | Eurotiomycetes | Chaetothyriales | Cyphellophoraceae | Anthopsis | ASV_12 |
| Leaf | ASV_55 | Basidiomycota | Tremellomycetes | Tremellales | Bulleribasidiaceae | Hannaella | ASV_4860 |
| Leaf | ASV_58 | Ascomycota | Dothideomycetes | Pleosporales | Phaeosphaeriaceae | Didymocyrtis | ASV_842, ASV_4017 |
| Leaf | ASV_66 | Ascomycota | Dothideomycetes | Pleosporales | Didymellaceae | Neodidymelliopsis | ASV_6310, ASV_166 |
| Leaf | ASV_7 | Ascomycota | Dothideomycetes | Pleosporales | Leptosphaeriaceae | Sphaerellopsis | ASV_116, ASV_4236 |
| Leaf | ASV_744 | Ascomycota | Dothideomycetes | Pleosporales | Phaeosphaeriaceae | Brunneomurispora | ASV_929 |

**Table S4** (continued)

| Compartment | ASV | Phylum | Class | Order | Family | Genus | Connections (bacterial) |
|---|---|---|---|---|---|---|---|
| Leaf | ASV_856 | Basidiomycota | Exobasidiomycetes | Exobasidiales | Brachybasidiaceae | Meira | ASV_1151 |
| Leaf | ASV_916 | Basidiomycota | Exobasidiomycetes | Microstromatales | unclassified | Pseudomicrostroma | ASV_2996 |
| Root | ASV_107 | Ascomycota | Dothideomycetes | Pleosporales | Lentitheciaceae | Poaceascoma | ASV_24, ASV_663 |
| Root | ASV_113 | Ascomycota | Leotiomycetes | unclassified | unclassified | Helicocentralis | ASV_503 |
| Root | ASV_12 | Ascomycota | Leotiomycetes | Helotiales | Leptodontidiaceae | Leptodontidium | ASV_402 |
| Root | ASV_134 | Mucoromycota | Glomeromycetes | Diversisporales | Glomeraceae | Paraglomus | ASV_887 |
| Root | ASV_140 | Basidiomycota | Tremellomycetes | Cystofilobasidiales | Mrakiaceae | Mrakia | ASV_699 |
| Root | ASV_150 | Basidiomycota | Agaricomycetes | Polyporales | Meripilaceae | Physisporinus | ASV_1334, ASV_539 |
| Root | ASV_161 | Ascomycota | Dothideomycetes | Pleosporales | Pyrenochaetopsidaceae | Pyrenochaetopsis | ASV_503 |
| Root | ASV_169 | Ascomycota | Eurotiomycetes | Chaetothyriales | Trichomeriaceae | Knufia | ASV_512 |
| Root | ASV_1787 | Ascomycota | Saccharomycetes | Saccharomycetales | Saccharomycetaceae | Citeromyces | ASV_38, ASV_397 |
| Root | ASV_179 | Chytridiomycota | Chytridiomycetes | Lobulomycetales | Lobulomycetaceae | Quaeritorhiza | ASV_442 |
| Root | ASV_209 | Mucoromycota | Glomeromycetes | Diversisporales | Glomeraceae | Paraglomus | ASV_84 |
| Root | ASV_231 | Ascomycota | Eurotiomycetes | Chaetothyriales | Herpotrichiellaceae | Cladophialophora | ASV_31, ASV_41, ASV_83 |
| Root | ASV_240 | Ascomycota | Dothideomycetes | Pleosporales | Lentitheciaceae | Poaceascoma | ASV_264 |
| Root | ASV_248 | Ascomycota | Eurotiomycetes | Chaetothyriales | unclassified | Strelitziana | ASV_625 |
| Root | ASV_254 | Mucoromycota | Glomeromycetes | Diversisporales | Glomeraceae | Paraglomus | ASV_630 |
| Root | ASV_26 | Basidiomycota | Agaricomycetes | Corticiales | Corticiaceae | Subulicystidium | ASV_28 |
| Root | ASV_282 | Basidiomycota | Agaricomycetes | Cantharellales | unclassified | Hydnum | ASV_103 |
| Root | ASV_291 | Ascomycota | Eurotiomycetes | Chaetothyriales | Trichomeriaceae | Knufia | ASV_21 |
| Root | ASV_319 | Mucoromycota | Glomeromycetes | Diversisporales | Glomeraceae | Paraglomus | ASV_165 |
| Root | ASV_334 | Basidiomycota | Agaricomycetes | Cantharellales | unclassified | Hydnum | ASV_38 |
| Root | ASV_370 | Mucoromycota | Mortierellomycetes | Mortierellales | Mortierellaceae | Entomortierella | ASV_2 |
| Root | ASV_375 | Ascomycota | Leotiomycetes | unclassified | unclassified | Helicocentralis | ASV_205 |
| Root | ASV_38 | Ascomycota | Eurotiomycetes | Chaetothyriales | Trichomeriaceae | Strelitziana | ASV_35, ASV_61 |
| Root | ASV_416 | Basidiomycota | Agaricomycetes | Cantharellales | unclassified | Hydnum | ASV_215 |
| Root | ASV_443 | Basidiomycota | Tremellomycetes | Filobasidiales | Piskurozymaceae | Solicoccozyma | ASV_143 |
| Root | ASV_5 | Basidiomycota | Ustilaginomycetes | Ustilaginales | Ustilaginaceae | Shivasia | ASV_89 |
| Root | ASV_56 | Basidiomycota | Tremellomycetes | Tremellales | Trimorphomycetaceae | Sugitazyma | ASV_371 |
| Root | ASV_567 | Mucoromycota | Glomeromycetes | Diversisporales | Glomeraceae | Paraglomus | ASV_69 |
| Root | ASV_57 | Basidiomycota | Agaricomycetes | Agaricales | Entolomataceae | Entoloma | ASV_96 |
| Root | ASV_654 | Chytridiomycota | Chytridiomycetes | Chytridiales | Chytriomycetaceae | Rhizidium | ASV_933 |
| Root | ASV_790 | Mucoromycota | Glomeromycetes | Diversisporales | Glomeraceae | Paraglomus | ASV_322 |
| Root | ASV_827 | Ascomycota | Dothideomycetes | Pleosporales | Pyrenochaetopsidaceae | Pyrenochaetopsis | ASV_595 |

**Table S5.** Bacterial taxa involved in putative diffuse (keystone) cross-kingdom interactions identified using an edge-wise variability estimate of 0.8 and 75th percentile threshold for degree and eigenvector centrality to define keystoneness. Assignments result from placing unknown sequences on T-BAS reference tree.

| Compart-ment | ASV | Phylum | Class | Order | Family | Genus | Connections (within-kingdom) |
|---|---|---|---|---|---|---|---|
| Leaf | ASV_5911 | Pseudomonadota | Gammaproteobacteria | unclassified | unclassified | Acidibacter | ASV_39, ASV_6715, ASV_7872, ASV_8842, ASV_8527, ASV_2026 |
| Leaf | ASV_689 | Actinomycetota | Actinobacteria | Pseudonocardiales | Pseudonocardiaceae | Actinomycetospora | ASV_2370, ASV_2248, ASV_743, ASV_5187, ASV_1997, ASV_3422, ASV_6164, ASV_768, ASV_1275, ASV_12 |
| Leaf | ASV_599 | Deltaproteobacteria | Myxococcales | Myxococcales | Myxococcaceae | Aggregicoccus | ASV_474, ASV_4016, ASV_748, ASV_10941, ASV_3886 |
| Leaf | ASV_768 | Deltaproteobacteria | Myxococcales | Myxococcales | Myxococcaceae | Aggregicoccus | ASV_597, ASV_689, ASV_1131, ASV_1495, ASV_9238, ASV_887, ASV_2009 |
| Leaf | ASV_4016 | Pseudomonadota | Alphaproteobacteria | Hyphomicrobiales | Aurantimonadaceae | Aureimonas | ASV_597, ASV_1751, ASV_443, ASV_599, ASV_1788 |
| Leaf | ASV_4622 | Proteobacteria | Alphaproteobacteria | Rhizobiales | Beijerinckiaceae | Beijerinckia | ASV_3967, ASV_887, ASV_1997, ASV_3046, ASV_637, ASV_7315, ASV_702, ASV_443 |
| Leaf | ASV_597 | Actinobacteria | Actinobacteria | Actinomycetales | Beutenbergiaceae | Beutenbergia | ASV_4016, ASV_4350, ASV_768, ASV_2009, ASV_2026, ASV_470 |
| Leaf | ASV_682 | Proteobacteria | Alphaproteobacteria | Sphingomonadales | Sphingomonadaceae | Blastomonas | ASV_1794, ASV_3422, ASV_4005, ASV_3298, ASV_775 |
| Leaf | ASV_2009 | Proteobacteria | Alphaproteobacteria | Rhizobiales | Beijerinckiaceae | Chelatococcus | ASV_597, ASV_470, ASV_768, ASV_8842, ASV_4546 |
| Leaf | ASV_2370 | Armatimonadetes | unclassified | unclassified | unclassified | Fimbriimonas | ASV_531, ASV_689, ASV_3274, ASV_3051, ASV_9928, ASV_6915 |
| Leaf | ASV_3051 | Armatimonadetes | unclassified | unclassified | unclassified | Fimbriimonas | ASV_748, ASV_4921, ASV_3846, ASV_7616, ASV_2370 |
| Leaf | ASV_136 | Proteobacteria | Alphaproteobacteria | Rhizobiales | Aurantimonadaceae | Fulvimarina | ASV_531, ASV_3142, ASV_3298, ASV_3422, ASV_659 |
| Leaf | ASV_199 | Proteobacteria | Alphaproteobacteria | Rhizobiales | Aurantimonadaceae | Martelella | ASV_2657, ASV_528, ASV_991, ASV_1247, ASV_497, ASV_2248 |
| Leaf | ASV_571 | Proteobacteria | Betaproteobacteria | Burkholderiales | Oxalobacteraceae | Massilia | ASV_7872, ASV_4005, ASV_1729, ASV_1714, ASV_2936, ASV_340 |
| Leaf | ASV_470 | Proteobacteria | Alphaproteobacteria | Hyphomicrobiales | Methylobacteriaceae | Methylobacterium | ASV_3682, ASV_9928, ASV_1503, ASV_1921, ASV_3276, ASV_597, ASV_2248, ASV_4350, ASV_1270, ASV_2009, ASV_8842 |
| Leaf | ASV_4005 | Actinomycetota | Actinobacteria | Propionibacteriales | Nocardioidaceae | Nocardioides | ASV_929, ASV_571, ASV_443, ASV_656, ASV_682 |
| Leaf | ASV_4059 | Actinomycetota | Actinobacteria | Propionibacteriales | Nocardioidaceae | Nocardioides | ASV_1176, ASV_689, ASV_5502, ASV_7616, ASV_9010 |
| Leaf | ASV_4546 | Actinomycetota | Actinobacteria | Propionibacteriales | Nocardioidaceae | Nocardioides | ASV_1921, ASV_237, ASV_112, ASV_9010, ASV_3422, ASV_2009 |
| Leaf | ASV_112 | Proteobacteria | Betaproteobacteria | Burkholderiales | Comamonadaceae | Ottowia | ASV_8591, ASV_227, ASV_811, ASV_1246, ASV_4350, ASV_5502, ASV_1794, ASV_4546 |
| Leaf | ASV_6715 | Proteobacteria | Gammaproteobacteria | Xanthomonadales | Xanthomonadaceae | Pseudoxanthomonas | ASV_121, ASV_2034, ASV_5911, ASV_7872, ASV_9928 |
| Leaf | ASV_743 | Proteobacteria | Betaproteobacteria | Burkholderiales | Comamonadaceae | Ramlibacter | ASV_391, ASV_406, ASV_689, ASV_1151, ASV_3422 |
| Leaf | ASV_8842 | Proteobacteria | Alphaproteobacteria | Rickettsiales | Rickettsiaceae | Rickettsia | ASV_470, ASV_10204, ASV_4415, ASV_5911, ASV_2009 |
| Leaf | ASV_237 | Pseudomonadota | Alphaproteobacteria | Sphingomonadales | Sphingomonadaceae | Sphingomonas | ASV_39, ASV_929, ASV_391, ASV_4460, ASV_1246, ASV_3142, ASV_4546, ASV_5912 |
| Leaf | ASV_3422 | Pseudomonadota | Alphaproteobacteria | Sphingomonadales | Sphingomonadaceae | Sphingomonas | ASV_1306, ASV_136, ASV_682, ASV_689, ASV_743, ASV_4546 |
| Leaf | ASV_9928 | Spirochaetes | Spirochaetia | Spirochaetales | Spirochaetaceae | Spirochaeta | ASV_3615, ASV_188, ASV_470, ASV_6715, ASV_340, ASV_2370 |
| Root | ASV_1018 | Pseudomonadota | Alphaproteobacteria | Hyphomicrobiales | Hyphomicrobiaceae | Filomicrobium | ASV_128, ASV_150, ASV_157, ASV_168, ASV_207, ASV_247, ASV_390, ASV_411, ASV_510, ASV_933 |
| Root | ASV_1042 | Proteobacteria | Alphaproteobacteria | Caulobacterales | Caulobacteraceae | Asticcacaulis | ASV_114, ASV_169, ASV_172, ASV_245, ASV_26, ASV_394, ASV_403, ASV_683 |
| Root | ASV_1078 | Acidobacteria | unclassified | Thermoanaerobaculales | Thermoanaerobaculaceae | Thermoanaerobaculum | ASV_230, ASV_254, ASV_263, ASV_321, ASV_366, ASV_390, ASV_591, ASV_683 |
| Root | ASV_1084 | Acidobacteria | Acidobacteriia | Acidobacteriales | Acidobacteriaceae | Acidobacterium | ASV_118, ASV_32, ASV_331, ASV_376, ASV_377, ASV_433, ASV_551, ASV_677, ASV_84, ASV_848 |

**Table S5** (continued)

| Compart-ment | ASV | Phylum | Class | Order | Family | Genus | Connections (within-kingdom) |
|---|---|---|---|---|---|---|---|
| Root | ASV_109 | Actinomycetota | Rubrobacteridae | Gaiellales | Gaiellaceae | Gaiella | ASV_1237, ASV_124, ASV_1436, ASV_150, ASV_1587, ASV_200, |
| Root | ASV_1115 | Acidobacteria | Acidobacteriia | Acidobacteriales | Acidobacteriaceae | Acidicapsa | ASV_1139, ASV_125, ASV_147, ASV_1436, ASV_191, ASV_193, ASV_254, ASV_314, ASV_469, ASV_532, ASV_77, ASV_944, ASV_949 |
| Root | ASV_1139 | Chloroflexi | Ktedonobacteria | Ktedonobacterales | Ktedonosporobacteraceae | Ktedonosporobacter | ASV_1115, ASV_1237, ASV_212, ASV_249, ASV_378, ASV_402, ASV_469, ASV_774, ASV_818 |
| Root | ASV_118 | Acidobacteria | Acidobacteriia | Acidobacteriales | Acidobacteriaceae | Koribacter | ASV_1084, ASV_1237, ASV_132, ASV_1436, ASV_1587, ASV_171, ASV_331, ASV_677, ASV_848, ASV_84, ASV_92 |
| Root | ASV_1237 | Verrucomicrobia | Opitutae | Opitutales | Opitutaceae | Opitutus | ASV_109, ASV_1139, ASV_118, ASV_1436, ASV_1587, ASV_124, ASV_182, ASV_274, ASV_295, ASV_322, ASV_366, ASV_61, ASV_677, ASV_712, ASV_92, ASV_920 |
| Root | ASV_124 | Gemmatimonadetes | unclassified | Gemmatimonadales | Gemmatimonadaceae | Gemmatimonas | ASV_109, ASV_1237, ASV_1436, ASV_150, ASV_1587, ASV_436, ASV_712, ASV_74, ASV_90 |
| Root | ASV_1292 | Actinomycetota | Actinobacteria | Corynebacteriales | Mycobacteriaceae | Mycobacterium | ASV_147, ASV_254, ASV_288, ASV_318, ASV_513, ASV_587, ASV_644 |
| Root | ASV_1369 | Pseudomonadota | Myxococcales | Myxococcales | Polyangiaceae | Aetherobacter | ASV_154, ASV_245, ASV_26, ASV_260, ASV_306, ASV_403, ASV_673, ASV_791 |
| Root | ASV_1395 | Chloroflexi | unclassified | Phototrophicales | Phototrophicaceae | Phototrophicus | ASV_1287, ASV_147, ASV_231, ASV_247, ASV_292, ASV_307, ASV_32, ASV_324, ASV_407 |
| Root | ASV_1436 | Chloroflexi | Ktedonobacteria | Ktedonobacterales | Ktedonosporobacteraceae | Ktedonosporobacter | ASV_109, ASV_1115, ASV_118, ASV_1237, ASV_124, ASV_1587, ASV_233, ASV_249, ASV_677, ASV_712, ASV_963 |
| Root | ASV_147 | Verrucomicrobia | unclassified | Terrimicrobiales | Terrimicrobiaceae | Chthoniobacter | ASV_1115, ASV_1292, ASV_1395, ASV_254, ASV_292, ASV_307, ASV_330, ASV_411, ASV_718, ASV_84, ASV_949 |
| Root | ASV_149 | Actinomycetota | Actinobacteria | Streptosporangiales | Treboniaceae | Trebonia | ASV_132, ASV_1359, ASV_1155, ASV_1334, ASV_1496, ASV_2100, ASV_266, ASV_302, ASV_469, ASV_529, ASV_800, ASV_967 |
| Root | ASV_150 | Deltaproteobacteria | Myxococcales | Myxococcales | Cystobacteraceae | Cystobacter | ASV_102, ASV_109, ASV_124, ASV_126, ASV_19, ASV_1018, ASV_436, ASV_456, ASV_712 |
| Root | ASV_1587 | Bacillota | Bacilli | Bacillales | Paenibacillaceae | Ammoniphilus | ASV_1064, ASV_109, ASV_118, ASV_1237, ASV_124, ASV_1436, AV_21, ASV_677, ASV_712, ASV_92 |
| Root | ASV_164 | Actinomycetota | Actinobacteria | Actinopolysporales | Actinopolysporaceae | Actinopolyspora | ASV_16, ASV_171, ASV_191, ASV_397, ASV_532, ASV_615, ASV_77, ASV_84, ASV_848 |
| Root | ASV_171 | Actinomycetota | Actinobacteria | Streptosporangiales | Treboniaceae | Trebonia | ASV_118, ASV_146, ASV_164, ASV_331, ASV_397, ASV_77, ASV_790, ASV_799, ASV_84, ASV_848 |
| Root | ASV_191 | Actinomycetota | Actinobacteria | Corynebacteriales | Mycobacteriaceae | Mycobacterium | ASV_125, ASV_15, ASV_164, ASV_1115, ASV_397, ASV_411, ASV_433, ASV_532, ASV_696, ASV_77, ASV_966 |
| Root | ASV_193 | Acidobacteria | Acidobacteriia | Acidobacteriales | Acidobacteriaceae | Koribacter | ASV_148, ASV_17, ASV_1115, ASV_2100, ASV_233, ASV_269, ASV_314, ASV_944 |
| Root | ASV_200 | Pseudomonadota | Gammaproteobacteria | Immundisolibacterales | Immundisolibacteraceae | Immundisolibacter | ASV_109, ASV_146, ASV_196, ASV_223, ASV_306, ASV_663, ASV_77 |
| Root | ASV_207 | Acidobacteria | unclassified | Bryobacterales | Bryobacteraceae | Bryobacter | ASV_128, ASV_157, ASV_205, ASV_1018, ASV_355, ASV_666, ASV_949 |
| Root | ASV_216 | Pseudomonadota | Alphaproteobacteria | Hyphomicrobiales | unclassified | Enhydrobacter | ASV_106, ASV_126, ASV_143, ASV_1184, ASV_1287, ASV_314, ASV_385, ASV_394, ASV_570, ASV_32, ASV_78, ASV_83 |
| Root | ASV_241 | Proteobacteria | Alphaproteobacteria | Sphingomonadales | Sphingomonadaceae | Sphingopyxis | ASV_190, ASV_394, ASV_481, ASV_551, ASV_595, ASV_666, ASV_59 |
| Root | ASV_245 | Acidobacteria | Acidobacteriia | Acidobacteriales | Acidobacteriaceae | Acidobacterium | ASV_213, ASV_1042, ASV_1184, ASV_119, ASV_1369, ASV_1826, ASV_407, ASV_414, ASV_433, ASV_683, ASV_26 |
| Root | ASV_249 | Actinobacteria | Rubrobacteridae | Solirubrobacterales | unclassified | Bactoderma | ASV_211, ASV_1139, ASV_1436, ASV_512, ASV_570, ASV_589, ASV_51 |
| Root | ASV_254 | Chloroflexi | unclassified | unclassified | unclassified | unclassified | ASV_147, ASV_1078, ASV_115, ASV_1292, ASV_288, ASV_390, ASV_411, ASV_427, ASV_644, ASV_726, ASV_933, ASV_949, ASV_992, ASV_74, ASV_86 |

**Table S5** (continued)

| Compart-ment | ASV | Phylum | Class | Order | Family | Genus | Connections (within-kingdom) |
|---|---|---|---|---|---|---|---|
| Root | ASV_26 | Verrucomicrobia | unclassified | Terrimicrobiales | Terrimicrobiaceae | Chthoniobacter | ASV_1042, ASV_1369, ASV_1826, ASV_245, ASV_403, ASV_510, ASV_745 |
| Root | ASV_269 | Deltaproteobacteria | Myxococcales | Nannocystineae | Kofleriaceae | Haliangium | ASV_193, ASV_2100, ASV_295, ASV_306, ASV_349, ASV_414, ASV_469, ASV_615, ASV_677, ASV_84 |
| Root | ASV_292 | Proteobacteria | Alphaproteobacteria | Sphingomonadales | Sphingomonadaceae | Blastomonas | ASV_147, ASV_1014, ASV_1395, ASV_307, ASV_380, ASV_966, ASV_59 |
| Root | ASV_306 | Deltaproteobacteria | Myxococcales | Nannocystineae | Kofleriaceae | Haliangium | ASV_109, ASV_173, ASV_196, ASV_200, ASV_269, ASV_1071, ASV_1369, ASV_663, ASV_790 |
| Root | ASV_307 | Pseudomonadota | Deltaproteobacteria | unclassified | unclassified | Deferrisoma | ASV_134, ASV_147, ASV_209, ASV_292, ASV_1395, ASV_510, ASV_966, ASV_48, ASV_90 |
| Root | ASV_314 | Pseudomonadota | Myxococcales | Myxococcales | Labilitrichaceae | Labilithrix | ASV_14, ASV_193, ASV_216, ASV_1115, ASV_377, ASV_425, ASV_460, ASV_565, ASV_570, ASV_944 |
| Root | ASV_32 | Pseudomonadota | Alphaproteobacteria | Micropepsales | Micropepsaceae | Micropepsis | ASV_1084, ASV_122, ASV_1395, ASV_216, ASV_376, ASV_377, ASV_551, ASV_879 |
| Root | ASV_330 | Pseudomonadota | Alphaproteobacteria | Hyphomicrobiales | Hyphomicrobiaceae | Filomicrobium | ASV_147, ASV_190, ASV_285, ASV_308, ASV_1761, ASV_371, ASV_551 |
| Root | ASV_331 | Bacteroidetes | Sphingobacteriia | Sphingobacteriales | Chitinophagaceae | Niastella | ASV_118, ASV_171, ASV_179, ASV_248, ASV_1084, ASV_1319, ASV_385, ASV_401, ASV_594, ASV_848, ASV_84 |
| Root | ASV_366 | Planctomycetota | Planctomycetia | Pirellulales | Thermoguttaceae | Thermogutta | ASV_163, ASV_167, ASV_215, ASV_1078, ASV_1237, ASV_1496, ASV_601, ASV_966 |
| Root | ASV_377 | Actinobacteria | Actinobacteria | Actinomycetales | Micromonosporaceae | Catelliglobosispora | ASV_314, ASV_32, ASV_324, ASV_355, ASV_376, ASV_1084, ASV_1391, ASV_425, ASV_551, ASV_565, 757 |
| Root | ASV_390 | Actinobacteria | Actinobacteria | Actinomycetales | Streptomycetaceae | Streptacidiphilus | ASV_217, ASV_230, ASV_254, ASV_263, ASV_321, ASV_1018, ASV_1078, ASV_403, ASV_591 |
| Root | ASV_394 | Actinomycetota | Rubrobacteridae | Gaiellales | Gaiellaceae | Gaiella | ASV_114, ASV_169, ASV_172, ASV_245, ASV_26, ASV_394, ASV_403, ASV_683 |
| Root | ASV_407 | Pseudomonadota | Alphaproteobacteria | Micropepsales | Micropepsaceae | Micropepsis | ASV_191, ASV_245, ASV_407, ASV_1084, ASV_1319, ASV_809, ASV_949 |
| Root | ASV_411 | Chloroflexi | Ktedonobacteria | Ktedonobacterales | Thermogemmatisporaceae | Thermogemmatispora | ASV_147, ASV_191, ASV_254, ASV_265, ASV_1018, ASV_558, ASV_818, ASV_949, ASV_963 |
| Root | ASV_414 | Proteobacteria | Betaproteobacteria | Burkholderiales | Burkholderiaceae | Burkholderia | ASV_213, ASV_223, ASV_245, 269, ASV_291, ASV_1184, ASV_550 |
| Root | ASV_433 | Proteobacteria | Alphaproteobacteria | Rhodospirillales | Rhodospirillaceae | Dongia | ASV_191, ASV_245, ASV_407, ASV_1084, ASV_1319, ASV_809, ASV_949 |
| Root | ASV_469 | Deltaproteobacteria | Myxococcales | Nannocystineae | Kofleriaceae | Haliangium | ASV_149, ASV_269, ASV_302, ASV_1115, ASV_1139, ASV_1184, ASV_529, ASV_589 |
| Root | ASV_510 | Proteobacteria | Alphaproteobacteria | Rhodospirillales | Rhodospirillaceae | Inquilinus | ASV_26, ASV_307, ASV_376, ASV_407, ASV_1018, ASV_1358, ASV_570 |
| Root | ASV_513 | Actinomycetota | Rubrobacteridae | Gaiellales | Gaiellaceae | Gaiella | ASV_123, ASV_139, ASV_163, ASV_230, ASV_288, ASV_378, ASV_1292, ASV_538, ASV_861 |
| Root | ASV_532 | Proteobacteria | Betaproteobacteria | Burkholderiales | Comamonadaceae | Verminephrobacter | ASV_125, ASV_164, ASV_191, ASV_217, ASV_318, ASV_397, ASV_1115, ASV_902, ASV_77 |
| Root | ASV_551 | Actinomycetota | Actinobacteria | Corynebacteriales | Mycobacteriaceae | Mycobacterium | ASV_241, ASV_28, ASV_30, ASV_32, ASV_330, ASV_376, ASV_377, ASV_394, ASV_1084, ASV_1231, ASV_595, ASV_666 |
| Root | ASV_558 | Pseudomonadota | Gammaproteobacteria | unclassified | unclassified | Acidibacter | ASV_114, ASV_133, ASV_411, ASV_543, ASV_1113, ASV_2166, ASV_673, ASV_690, ASV_949, ASV_967 |
| Root | ASV_570 | Actinomycetota | Actinobacteria | Streptosporangiales | Treboniaceae | Trebonia | ASV_211, ASV_216, ASV_249, ASV_314, ASV_4, ASV_510, ASV_512, ASV_538, ASV_570 |
| Root | ASV_644 | Pseudomonadota | Alphaproteobacteria | Hyphomicrobiales | Hyphomicrobiaceae | Filomicrobium | ASV_112, ASV_254, ASV_285, ASV_295, ASV_318, ASV_36, ASV_1292, ASV_726, ASV_933, ASV_70 |
| Root | ASV_650 | Actinomycetota | Actinobacteria | Corynebacteriales | Mycobacteriaceae | Mycobacterium | ASV_173, ASV_198, ASV_318, ASV_429, ASV_566, ASV_606, ASV_1560, ASV_1672, ASV_2166, ASV_654, ASV_949 |
| Root | ASV_666 | Proteobacteria | Gammaproteobacteria | Chromatiales | Ectothiorhodospiraceae | Thiohalospira | ASV_100, ASV_137, ASV_207, ASV_241, ASV_355, ASV_456, ASV_551, ASV_595, ASV_963, ASV_83 |
| Root | ASV_677 | Verrucomicrobia | Verrucomicrobiae | Verrucomicrobiales | Verrucomicrobia | Verrucomicrobia | ASV_118, ASV_127, ASV_269, ASV_1084, ASV_1237, ASV_1436, ASV_1587, ASV_944, ASV_83, ASV_92 |
| Root | ASV_70 | Pseudomonadota | Gammaproteobacteria | Nevskiales | Steroidobacteraceae | Povalibacter | ASV_1826, ASV_205, ASV_253, ASV_260, ASV_266, ASV_270, ASV_285, ASV_484, ASV_546, ASV_644, ASV_86, ASV_966 |

**Table S5** (continued)

| Compart-ment | ASV | Phylum | Class | Order | Family | Genus | Connections (within-kingdom) |
|---|---|---|---|---|---|---|---|
| Root | ASV_712 | Pseudomonadota | Gammaproteobacteria | Nevskiales | Steroidobacteraceae | Povalibacter | ASV_109, ASV_124, ASV_150, ASV_153, ASV_27, ASV_436, ASV_1237, ASV_1436, ASV_1587, ASV_715 |
| Root | ASV_77 | Pseudomonadota | Gammaproteobacteria | unclassified | unclassified | Acidibacter | ASV_1115, ASV_125, ASV_1334, ASV_164, ASV_171, ASV_191, ASV_2100, ASV_310, ASV_342, ASV_397, ASV_417, ASV_532, ASV_800, ASV_84 |
| Root | ASV_78 | Acidobacteria | Acidobacteriia | Acidobacteriales | Acidobacteriaceae | Koribacter | ASV_123, ASV_177, ASV_216, ASV_276, ASV_294, ASV_346, ASV_429, ASV_550, ASV_745, ASV_920 |
| Root | ASV_800 | Actinobacteria | Rubrobacteridae | Thermoleophilales | Thermoleophilaceae | Thermoleophilum | ASV_101, ASV_123, ASV_149, ASV_310, ASV_382, ASV_483, ASV_529, ASV_729, ASV_77, ASV_1334, ASV_1386, ASV_2100, ASV_944 |
| Root | ASV_848 | Actinobacteria | Actinobacteria | Actinomycetales | Catenulisporaceae | Catenulispora | ASV_118, ASV_164, ASV_171, ASV_331, ASV_8, ASV_84, ASV_1084 |
| Root | ASV_86 | Acidobacteria | Holophagae | Vicinamibacterales | Vicinamibacteraceae | Luteitalea | ASV_256, ASV_339, ASV_451, ASV_484, ASV_619 |
| Root | ASV_92 | Acidobacteria | Acidobacteriia | Acidobacteriales | Acidobacteriaceae | Koribacter | ASV_117, ASV_118, ASV_1237, ASV_1587, ASV_403, ASV_482, ASV_677, ASV_949, ASV_963 |
| Root | ASV_944 | Proteobacteria | Alphaproteobacteria | Rhodospirillales | Rhodospirillaceae | Inquilinus | ASV_134, ASV_154, ASV_193, ASV_223, ASV_314, ASV_429, ASV_677, ASV_800, ASV_1115, ASV_1285, ASV_1386 |
| Root | ASV_949 | Actinobacteria | Rubrobacteridae | Thermoleophilales | Thermoleophilaceae | Thermoleophilum | ASV_147, ASV_207, ASV_254, ASV_387, ASV_411, ASV_433, ASV_558, ASV_650, ASV_791, ASV_92, ASV_1115, ASV_1977 |
| Root | ASV_963 | Pseudomonadota | Alphaproteobacteria | Hyphomicrobiales | Hyphomicrobiaceae | Filomicrobium | ASV_233, ASV_270, ASV_411, ASV_595, ASV_666, ASV_92, ASV_1436 |
| Root | ASV_966 | Pseudomonadota | Alphaproteobacteria | Kordiimonadales | Kordiimonadaceae | Kordiimonas | ASV_134, ASV_191, ASV_205, ASV_292, ASV_307, ASV_366, ASV_517, ASV_546, ASV_70, ASV_799 |

**Table S6.** Fungal taxa involved in putative diffuse (keystone) cross-kingdom interactions identified from the network using an edge-wise variability estimate of 0.8 and 75th percentile threshold for degree and eigenvector centrality to define keystoneness. Assignments results from placing unknown sequences on T-BAS reference tree.

| Compartment | ASV | Phylum | Class | Order | Family | Genus | Connections (within-kingdom) |
|---|---|---|---|---|---|---|---|
| Leaf | ASV_122 | Ascomycota | Eurotiomycetes | Chaetothyriales | Chaetothyriaceae | Aphanophora | ASV_833, ASV_531, ASV_1519, ASV_312 |
| Leaf | ASV_278 | Ascomycota | Dothideomycetes | Pleosporales | Didymellaceae | Cumuliphoma | ASV_806, ASV_646, ASV_136, ASV_225, ASV_526 |
| Leaf | ASV_1026 | Ascomycota | Dothideomycetes | Pleosporales | Pleosporaceae | Curvularia | ASV_3134, ASV_1519, ASV_639, ASV_526, ASV_525 |
| Leaf | ASV_135 | Ascomycota | Eurotiomycetes | Chaetothyriales | Trichomeriaceae | Knufia | ASV_39, ASV_55, ASV_362, ASV_272, ASV_531 |
| Leaf | ASV_639 | Ascomycota | Dothideomycetes | Pleosporales | Didymosphaeriaceae | Montagnula | ASV_1026, ASV_373, ASV_646, ASV_3373, ASV_1167, ASV_2180, ASV_796 |
| Leaf | ASV_1519 | Basidiomycota | Tremellomycetes | Tremellales | Rhynchogastremataceae | Papiliotrema | ASV_646, ASV_473, ASV_1026, ASV_122 |
| Leaf | ASV_272 | Ascomycota | Dothideomycetes | Capnodiales | unclassified | Paradevriesia | ASV_66, ASV_135, ASV_1345, ASV_717, ASV_806, ASV_236 |
| Leaf | ASV_646 | Ascomycota | Dothideomycetes | Pleosporales | Phaeosphaeriaceae | Phaeosphaeria | ASV_1519, ASV_182, ASV_278, ASV_639 |
| Leaf | ASV_526 | Basidiomycota | Spiculogloeomycetes | Spiculogloeales | Spiculogloeaceae | Phyllozyma | ASV_278, ASV_55, ASV_208, ASV_1026, ASV_1962 |
| Root | ASV_324 | Mucoromycota | Glomeromycetes | Diversisporales | Glomeraceae | Paraglomus | ASV_258, ASV_297, ASV_790 |
| Root | ASV_314 | Mucoromycota | Glomeromycetes | Diversisporales | Glomeraceae | Paraglomus | ASV_209 |
| Root | ASV_798 | Basidiomycota | Ustilaginomycetes | Ustilaginales | Ustilaginaceae | Shivasia | ASV_241, ASV_844 |

**Table S7.** Putative ecological roles for all taxa identified from the network using an edge-wise variability estimate of 0.8 and 75th percentile threshold for degree and eigenvector centrality to define keystoneness. Relying on assignments from placing unknown sequences on T-BAS reference tree.

| Compartment | Kingdom | ASV | Ecological Role | Reference |
|---|---|---|---|---|
| Leaf | Bacteria | ASV_112 | NA | |
| Leaf | Bacteria | ASV_1145 | NA | |
| Leaf | Bacteria | ASV_1151 | NA | |
| Leaf | Bacteria | ASV_11551 | NA | |
| Leaf | Bacteria | ASV_116 | plant growth promoting | Cordovez et al., 2018 |
| Leaf | Bacteria | ASV_12 | NA | |
| Leaf | Bacteria | ASV_121 | plant growth promoting | Holochov et al., 2020 |
| Leaf | Bacteria | ASV_1246 | NA | |
| Leaf | Bacteria | ASV_13 | biodegradation | Sharma et al., 2021 |
| Leaf | Bacteria | ASV_1306 | plant growth promoting | Asaf et al., 2020 |
| Leaf | Bacteria | ASV_136 | NA | |
| Leaf | Bacteria | ASV_1454 | NA | |
| Leaf | Bacteria | ASV_166 | NA | |
| Leaf | Bacteria | ASV_1729 | nitrogen fixing, carbon decomposition | Xu et al., 2024; Zhang et al., 2024 |
| Leaf | Bacteria | ASV_1751 | plant growth promoting | Asaf et al., 2020 |
| Leaf | Bacteria | ASV_189 | NA | |
| Leaf | Bacteria | ASV_199 | halophyte, endophyte, antifungal | Chung et al., 2016 |
| Leaf | Bacteria | ASV_2009 | NA | |
| Leaf | Bacteria | ASV_2248 | plant growth promoting | Asaf et al., 2020 |
| Leaf | Bacteria | ASV_237 | plant growth promoting | Asaf et al., 2020 |
| Leaf | Bacteria | ASV_2370 | NA | |
| Leaf | Bacteria | ASV_2996 | NA | |
| Leaf | Bacteria | ASV_3051 | NA | |
| Leaf | Bacteria | ASV_3132 | pathogen | Ryan et al., 2011 |
| Leaf | Bacteria | ASV_320 | plant growth promoting | Asaf et al., 2020 |
| Leaf | Bacteria | ASV_3230 | sulfide oxidizing | Reeder et al., 2022 |
| Leaf | Bacteria | ASV_3422 | plant growth promoting | Asaf et al., 2020 |
| Leaf | Bacteria | ASV_3885 | NA | |
| Leaf | Bacteria | ASV_4005 | biodegradation; plant growth promoting under high metal | Fan et al., 2022; Ma et al., 2023 |
| Leaf | Bacteria | ASV_4016 | NA | |
| Leaf | Bacteria | ASV_4017 | NA | |
| Leaf | Bacteria | ASV_4059 | biodegradation; plant growth promoting under high metal | Fan et al., 2022; Ma et al., 2023 |
| Leaf | Bacteria | ASV_4236 | phytoremediation | Wu et al., 2022 |
| Leaf | Bacteria | ASV_443 | endophyte, plant growth promoting (under high salinity) | Bouam et al., 2018 |
| Leaf | Bacteria | ASV_45 | pathogen resistance, plant growth promoting | Chase et al., 2016 |
| Leaf | Bacteria | ASV_4546 | biodegradation; plant growth promoting under high metal | Fan et al., 2022; Ma et al., 2023 |
| Leaf | Bacteria | ASV_4622 | plant growth promoting, nitrogen fixing | Patel and Patel, 2023 |
| Leaf | Bacteria | ASV_4693 | NA | |
| Leaf | Bacteria | ASV_470 | plant growth promoting | Madhaiyan et al., 2015 |
| Leaf | Bacteria | ASV_474 | nitrogen cycling | Liu et al., 2023 |
| Leaf | Bacteria | ASV_4860 | biocontrol | Liu et al., 2018 |
| Leaf | Bacteria | ASV_4880 | osmotic stress tolerance, ammonia oxidizing | Lengrand et al., 2024 |
| Leaf | Bacteria | ASV_571 | plant growth promoting | Holochov et al., 2020 |
| Leaf | Bacteria | ASV_5911 | nutrient cycling | Li et al. 2023 |
| Leaf | Bacteria | ASV_597 | NA | |
| Leaf | Bacteria | ASV_599 | NA | |
| Leaf | Bacteria | ASV_6 | plant growth promoting, root nodule endophyte | Zheng et al., 2023 |
| Leaf | Bacteria | ASV_626 | NA | |
| Leaf | Bacteria | ASV_6310 | NA | |
| Leaf | Bacteria | ASV_6358 | nitrogen fixing | Masuda et al., 2020 |
| Leaf | Bacteria | ASV_6715 | plant growth promoting | Datta and Nag, 2022 |
| Leaf | Bacteria | ASV_682 | NA | |

**Table S7** (continued)

| Compartment | Kingdom | ASV | Ecological Role | Reference |
|---|---|---|---|---|
| Leaf | Bacteria | ASV_689 | NA | |
| Leaf | Bacteria | ASV_743 | plant growth promoting, disease suppression | Berg et al., 2021; Zhang et al., 2019 |
| Leaf | Bacteria | ASV_747 | plant growth promoting | Madhaiyan et al., 2015 |
| Leaf | Bacteria | ASV_768 | NA | |
| Leaf | Bacteria | ASV_842 | plant degradation, biocontrol | Huang et al., 2019 |
| Leaf | Bacteria | ASV_8842 | biocontrol | Shi et al., 2024 |
| Leaf | Bacteria | ASV_9 | NA | |
| Leaf | Bacteria | ASV_929 | predator | Wang et al., 2011 |
| Leaf | Bacteria | ASV_9928 | NA | |
| Leaf | Bacteria | ASV_993 | plant growth promoting | Kumar et al., 2019 |
| Leaf | Fungi | ASV_102 | endophyte | Li et al., 2022 |
| Leaf | Fungi | ASV_1026 | pathogen | Khiralla et al., 2019 |
| Leaf | Fungi | ASV_1044 | leaf yeast epiphyte | Chai et al., 2023 |
| Leaf | Fungi | ASV_115 | endophyte | Rush and Aime, 2013 |
| Leaf | Fungi | ASV_122 | endophyte | Yang et al., 2022 |
| Leaf | Fungi | ASV_130 | yeast | Sampaio et al., 2002 |
| Leaf | Fungi | ASV_135 | rock inhabiting, drought tolerance | Gerrits et al., 2020; Yao et al., 2022 |
| Leaf | Fungi | ASV_138 | yeast | Schoutteten et al., 2025 |
| Leaf | Fungi | ASV_144 | NA | |
| Leaf | Fungi | ASV_1519 | saprophytic | Palmier et al., 2021 |
| Leaf | Fungi | ASV_17 | pathogen | Ali-Arous et al., 2023 |
| Leaf | Fungi | ASV_177 | endophyte, pathogen | Luo et al., 2017; Sodhi et al., 2023 |
| Leaf | Fungi | ASV_18 | pathogen | Luo et al., 2017; Sodhi et al., 2023 |
| Leaf | Fungi | ASV_1833 | NA | |
| Leaf | Fungi | ASV_1962 | endophyte | Rush and Aime, 2013 |
| Leaf | Fungi | ASV_2 | pathogen | Gavrilova et al., 2020 |
| Leaf | Fungi | ASV_20 | NA | |
| Leaf | Fungi | ASV_225 | NA | |
| Leaf | Fungi | ASV_24 | pathogen, potentially endophyte | Liu et al., 2019 |
| Leaf | Fungi | ASV_256 | pathogen | Shoemaker et al., 1989; El-Damerdash, 2018 |
| Leaf | Fungi | ASV_259 | endophyte, pathogen | Ali-Arous et al., 2023; Impullitti and Malvick, 2014 |
| Leaf | Fungi | ASV_2659 | NA | |
| Leaf | Fungi | ASV_272 | plant and rock living | Crous et al., 2019 |
| Leaf | Fungi | ASV_3373 | yeast, biodegradation | Al-Tohamy et al., 2020 |
| Leaf | Fungi | ASV_4 | NA | |
| Leaf | Fungi | ASV_406 | yeast | Sampaio et al., 2002 |
| Leaf | Fungi | ASV_458 | yeast, mycoparasite | Sampaio et al., 2002 |
| Leaf | Fungi | ASV_473 | endophyte, pathogen | Luo et al., 2017; Sodhi et al., 2023 |
| Leaf | Fungi | ASV_526 | yeast, endophyte | Li et al., 2020 |
| Leaf | Fungi | ASV_53 | NA | |
| Leaf | Fungi | ASV_531 | NA | |
| Leaf | Fungi | ASV_55 | epiphytic yeast, biocontrol | Li et al., 2021 |
| Leaf | Fungi | ASV_58 | lichenicolous | Monteiro et al., 2021 |
| Leaf | Fungi | ASV_639 | saprophytic | Wanasinghe et al., 2024 |
| Leaf | Fungi | ASV_646 | pathogen | Shoemaker et al., 1989; El-Damerdash, 2018 |
| Leaf | Fungi | ASV_66 | pathogen | Ali-Arous et al., 2023 |
| Leaf | Fungi | ASV_7 | pathogen | Gomez-Zapata et al., 2024 |
| Leaf | Fungi | ASV_744 | NA | |
| Leaf | Fungi | ASV_856 | endophyte | Rush and Aime, 2013 |
| Leaf | Fungi | ASV_916 | pathogen | Lutz et al., 2018 |
| Root | Bacteria | ASV_1018 | nitrogen, lignin degrading | Tarquinio et al., 2021; Tom et al., 2022 |
| Root | Bacteria | ASV_103 | carbon cycling | Bill et al., 2021 |
| Root | Bacteria | ASV_1042 | plant growth promoting | Zhou et al., 2022 |
| Root | Bacteria | ASV_1078 | Fe and Mn reducing | Boutsika et al., 2024 |
| Root | Bacteria | ASV_1084 | plant growth promoting, nutrient cycling | Kalam et al., 2020 |
| Root | Bacteria | ASV_109 | biocontrol | Lazcano et al., 2021 |
| Root | Bacteria | ASV_1115 | Fe reducing, wood degradation | Gavrilov et al., 2019 |
| Root | Bacteria | ASV_1139 | NA | |
| Root | Bacteria | ASV_118 | acidophilic | Eo et al., 2021 |
| Root | Bacteria | ASV_1237 | NA | |
| Root | Bacteria | ASV_124 | plant growth promoting, nutrient cycling | Liu et al., 2022 |
| Root | Bacteria | ASV_1292 | plant growth promoting | Bouam et al., 2018 |

**Table S7** (continued)

| Compartment | Kingdom | ASV | Ecological Role | Reference |
|---|---|---|---|---|
| Root | Bacteria | ASV_1334 | NA | |
| Root | Bacteria | ASV_1369 | biocontrol | Bhat et al., 2021 |
| Root | Bacteria | ASV_1395 | NA | |
| Root | Bacteria | ASV_143 | sulphur oxidizing | Tsallagov et al., 2019 |
| Root | Bacteria | ASV_1436 | NA | |
| Root | Bacteria | ASV_147 | carbon cycling | Bill et al., 2021 |
| Root | Bacteria | ASV_149 | acidophilic | Rapoport et al., 2020 |
| Root | Bacteria | ASV_150 | NA | |
| Root | Bacteria | ASV_1587 | biocontrol | B. Yang et al., 2023 |
| Root | Bacteria | ASV_164 | endophyte, plant growth promoting | Gangwar et al., 2012 |
| Root | Bacteria | ASV_165 | plant growth promoting, carbon cycling | Darriaut et al., 2024 |
| Root | Bacteria | ASV_171 | acidophilic | Chen et al., 2020 |
| Root | Bacteria | ASV_191 | plant growth promoting | Bouam et al., 2018 |
| Root | Bacteria | ASV_193 | acidophilic | Eo et al., 2021 |
| Root | Bacteria | ASV_2 | rhizobia, nitrogen fixing | Jiang et al., 2024 |
| Root | Bacteria | ASV_200 | biodegradation | Poria et al., 2022 |
| Root | Bacteria | ASV_205 | nitrogen fixing | Benson et al., 1993 |
| Root | Bacteria | ASV_207 | plant growth promoting | Li et al. 2023 |
| Root | Bacteria | ASV_21 | plant growth promoting, nitrogen fixing | Figueiredo et al., 2020 |
| Root | Bacteria | ASV_215 | biocontrol | Bhat et al., 2021 |
| Root | Bacteria | ASV_216 | endophyte | Ricardo et al., 2020 |
| Root | Bacteria | ASV_24 | carbon cycling | Bill et al., 2021 |
| Root | Bacteria | ASV_241 | biodegradation | Sharma et al., 2021 |
| Root | Bacteria | ASV_245 | plant growth promoting, nutrient cycling | Kalam et al., 2020 |
| Root | Bacteria | ASV_249 | nitrogen mineralization | Tamas et al., 2012 |
| Root | Bacteria | ASV_254 | NA | |
| Root | Bacteria | ASV_26 | carbon cycling | Bill et al., 2021 |
| Root | Bacteria | ASV_264 | pathogen | Thompson et al., 2018 |
| Root | Bacteria | ASV_269 | plant growth promoting | Lin et al., 2022 |
| Root | Bacteria | ASV_28 | vitamin producer | Astafyeva et al., 2022 |
| Root | Bacteria | ASV_292 | NA | |
| Root | Bacteria | ASV_306 | plant growth promoting | Lin et al., 2022 |
| Root | Bacteria | ASV_307 | NA | |
| Root | Bacteria | ASV_31 | rhizobia, nitrogen fixing | Jiang et al., 2024 |
| Root | Bacteria | ASV_314 | water stress tolerance, pH | Sun et al., 2022; L. Wang et al., 2023 |
| Root | Bacteria | ASV_32 | NA | |
| Root | Bacteria | ASV_322 | carbon cycling | Gillan et al., 2023 |
| Root | Bacteria | ASV_330 | nitrogen, lignin degrading | Tarquinio et al., 2021; Tom et al., 2022 |
| Root | Bacteria | ASV_331 | plant growth promoting, carbon cycling | Darriaut et al., 2024 |
| Root | Bacteria | ASV_35 | plant growth promoting | Preston, 2004 |
| Root | Bacteria | ASV_366 | NA | |
| Root | Bacteria | ASV_371 | NA | |
| Root | Bacteria | ASV_377 | NA | |
| Root | Bacteria | ASV_38 | plant growth promoting | Olanrewaju et al., 2019 |
| Root | Bacteria | ASV_390 | NA | |
| Root | Bacteria | ASV_394 | biocontrol | Lazcano et al., 2021 |
| Root | Bacteria | ASV_397 | biocontrol, endophyte | Naama et al., 2020 |
| Root | Bacteria | ASV_402 | NA | |
| Root | Bacteria | ASV_407 | NA | |
| Root | Bacteria | ASV_41 | nutrient cycling | Jia et al., 2022 |
| Root | Bacteria | ASV_411 | NA | |
| Root | Bacteria | ASV_414 | plant growth promoting, biocontrol | D. Wang et al., 2023 |
| Root | Bacteria | ASV_433 | nutrient cycling | Jia et al., 2022 |
| Root | Bacteria | ASV_442 | nutrient cycling | Jia et al., 2022 |
| Root | Bacteria | ASV_469 | plant growth promoting | Lin et al., 2022 |
| Root | Bacteria | ASV_503 | AMF endobacteria | Jargeat et al., 2004 |
| Root | Bacteria | ASV_510 | plant growth promoting | Rat et al., 2021 |
| Root | Bacteria | ASV_512 | nutrient cycling, biocontrol, plant growth promoting | Y. Wang et al., 2023 |

**Table S7** (continued)

| Compartment | Kingdom | ASV | Ecological Role | Reference |
|---|---|---|---|---|
| Root | Bacteria | ASV_513 | biocontrol | Lazcano et al., 2021 |
| Root | Bacteria | ASV_532 | NA | |
| Root | Bacteria | ASV_539 | nitrogen fixing | Liao et al., 2022 |
| Root | Bacteria | ASV_551 | plant growth promoting | Bouam et al., 2018 |
| Root | Bacteria | ASV_558 | nutrient cycling | Li et al. 2023 |
| Root | Bacteria | ASV_570 | acidophilic | Chen et al., 2020 |
| Root | Bacteria | ASV_595 | NA | |
| Root | Bacteria | ASV_61 | biocontrol | Piao et al., 2017 |
| Root | Bacteria | ASV_558 | nutrient cycling | Li et al. 2023 |
| Root | Bacteria | ASV_570 | acidophilic | Chen et al., 2020 |
| Root | Bacteria | ASV_595 | NA | |
| Root | Bacteria | ASV_61 | biocontrol | Piao et al., 2017 |
| Root | Bacteria | ASV_625 | biocontrol | Lazcano et al., 2021 |
| Root | Bacteria | ASV_630 | plant growth promoting | Lin et al., 2022 |
| Root | Bacteria | ASV_644 | nitrogen, lignin degrading | Tarquinio et al., 2021; Tom et al., 2022 |
| Root | Bacteria | ASV_650 | plant growth promoting | Bouam et al., 2018 |
| Root | Bacteria | ASV_663 | biocontrol | Lazcano et al., 2021 |
| Root | Bacteria | ASV_666 | sulphur oxidizing | Tsallagov et al., 2019 |
| Root | Bacteria | ASV_677 | beneficial | Aguirre-von-Wobeser et al., 2018 |
| Root | Bacteria | ASV_69 | NA | |
| Root | Bacteria | ASV_699 | nitrogen fixing | Xie et al., 2023 |
| Root | Bacteria | ASV_70 | saprophytic | Lazcano et al., 2021 |
| Root | Bacteria | ASV_712 | saprophytic | Lazcano et al., 2021 |
| Root | Bacteria | ASV_77 | nutrient cycling | Li et al. 2023 |
| Root | Bacteria | ASV_78 | acidophilic | Eo et al., 2021 |
| Root | Bacteria | ASV_800 | NA | |
| Root | Bacteria | ASV_83 | acidophilic | Eo et al., 2021 |
| Root | Bacteria | ASV_84 | nutrient cycling | Li et al. 2023 |
| Root | Bacteria | ASV_848 | plant-associated | Chen et al., 2020 |
| Root | Bacteria | ASV_86 | chemo-organotrophic mesophile | Vieira et al., 2017 |
| Root | Bacteria | ASV_887 | plant growth promoting | Asaf et al., 2020 |
| Root | Bacteria | ASV_89 | plant growth promoting | Olanrewaju et al., 2019 |
| Root | Bacteria | ASV_92 | acidophilic | Eo et al., 2021 |
| Root | Bacteria | ASV_933 | NA | |
| Root | Bacteria | ASV_944 | plant growth promoting | Rat et al., 2021 |
| Root | Bacteria | ASV_949 | NA | |
| Root | Bacteria | ASV_96 | NA | |
| Root | Bacteria | ASV_963 | nitrogen, lignin degrading | Tarquinio et al., 2021; Tom et al., 2022 |
| Root | Bacteria | ASV_966 | halotolerant | Yang et al., 2016 |
| Root | Fungi | ASV_107 | endophyte | D. Yang et al., 2023 |
| Root | Fungi | ASV_113 | NA | |
| Root | Fungi | ASV_12 | NA | |
| Root | Fungi | ASV_134 | AMF | Redecker et al., 2013 |
| Root | Fungi | ASV_140 | NA | |
| Root | Fungi | ASV_150 | NA | |
| Root | Fungi | ASV_161 | NA | |
| Root | Fungi | ASV_169 | rock inhabiting | Gerrits et al., 2020; Yao et al., 2022 |
| Root | Fungi | ASV_1787 | epiphytic yeast, biocontrol | Khreireddine et al., 2018 |
| Root | Fungi | ASV_179 | NA | |
| Root | Fungi | ASV_209 | AMF | Redecker et al., 2013 |
| Root | Fungi | ASV_231 | decomposer, pathogen | Badali et al., 2008 |
| Root | Fungi | ASV_240 | endophyte | D. Yang et al., 2023 |
| Root | Fungi | ASV_248 | pathogen, endophyte | Shetty et al., 2016 |
| Root | Fungi | ASV_254 | AMF | Redecker et al., 2013 |
| Root | Fungi | ASV_26 | NA | |
| Root | Fungi | ASV_282 | NA | |
| Root | Fungi | ASV_291 | rock inhabiting | Gerrits et al., 2020; Yao et al., 2022 |
| Root | Fungi | ASV_314 | AMF | Redecker et al., 2013 |
| Root | Fungi | ASV_319 | AMF | Redecker et al., 2013 |
| Root | Fungi | ASV_324 | AMF | Redecker et al., 2013 |
| Root | Fungi | ASV_334 | NA | |

**Table S7** (continued)

| Compartment | Kingdom | ASV | Ecological Role | Reference |
|---|---|---|---|---|
| Root | Fungi | ASV_370 | NA | |
| Root | Fungi | ASV_375 | NA | |
| Root | Fungi | ASV_38 | pathogen, endophyte | Shetty et al., 2016 |
| Root | Fungi | ASV_416 | NA | |
| Root | Fungi | ASV_443 | NA | |
| Root | Fungi | ASV_5 | pathogen | Lutz et al., 2012 |
| Root | Fungi | ASV_56 | yeast | Tian et al., 2021; Yurkov et al., 2021 |
| Root | Fungi | ASV_567 | AMF | Redecker et al., 2013 |
| Root | Fungi | ASV_57 | pathogen | Gryndler et al., 2009 |
| Root | Fungi | ASV_654 | NA | |
| Root | Fungi | ASV_790 | AMF | Redecker et al., 2013 |
| Root | Fungi | ASV_798 | pathogen | Lutz et al., 2012 |
| Root | Fungi | ASV_827 | NA | |

**Table S8.** Independent variables included in the environmental and spatial matrices for variance partitioning analysis.

| Variable | Measurement Scale | Unit |
|---|---|---|
| *Environmental* | | |
| pH | plant | NA |
| soil moisture | plant | g/g |
| P | plant | ug/g |
| K | plant | ug/g |
| TIN | plant | ug/g |
| C | plant | % |
| MBC | plant | ug/g |
| Sand | plant | % |
| Clay | plant | % |
| plant height | plant | m |
| plant basal area | plant | $m^2$ |
| MAT | site | C |
| MAP | site | m |
| *Spatial* | | |
| latitude | plant | DD |
| longitude | plant | DD |
| plot area | site | $m^2$ |

**Table S9.** Results from variance partitioning analysis for (A) leaf and (B) root communities. Explanatory matrices include standardized environmental and spatial variables (Table S7), and standardized ASV count of the directly interacting taxa from the other kingdom, defined from the networks with parameters discussed in the main text (edge-wise variability estimate of 0.8). Response ASV table is Hellinger-transformed. Bold values are significant ($P < 0.05$), determined from redundancy analysis (RDA).

#### A. Leaf community variance partitioning analysis

| **Target community:** | **Bacteria** | | | | **Fungi** | | | |
|---|---|---|---|---|---|---|---|---|
| **Cross-kingdom interaction type:** | S + E (only) | Direct | Diffuse - keystone | Diffuse - alpha | S + E (only) | Direct | Diffuse - keystone | Diffuse - alpha |
| **Spatial (S)** | **0.050** | **0.027** | **0.045** | 0.050 | **0.057** | **0.028** | **0.053** | 0.051 |
| **Environmental (E)** | **0.222** | **0.078** | **0.153** | 0.218 | **0.275** | **0.123** | **0.211** | 0.226 |
| **Biotic (B)** | NA | **0.114** | **0.056** | **-0.001** | **NA** | **0.104** | 0.026 | 0.015 |
| **E + S** | 0.054 | 0.021 | 0.048 | **0.053** | 0.024 | 0.028 | 0.023 | **0.033** |
| **E + B** | NA | 0.144 | 0.069 | **0.004** | NA | 0.152 | 0.064 | **0.049** |
| **S + B** | NA | 0.023 | 0.005 | **0.000** | NA | 0.029 | 0.004 | **0.006** |
| **E + S + B** | NA | 0.033 | 0.006 | **0.001** | NA | -0.004 | 0.001 | **-0.009** |
| **Residuals** | 0.674 | 0.560 | 0.618 | **0.675** | 0.645 | 0.540 | 0.619 | **0.630** |

#### B. Root community variance partitioning analysis

| **Target community:** | **Bacteria** | | | | **Fungi** | | | |
|---|---|---|---|---|---|---|---|---|
| **Cross-kingdom interaction type:** | S + E (only) | Direct | Diffuse - keystone | Diffuse - alpha | S + E (only) | Direct | Diffuse - keystone | Diffuse - alpha |
| **Spatial** | **0.027** | **0.025** | **0.029** | **0.028** | **0.033** | **0.025** | **0.039** | **0.033** |
| **Environmental** | **0.219** | **0.139** | **0.217** | **0.186** | **0.186** | **0.099** | **0.080** | **0.162** |
| **Biotic** | NA | **0.046** | **0.007** | **0.006** | **NA** | **0.040** | 0.084 | 0.003 |
| **S + E** | 0.109 | 0.075 | 0.109 | 0.105 | 0.043 | 0.027 | 0.019 | 0.050 |
| **E + B** | NA | 0.080 | 0.003 | 0.033 | NA | 0.087 | 0.107 | 0.024 |
| **S + B** | NA | 0.002 | -0.002 | -0.001 | NA | 0.008 | -0.006 | 0.000 |
| **S + E + B** | NA | 0.034 | 0.000 | 0.004 | NA | 0.016 | 0.024 | -0.007 |
| **Residuals** | 0.640 | 0.599 | 0.638 | 0.639 | 0.738 | 0.698 | 0.654 | 0.735 |

**Table S10.** Results of variance partitioning analyses for leaf communities, repeated for the different network parameters tested to compare sensitivity of outcomes. Dependent variable is the Hellinger-transformed (A) fungal or (B) bacterial leaf community. Explanatory matrices include standardized environmental and spatial variables (Table S7), and a matrix to represent biotic interactions composed of the standardized ASV count of the directly interacting taxa from the other kingdom. The ASVs in the biotic matrix are identified from each network resulting from using different edge-wise variability estimates (0.8, 0.9, 0.95). P-values for each variance partitioning, or redundancy analysis (RDA), model for leaf (C) fungal or (D) bacterial community.

| | **A. Fungi as target community** | | | | **B. Bacteria as target community** | | | |
|---|---|---|---|---|---|---|---|---|
| | S + E (only) | Edge 0.8 | Edge 0.9 | Edge 0.95 | S + E (only) | Edge 0.8 | Edge 0.9 | Edge 0.95 |
| Spatial (S) | 0.057 | 0.028 | 0.038 | 0.047 | 0.050 | 0.027 | 0.043 | 0.042 |
| Environmental (E) | 0.275 | 0.123 | 0.157 | 0.181 | 0.222 | 0.078 | 0.104 | 0.112 |
| Biotic (B) | | 0.104 | 0.052 | 0.033 | | 0.114 | 0.063 | 0.045 |
| E + S | 0.024 | 0.028 | 0.025 | 0.028 | 0.054 | 0.021 | 0.042 | 0.038 |
| E + B | | 0.152 | 0.118 | 0.094 | | 0.144 | 0.118 | 0.110 |
| S + B | | 0.029 | 0.019 | 0.010 | | 0.023 | 0.007 | 0.008 |
| E + S + B | | -0.004 | -0.001 | -0.004 | | 0.033 | 0.012 | 0.015 |
| Residuals | 0.645 | 0.540 | 0.593 | 0.612 | 0.674 | 0.560 | 0.611 | 0.629 |

| | **C. *P*-values from RDA for fungi as target community** | | | | **D. *P*-vales from RDA for bacteria as target community** | | | |
|---|---|---|---|---|---|---|---|---|
| | S + E (only) | Edge 0.8 | Edge 0.9 | Edge 0.95 | S + E (only) | Edge 0.8 | Edge 0.9 | Edge 0.95 |
| Spatial | 0.001 | 0.005 | 0.001 | 0.001 | 0.001 | 0.001 | 0.001 | 0.001 |
| Environmental | 0.001 | 0.001 | 0.001 | 0.001 | 0.001 | 0.001 | 0.001 | 0.001 |
| Biotic | | 0.001 | 0.001 | 0.003 | | 0.001 | 0.001 | 0.001 |
| global model | 0.001 | 0.001 | 0.001 | 0.001 | 0.001 | 0.001 | 0.001 | 0.001 |

**Table S11.** Results of variance partitioning analyses for leaf communities, repeated for the different network and keystone parameters tested to compare sensitivity of outcomes. Dependent variable is the Hellinger-transformed (A) fungal or (B) bacterial leaf community. Explanatory matrices include standardized environmental and spatial variables (Table S7), and biotic interactions represented by the standardized ASV count of the diffusely interacting taxa from the other kingdom identified at different thresholds for each network edge-wise variability estimate (0.8, 0.9, 0.95). Diffuse interactions are keystone taxa defined as nodes with degree and eigenvector centrality values in either the 75th, 80th, or 90th percentile. P-values for each variance partitioning, or redundancy analysis (RDA), model for leaf (C) fungal or (D) bacterial community.

| | **A. Fungi as target community** | | | | | | | | | | **B. Bacteria as target community** | | | | | | | | | |
|---|---|---|---|---|---|---|---|---|---|---|---|---|---|---|---|---|---|---|---|---|
| | S + E (only) | Edge 0.8 | | | Edge 0.9 | | | Edge 0.95 | | | S + E (only) | Edge 0.8 | | | Edge 0.9 | | | Edge 0.95 | | |
| Keystone Percentile | | 75th | 80th | 90th | 75th | 80th | 90th | 75th | 80th | 90th | | 75th | 80th | 90th | 75th | 80th | 90th | 75th | 80th | 90th |
| Spatial (S) | 0.057 | 0.053 | 0.054 | 0.056 | 0.050 | 0.046 | 0.053 | 0.055 | 0.054 | 0.053 | 0.050 | 0.045 | 0.048 | | 0.035 | 0.038 | 0.044 | 0.042 | 0.043 | 0.043 |
| Environmental (E) | 0.275 | 0.211 | 0.214 | 0.221 | 0.217 | 0.215 | 0.238 | 0.209 | 0.232 | 0.237 | 0.222 | 0.153 | 0.202 | | 0.121 | 0.146 | 0.200 | 0.138 | 0.144 | 0.179 |
| Biotic (B) | | 0.026 | 0.037 | 0.005 | 0.031 | 0.029 | 0.001 | 0.006 | -0.003 | -0.002 | | 0.056 | 0.021 | | 0.028 | 0.025 | 0.017 | 0.039 | 0.031 | 0.020 |
| E + S | 0.024 | 0.023 | 0.027 | 0.023 | 0.025 | 0.028 | 0.027 | 0.013 | 0.013 | 0.022 | 0.054 | 0.048 | 0.054 | | 0.053 | 0.052 | 0.054 | 0.055 | 0.053 | 0.053 |
| E + B | | 0.064 | 0.061 | 0.054 | 0.058 | 0.060 | 0.037 | 0.066 | 0.043 | 0.038 | | 0.069 | 0.020 | | 0.101 | 0.077 | 0.022 | 0.084 | 0.078 | 0.043 |
| S + B | | 0.004 | 0.003 | 0.001 | 0.007 | 0.011 | 0.003 | 0.002 | 0.003 | 0.004 | | 0.005 | 0.002 | | 0.015 | 0.012 | 0.006 | 0.008 | 0.007 | 0.007 |
| E + S + B | | 0.001 | -0.003 | 0.001 | -0.001 | -0.005 | -0.003 | 0.011 | 0.010 | 0.002 | | 0.006 | 0.000 | | 0.001 | 0.002 | 0.000 | -0.001 | 0.001 | 0.001 |
| Residuals | 0.645 | 0.619 | 0.608 | 0.639 | 0.613 | 0.615 | 0.644 | 0.638 | 0.647 | 0.646 | 0.674 | 0.618 | 0.653 | | 0.646 | 0.649 | 0.657 | 0.635 | 0.643 | 0.654 |

| | **C. *P*-values from RDA for fungi as target community** | | | | | | | | | | **D. *P*-vales from RDA for bacteria as target community** | | | | | | | | | |
|---|---|---|---|---|---|---|---|---|---|---|---|---|---|---|---|---|---|---|---|---|
| | S + E (only) | Edge 0.8 | | | Edge 0.9 | | | Edge 0.95 | | | S + E (only) | Edge 0.8 | | | Edge 0.9 | | | Edge 0.95 | | |
| Keystone Percentile | | 75th | 80th | 90th | 75th | 80th | 90th | 75th | 80th | 90th | | 75th | 80th | 90th | 75th | 80th | 90th | 75th | 80th | 90th |
| Spatial | 0.001 | 0.001 | 0.001 | 0.001 | 0.001 | 0.001 | 0.001 | 0.001 | 0.001 | 0.001 | 0.001 | 0.001 | 0.001 | | 0.001 | 0.001 | 0.001 | 0.001 | 0.001 | 0.001 |
| Environmental | 0.001 | 0.001 | 0.001 | 0.001 | 0.001 | 0.001 | 0.001 | 0.001 | 0.001 | 0.001 | 0.001 | 0.001 | 0.001 | | 0.001 | 0.001 | 0.001 | 0.001 | 0.001 | 0.001 |
| Biotic | | 0.077 | 0.015 | 0.304 | 0.031 | 0.030 | 0.456 | 0.305 | 0.593 | 0.564 | | 0.001 | 0.001 | | 0.004 | 0.004 | 0.004 | 0.001 | 0.001 | 0.002 |
| global model | 0.001 | 0.001 | 0.001 | 0.001 | 0.001 | 0.001 | 0.001 | 0.001 | 0.001 | 0.001 | 0.001 | 0.001 | 0.001 | | 0.001 | 0.001 | 0.001 | 0.001 | 0.001 | 0.001 |

**Table S12.** Results of variance partitioning analyses for leaf communities, repeated for the different network parameters tested to compare sensitivity of outcomes. Dependent variable is the Hellinger-transformed (A) fungal or (B) bacterial leaf community. Explanatory tables include standardized environmental and spatial variables (Table S7), and biotic interactions represented as the alpha diversity of the other kingdom (e.g. for fungi as the dependent matrix, alpha diversity of bacteria is the independent matrix) at different thresholds for each network edge-wise variability estimate (0.8, 0.9, 0.95). P-values for each variance partitioning, or redundancy analysis (RDA), model for leaf (C) fungal or (D) bacterial community.

| | **A. Fungi as target community** | | | | **B. Bacteria as target community** | | | |
|---|---|---|---|---|---|---|---|---|
| | S + E (only) | Edge 0.8 | Edge 0.9 | Edge 0.95 | S + E (only) | Edge 0.8 | Edge 0.9 | Edge 0.95 |
| Spatial (S) | 0.057 | 0.051 | 0.051 | 0.050 | 0.050 | 0.050 | 0.050 | 0.050 |
| Environmental (E) | 0.275 | 0.226 | 0.226 | 0.227 | 0.222 | 0.218 | 0.220 | 0.220 |
| Biotic (B) | | 0.015 | 0.013 | 0.015 | | -0.001 | 0.000 | -0.002 |
| E + S | 0.024 | 0.033 | 0.034 | 0.034 | 0.054 | 0.053 | 0.055 | 0.055 |
| E + B | | 0.049 | 0.049 | 0.048 | | 0.004 | 0.002 | 0.002 |
| S + B | | 0.006 | 0.006 | 0.007 | | 0.000 | 0.000 | 0.000 |
| E + S + B | | -0.009 | -0.010 | -0.011 | | 0.001 | -0.001 | -0.001 |
| Residuals | 0.645 | 0.630 | 0.631 | 0.630 | 0.674 | 0.675 | 0.674 | 0.676 |

| | **C. P-values from RDA for fungi as target community** | | | | **D. P-vales from RDA for bacteria as target community** | | | |
|---|---|---|---|---|---|---|---|---|
| | S + E (only) | Edge 0.8 | Edge 0.9 | Edge 0.95 | S + E (only) | Edge 0.8 | Edge 0.9 | Edge 0.95 |
| Spatial | 0.001 | 0.001 | 0.001 | 0.001 | 0.001 | 0.001 | 0.001 | 0.001 |
| Environmental | 0.001 | 0.001 | 0.001 | 0.001 | 0.001 | 0.001 | 0.001 | 0.001 |
| Biotic | | 0.001 | 0.001 | 0.001 | | 0.674 | 0.519 | 0.791 |
| global model | 0.001 | 0.001 | 0.001 | 0.001 | 0.001 | 0.001 | 0.001 | 0.001 |

**Table S13.** Results of variance partitioning analyses for root communities, repeated for the different network parameters tested to compare sensitivity of outcomes. Dependent variable is the Hellinger-transformed (A) fungal or (B) bacterial root community. Explanatory matrices include standardized environmental and spatial variables (Table S7), and a matrix to represent biotic interactions composed of the standardized ASV count of the directly interacting taxa from the other kingdom. The ASVs in the biotic matrix are identified from each networks resulting from setting different edge-wise variability estimates (0.8, 0.9, 0.95). P-values for each variance partitioning, or redundancy analysis (RDA), model for leaf (C) fungal or (D) bacterial community.

| | **A. Fungi as target community** | | | | **B. Bacteria as target community** | | | |
|---|---|---|---|---|---|---|---|---|
| | S + E (only) | Edge 0.8 | Edge 0.9 | Edge 0.95 | S + E (only) | Edge 0.8 | Edge 0.9 | Edge 0.95 |
| Spatial (S) | 0.033 | 0.025 | 0.032 | 0.035 | 0.027 | 0.025 | 0.030 | 0.030 |
| Environmental (E) | 0.186 | 0.099 | 0.151 | 0.158 | 0.219 | 0.139 | 0.191 | 0.205 |
| Biotic (B) | | 0.040 | 0.026 | 0.024 | | 0.046 | 0.018 | 0.015 |
| E + S | 0.043 | 0.027 | 0.036 | 0.044 | 0.109 | 0.075 | 0.096 | 0.098 |
| E + B | | 0.087 | 0.035 | 0.028 | | 0.080 | 0.028 | 0.014 |
| S + B | | 0.008 | 0.001 | -0.002 | | 0.002 | -0.004 | -0.003 |
| E + S + B | | 0.016 | 0.007 | -0.001 | | 0.034 | 0.013 | 0.011 |
| Residuals | 0.738 | 0.698 | 0.712 | 0.714 | 0.640 | 0.599 | 0.627 | 0.630 |

| | **C. P-values from RDA for fungi as target community** | | | | **D. P-vales from RDA for bacteria as target community** | | | |
|---|---|---|---|---|---|---|---|---|
| | S + E (only) | Edge 0.8 | Edge 0.9 | Edge 0.95 | S + E (only) | Edge 0.8 | Edge 0.9 | Edge 0.95 |
| Spatial | 0.001 | 0.003 | 0.001 | 0.001 | 0.001 | 0.001 | 0.001 | 0.001 |
| Environmental | 0.001 | 0.001 | 0.001 | 0.001 | 0.001 | 0.001 | 0.001 | 0.001 |
| Biotic | | 0.010 | 0.002 | 0.002 | | 0.001 | 0.002 | 0.002 |
| global model | 0.001 | 0.001 | 0.001 | 0.001 | 0.001 | 0.001 | 0.001 | 0.001 |

**Table S14.** Results of variance partitioning analyses for root communities, repeated for the different network and keystone parameters tested to compare sensitivity of outcomes. Dependent variable is the Hellinger-transformed (A) fungal or (B) bacterial root community. Explanatory matrices include standardized environmental and spatial variables (Table S7), and biotic interactions represented by the standardized ASV count of the diffusely interacting taxa from the other kingdom identified at different thresholds for each network edge-wise variability estimate (0.8, 0.9, 0.95). Diffuse interactions are keystone taxa defined as nodes with degree and eigenvector centrality values in either the 75th, 80th, or 90th percentile. P-values for each variance partitioning, or redundancy analysis (RDA), model for leaf (C) fungal or (D) bacterial community.

**A. Fungi as target community**

| | S + E (only) | Edge 0.8 | | | Edge 0.9 | | | Edge 0.95 | | |
|---|---|---|---|---|---|---|---|---|---|---|
| Keystone Percentile | | 75th | 80th | 90th | 75th | 80th | 90th | 75th | 80th | 90th |
| Spatial (S) | 0.033 | 0.039 | 0.037 | 0.027 | 0.030 | 0.022 | 0.031 | 0.013 | 0.028 | 0.035 |
| Environmental (E) | 0.186 | 0.080 | 0.091 | 0.116 | 0.053 | 0.066 | 0.097 | 0.053 | 0.077 | 0.124 |
| Biotic (B) | | 0.084 | 0.088 | 0.035 | 0.062 | 0.033 | 0.042 | 0.023 | 0.028 | 0.031 |
| E + S | 0.043 | 0.019 | 0.028 | 0.037 | 0.027 | 0.029 | 0.034 | 0.050 | 0.029 | 0.030 |
| E + B | | 0.107 | 0.095 | 0.070 | 0.133 | 0.121 | 0.090 | 0.134 | 0.109 | 0.062 |
| S + B | | -0.006 | -0.005 | 0.006 | 0.002 | 0.011 | 0.002 | 0.019 | 0.004 | -0.002 |
| E + S + B | | 0.024 | 0.015 | 0.006 | 0.017 | 0.014 | 0.009 | -0.007 | 0.014 | 0.013 |
| Residuals | 0.738 | 0.654 | 0.650 | 0.703 | 0.676 | 0.705 | 0.696 | 0.715 | 0.710 | 0.707 |

**B. Bacteria as target community**

| | S + E (only) | Edge 0.8 | | | Edge 0.9 | | | Edge 0.95 | | |
|---|---|---|---|---|---|---|---|---|---|---|
| Keystone Percentile | | 75th | 80th | 90th | 75th | 80th | 90th | 75th | 80th | 90th |
| Spatial (S) | 0.027 | 0.029 | 0.028 | 0.027 | 0.027 | 0.027 | 0.027 | 0.028 | 0.028 | 0.027 |
| Environmental (E) | 0.219 | 0.217 | 0.219 | 0.217 | 0.187 | 0.212 | 0.217 | 0.190 | 0.190 | 0.217 |
| Biotic (B) | | 0.007 | 0.003 | 0.002 | 0.007 | 0.001 | 0.002 | 0.009 | 0.009 | 0.002 |
| E + S | 0.109 | 0.109 | 0.111 | 0.111 | 0.073 | 0.104 | 0.111 | 0.077 | 0.077 | 0.111 |
| E + B | | 0.003 | 0.001 | 0.003 | 0.033 | 0.007 | 0.003 | 0.029 | 0.029 | 0.003 |
| S + B | | -0.002 | -0.001 | -0.001 | 0.000 | 0.000 | -0.001 | -0.001 | -0.001 | -0.001 |
| E + S + B | | 0.000 | -0.002 | -0.002 | 0.036 | 0.005 | -0.002 | 0.032 | 0.032 | -0.002 |
| Residuals | 0.640 | 0.638 | 0.642 | 0.643 | 0.638 | 0.643 | 0.643 | 0.636 | 0.636 | 0.643 |

**C. P-values from RDA for fungi as target community**

| | S + E (only) | Edge 0.8 | | | Edge 0.9 | | | Edge 0.95 | | |
|---|---|---|---|---|---|---|---|---|---|---|
| Keystone Percentile | | 75th | 80th | 90th | 75th | 80th | 90th | 75th | 80th | 90th |
| Spatial | 0.001 | 0.024 | 0.014 | 0.001 | 0.034 | 0.026 | 0.001 | 0.168 | 0.006 | 0.001 |
| Environmental | 0.001 | 0.011 | 0.002 | 0.001 | 0.024 | 0.001 | 0.001 | 0.027 | 0.001 | 0.001 |
| Biotic | | 0.014 | 0.008 | 0.004 | 0.026 | 0.098 | 0.005 | 0.242 | 0.105 | 0.006 |
| global model | 0.001 | 0.001 | 0.001 | 0.001 | 0.001 | 0.001 | 0.001 | 0.001 | 0.001 | 0.001 |

**D. P-vales from RDA for bacteria as target community**

| | S + E (only) | Edge 0.8 | | | Edge 0.9 | | | Edge 0.95 | | |
|---|---|---|---|---|---|---|---|---|---|---|
| Keystone Percentile | | 75th | 80th | 90th | 75th | 80th | 90th | 75th | 80th | 90th |
| Spatial | 0.001 | 0.001 | 0.001 | 0.001 | 0.001 | 0.001 | 0.001 | 0.001 | 0.001 | 0.001 |
| Environmental | 0.001 | 0.001 | 0.001 | 0.001 | 0.001 | 0.001 | 0.001 | 0.001 | 0.001 | 0.001 |
| Biotic | | 0.028 | 0.136 | 0.118 | 0.077 | 0.277 | 0.118 | 0.035 | 0.035 | 0.118 |
| global model | 0.001 | 0.001 | 0.001 | 0.001 | 0.001 | 0.001 | 0.001 | 0.001 | 0.001 | 0.001 |

**Table S15.** Results of variance partitioning analyses for root communities, repeated for the different network parameters tested to compare sensitivity of outcomes. Dependent variable is the Hellinger-transformed (A) fungal or (B) bacterial root community. Explanatory tables include standardized environmental and spatial variables (Table S7), and biotic interactions represented as the alpha diversity of the other kingdom (e.g. for fungi as the dependent matrix, alpha diversity of bacteria is the independent matrix) at different thresholds for each network edge-wise variability estimate (0.8, 0.9, 0.95). P-values for each variance partitioning, or redundancy analysis (RDA), model for leaf (C) fungal or (D) bacterial community.

| | **A. Fungi as target community** | | | | **B. Bacteria as target community** | | | |
|---|---|---|---|---|---|---|---|---|
| | S + E (only) | Edge 0.8 | Edge 0.9 | Edge 0.95 | S + E (only) | Edge 0.8 | Edge 0.9 | Edge 0.95 |
| Spatial (S) | 0.033 | 0.033 | 0.033 | 0.033 | 0.027 | 0.028 | 0.028 | 0.028 |
| Environmental (E) | 0.186 | 0.162 | 0.163 | 0.165 | 0.219 | 0.186 | 0.188 | 0.191 |
| Biotic (B) | | 0.003 | 0.003 | 0.003 | | 0.006 | 0.006 | 0.005 |
| E + S | 0.043 | 0.050 | 0.050 | 0.050 | 0.109 | 0.105 | 0.107 | 0.109 |
| E + B | | 0.024 | 0.023 | 0.021 | | 0.033 | 0.032 | 0.029 |
| S + B | | 0.000 | -0.001 | -0.001 | | -0.001 | -0.001 | -0.001 |
| E + S + B | | -0.007 | -0.007 | -0.007 | | 0.004 | 0.002 | 0.000 |
| Residuals | 0.738 | 0.735 | 0.735 | 0.735 | 0.640 | 0.639 | 0.639 | 0.640 |

| | **C. P-values from RDA for fungi as target community** | | | | **D. P-values from RDA for bacteria as target community** | | | |
|---|---|---|---|---|---|---|---|---|
| | S + E (only) | Edge 0.8 | Edge 0.9 | Edge 0.95 | S + E (only) | Edge 0.8 | Edge 0.9 | Edge 0.95 |
| Spatial | 0.001 | 0.001 | 0.001 | 0.001 | 0.001 | 0.001 | 0.001 | 0.001 |
| Environmental | 0.001 | 0.001 | 0.001 | 0.001 | 0.001 | 0.001 | 0.001 | 0.001 |
| Biotic | | 0.092 | 0.099 | 0.106 | | 0.003 | 0.002 | 0.005 |
| global model | 0.001 | 0.001 | 0.001 | 0.001 | 0.001 | 0.001 | 0.001 | 0.001 |

**Table S16.** Frequency tables of (A) direct and (B) diffuse interacting taxa, identified from the networks resulting from a 0.8 edge-wise variability estimate and keystones as nodes in the 75th percentile for both degree and eigenvector centrality.

**A. Frequency of the number of direct cross-kingdom interactions within each community. From the network with 0.8 edge-wise variability estimate.**

| | Leaf | | Root | |
|---|---|---|---|---|
| Direct interactions | Fungi | Bacteria | Fungi | Bacteria |
| 0 | 72 | 175 | 39 | 340 |
| 1 | 22 | 32 | 27 | 34 |
| 2 | 9 | 4 | 4 | 2 |
| 3 | 1 | 1 | 1 | 0 |
| Total | 104 | 212 | 71 | 376 |

**B. Frequency of genera of the diffuse cross-kingdom interactions within each community. From the network with 0.8 edge-wise variability estimate.**

| Leaf | | | | Root | | | |
|---|---|---|---|---|---|---|---|
| Bacteria Keystones | | Fungi Keystones | | Bacteria Keystones | | Fungi Keystones | |
| Genus | Frequency | Genus | Frequency | Genus | Frequency | Genus | Frequency |
| Acidibacter | 1 | Aphanophora | 1 | Acidibacter | 2 | Shivasia | 1 |
| Actinomycetospora | 1 | Cumuliphoma | 1 | Acidicapsa | 1 | Paraglomus | 2 |
| Aggregicoccus | 2 | Curvularia | 1 | Acidobacterium | 2 | | |
| Aureimonas | 1 | Knufia | 1 | Actinopolyspora | 1 | | |
| Beijerinckia | 1 | Montagnula | 1 | Aetherobacter | 1 | | |
| Beutenbergia | 1 | Papiliotrema | 1 | Ammoniphilus | 1 | | |
| Blastomonas | 1 | Paradevriesia | 1 | Asticcacaulis | 1 | | |
| Chelatococcus | 1 | Phaeosphaeria | 1 | Bactoderma | 1 | | |
| Fimbriimonas | 2 | Phyllozyma | 1 | Blastomonas | 1 | | |
| Fulvimarina | 1 | | | Bryobacter | 1 | | |
| Martelella | 1 | | | Burkholderia | 1 | | |
| Massilia | 1 | | | Catelliglobosispora | 1 | | |
| Methylobacterium | 1 | | | Catenulispora | 1 | | |
| Nocardioides | 3 | | | Chthoniobacter | 2 | | |
| Ottowia | 1 | | | Cystobacter | 1 | | |
| Pseudoxanthomonas | 1 | | | Deferrisoma | 1 | | |
| Ramlibacter | 1 | | | Dongia | 1 | | |
| Rickettsia | 1 | | | Enhydrobacter | 1 | | |
| Sphingomonas | 2 | | | Filomicrobium | 4 | | |
| Spirochaeta | 1 | | | Gaiella | 3 | | |
| | | | | Gemmatimonas | 1 | | |
| | | | | Haliangium | 3 | | |
| | | | | Immundisolibacter | 1 | | |
| | | | | Inquilinus | 2 | | |
| | | | | Kordiimonas | 1 | | |
| | | | | Koribacter | 4 | | |
| | | | | Ktedonosporobacter | 2 | | |
| | | | | Labilithrix | 1 | | |
| | | | | Luteitalea | 1 | | |
| | | | | Micropepsis | 2 | | |
| | | | | Mycobacterium | 4 | | |
| | | | | Niastella | 1 | | |
| | | | | Opitutus | 1 | | |
| | | | | Phototrophicus | 1 | | |
| | | | | Povalibacter | 2 | | |
| | | | | Sphingopyxis | 1 | | |
| | | | | Streptacidiphilus | 1 | | |
| | | | | Thermoanaerobaculum | 1 | | |
| | | | | Thermogemmatispora | 1 | | |
| | | | | Thermogutta | 1 | | |
| | | | | Thermoleophilum | 2 | | |
| | | | | Thiohalospira | 1 | | |
| | | | | Trebonia | 3 | | |
| | | | | unclassified | 1 | | |
| | | | | Verminephrobacter | 1 | | |
| | | | | Verrucomicrobia | 1 | | |
| TOTAL | 25 | | 9 | | 69 | | 3 |

**Table S17.** Results of testing for phylogenetic signal of interaction type (i.e., direct, diffuse, within-kingdom) using Pagel's λ for each of the (A) leaf and (B) root microbial communities.

**A. Leaf communites**

| | null AIC | null log-likelihood | observed AIC | observed log-likelihood | Chi-squared p-value |
|---|---|---|---|---|---|
| Bacteria | 18.307 | -6.154 | 18.307 | -6.154 | 0.9999991 |
| Fungi | 132.505 | -64.193 | 132.505 | -64.193 | 0.9999996 |

**B. Root communites**

| | null AIC | null log-likelihood | observed AIC | observed log-likelihood | Chi-squared p-value |
|---|---|---|---|---|---|
| Bacteria | 617.303 | -306.636 | 617.303 | -306.636 | 0.9999947 |
| Fungi | 102.216 | -49.0198 | 102.216 | -49.020 | 0.9999997 |

**Table S18.** Results of permANOVA of interacting (A-D) leaf and (E-H) root taxa on environmental variables. Environmental variables that were both significant and had an $R^2 > 0.02$ were used in the environmental gradient analysis. If no environmental variables had $R^2 > 0.02$ then the top two significant variables were selected. Variables selected for further analysis are in bold text.

**Leaf**

**A. Directly interacting bacteria as dependent variable**

| | df | SS | MS | $R^2$ | *F* | *Z* | *P* |
|---|---|---|---|---|---|---|---|
| MAT | 1 | 43.2 | 43.2 | 0.012 | 1.386 | 1.115 | 0.137 |
| MAP | 1 | 54.7 | 54.7 | 0.015 | 1.753 | 1.927 | 0.033 |
| moisture | 1 | 41.6 | 41.6 | 0.011 | 1.334 | 1.015 | 0.138 |
| **pH** | **1** | **106.1** | **106.1** | **0.029** | **3.405** | **3.662** | **0.001** |
| P | 1 | 60.1 | 60.1 | 0.016 | 1.927 | 1.949 | 0.029 |
| K | 1 | 72.4 | 72.4 | 0.020 | 2.322 | 2.644 | 0.005 |
| **sand** | **1** | **78.5** | **78.5** | **0.021** | **2.519** | **2.735** | **0.004** |
| clay | 1 | 51.8 | 51.8 | 0.014 | 1.663 | 1.621 | 0.053 |
| carbon | 1 | 43.2 | 43.2 | 0.012 | 1.387 | 1.168 | 0.120 |
| nitrogen | 1 | 46.3 | 46.3 | 0.013 | 1.485 | 1.311 | 0.095 |
| MBC | 1 | 33.7 | 33.7 | 0.009 | 1.081 | 0.366 | 0.347 |
| **plant basal area** | **1** | **84.3** | **84.3** | **0.023** | **2.704** | **2.987** | **0.003** |
| plant height | 1 | 51.1 | 51.1 | 0.014 | 1.638 | 1.545 | 0.069 |
| Residuals | 86 | 2680.6 | 31.2 | 0.732 | | | |
| Total | 99 | 3663 | | | | | |

**Root**

**E. Directly interacting bacteria as dependent variable**

| | df | SS | MS | $R^2$ | *F* | *Z* | *P* |
|---|---|---|---|---|---|---|---|
| MAT | 1 | 56.5 | 56.5 | 0.014 | 1.873 | 2.113 | 0.016 |
| MAP | 1 | 77.4 | 77.4 | 0.020 | 2.568 | 2.586 | 0.004 |
| moisture | 1 | 57.4 | 57.4 | 0.015 | 1.903 | 2.006 | 0.028 |
| pH | **1** | **111.2** | **111.2** | **0.028** | **3.688** | **3.997** | **0.001** |
| P | 1 | 48.9 | 48.9 | 0.012 | 1.623 | 1.557 | 0.057 |
| K | 1 | 40.7 | 40.7 | 0.010 | 1.350 | 1.064 | 0.141 |
| sand | 1 | 36.9 | 36.9 | 0.009 | 1.226 | 0.737 | 0.229 |
| clay | 1 | 34.5 | 34.5 | 0.009 | 1.144 | 0.565 | 0.278 |
| carbon | **1** | **82.9** | **82.9** | **0.021** | **2.752** | **3.021** | **0.002** |
| nitrogen | 1 | 34.6 | 34.6 | 0.009 | 1.148 | 0.519 | 0.294 |
| MBC | 1 | 39.5 | 39.5 | 0.010 | 1.311 | 0.930 | 0.182 |
| plant basal area | 1 | 33.4 | 33.4 | 0.009 | 1.108 | 0.466 | 0.316 |
| plant height | 1 | 28.2 | 28.2 | 0.007 | 0.936 | -0.051 | 0.506 |
| Residuals | 96 | 2893.4 | 30.1 | 0.737 | | | |
| Total | 109 | 3924 | | | | | |

**Leaf**

**B. Directly interacting fungi as dependent variable**

| | df | SS | MS | $R^2$ | *F* | *Z* | *P* |
|---|---|---|---|---|---|---|---|
| **MAT** | **1** | **101.4** | **101.4** | **0.033** | **3.945** | **4.746** | **0.001** |
| MAP | 1 | 60.7 | 60.7 | 0.020 | 2.364 | 2.676 | 0.003 |
| moisture | 1 | 38.1 | 38.1 | 0.012 | 1.484 | 1.379 | 0.084 |
| pH | 1 | 47.0 | 47.0 | 0.015 | 1.828 | 2.062 | 0.026 |
| P | 1 | 38.1 | 38.1 | 0.012 | 1.482 | 1.243 | 0.113 |
| K | 1 | 53.4 | 53.4 | 0.017 | 2.078 | 2.260 | 0.016 |
| **sand** | **1** | **68.8** | **68.8** | **0.022** | **2.679** | **3.032** | **0.001** |
| clay | 1 | 51.0 | 51.0 | 0.017 | 1.985 | 2.154 | 0.014 |
| carbon | 1 | 41.3 | 41.3 | 0.013 | 1.607 | 1.612 | 0.055 |
| nitrogen | 1 | 41.2 | 41.2 | 0.013 | 1.605 | 1.559 | 0.058 |
| MBC | 1 | 24.6 | 24.6 | 0.008 | 0.958 | 0.013 | 0.497 |
| plant basal area | 1 | 37.0 | 37.0 | 0.012 | 1.441 | 1.307 | 0.111 |
| plant height | 1 | 46.9 | 46.9 | 0.015 | 1.824 | 1.997 | 0.025 |
| Residuals | 86 | 2209.7 | 25.7 | 0.720 | | | |
| Total | 99 | 3069.0 | | | | | |

**Root**

**F. Directly interacting fungi as dependent variable**

| | df | SS | MS | $R^2$ | *F* | *Z* | *P* |
|---|---|---|---|---|---|---|---|
| MAT | 1 | 43.3 | 43.3 | 0.013 | 1.471 | 1.477 | 0.067 |
| **MAP** | **1** | **45.6** | **45.6** | **0.013** | **1.550** | **1.773** | **0.037** |
| moisture | 1 | 45.0 | 45.0 | 0.013 | 1.531 | 1.613 | 0.057 |
| pH | 1 | 36.0 | 36.0 | 0.011 | 1.224 | 0.809 | 0.206 |
| P | 1 | 42.7 | 42.7 | 0.012 | 1.453 | 1.342 | 0.094 |
| K | 1 | 37.6 | 37.6 | 0.011 | 1.279 | 0.981 | 0.160 |
| sand | 1 | 34.5 | 34.5 | 0.010 | 1.173 | 0.653 | 0.260 |
| clay | 1 | 32.6 | 32.6 | 0.010 | 1.109 | 0.514 | 0.307 |
| carbon | 1 | 33.3 | 33.3 | 0.010 | 1.131 | 0.531 | 0.311 |
| **nitrogen** | **1** | **46.0** | **46.0** | **0.013** | **1.566** | **1.674** | **0.046** |
| MBC | 1 | 24.4 | 24.4 | 0.007 | 0.831 | -0.473 | 0.674 |
| plant basal area | 1 | 26.6 | 26.6 | 0.008 | 0.905 | -0.237 | 0.607 |
| plant height | 1 | 38.0 | 38.0 | 0.011 | 1.293 | 1.010 | 0.151 |
| Residuals | 94 | 2764.2 | 29.4 | 0.807 | | | |
| Total | 107 | 3424.0 | | | | | |

**Table S18** (continued)

**Leaf**

**C. Diffusely interacting bacteria as dependent variable**

| | df | SS | MS | $R^2$ | *F* | *Z* | *P* |
|---|---|---|---|---|---|---|---|
| MAT | 1 | 32.9 | 32.9 | 0.013 | 1.537 | 1.166 | 0.128 |
| MAP | 1 | 34.0 | 34.0 | 0.014 | 1.588 | 1.285 | 0.091 |
| moisture | 1 | 37.9 | 37.9 | 0.015 | 1.769 | 1.510 | 0.059 |
| **pH** | **1** | **74.1** | **74.1** | **0.030** | **3.455** | **3.344** | **0.001** |
| P | 1 | 21.0 | 21.0 | 0.009 | 0.981 | 0.191 | 0.406 |
| K | 1 | 33.0 | 33.0 | 0.013 | 1.540 | 1.159 | 0.122 |
| **sand** | **1** | **101.3** | **101.3** | **0.041** | **4.728** | **3.739** | **0.001** |
| clay | 1 | 34.4 | 34.4 | 0.014 | 1.605 | 1.287 | 0.100 |
| carbon | 1 | 28.2 | 28.2 | 0.011 | 1.314 | 0.850 | 0.192 |
| nitrogen | 1 | 23.0 | 23.0 | 0.009 | 1.074 | 0.330 | 0.370 |
| MBC | 1 | 32.0 | 32.0 | 0.013 | 1.492 | 1.163 | 0.119 |
| **plant basal area** | **1** | **63.8** | **63.8** | **0.026** | **2.977** | **2.532** | **0.006** |
| plant height | 1 | 21.2 | 21.2 | 0.009 | 0.990 | 0.187 | 0.429 |
| Residuals | 86 | 1843.2 | 21.4 | 0.745 | | | |
| Total | 99 | 2475.0 | | | | | |

**Root**

**G. Diffusely interacting bacteria as dependent variable**

| | df | SS | MS | $R^2$ | *F* | *Z* | *P* |
|---|---|---|---|---|---|---|---|
| MAT | 1 | 87.8 | 87.8 | 0.012 | 1.586 | 1.299 | 0.070 |
| MAP | 1 | 132.8 | 132.8 | 0.018 | 2.397 | 2.280 | 0.020 |
| moisture | 1 | 86.6 | 86.6 | 0.012 | 1.563 | 1.410 | 0.100 |
| **pH** | **1** | **183.6** | **183.6** | **0.025** | **3.315** | **4.186** | **0.010** |
| P | 1 | 80.6 | 80.6 | 0.011 | 1.456 | 1.199 | 0.110 |
| K | 1 | 59.1 | 59.1 | 0.008 | 1.068 | 0.311 | 0.350 |
| sand | 1 | 87.3 | 87.3 | 0.012 | 1.576 | 1.690 | 0.060 |
| clay | 1 | 80.3 | 80.3 | 0.011 | 1.450 | 1.235 | 0.120 |
| carbon | 1 | 96.8 | 96.8 | 0.013 | 1.747 | 1.798 | 0.050 |
| nitrogen | 1 | 64.1 | 64.1 | 0.009 | 1.157 | 0.651 | 0.290 |
| MBC | 1 | 57.5 | 57.5 | 0.008 | 1.038 | 0.246 | 0.370 |
| plant basal area | 1 | 57.0 | 57.0 | 0.008 | 1.029 | 0.088 | 0.490 |
| plant height | 1 | 111.3 | 111.3 | 0.015 | 2.009 | 1.874 | 0.040 |
| Residuals | 94 | 5206.6 | 55.4 | 0.705 | | | |
| Total | 107 | 7383.0 | | | | | |

**Leaf**

**D. Diffusely interacting fungi as dependent variable**

| | df | SS | MS | $R^2$ | *F* | *Z* | *P* |
|---|---|---|---|---|---|---|---|
| MAT | 1 | 12.7 | 12.7 | 0.014 | 1.582 | 1.111 | 0.120 |
| MAP | 1 | 6.6 | 6.6 | 0.007 | 0.820 | -0.137 | 0.571 |
| moisture | 1 | 6.8 | 6.8 | 0.008 | 0.846 | -0.156 | 0.560 |
| pH | 1 | 14.7 | 14.7 | 0.016 | 1.833 | 1.444 | 0.076 |
| P | 1 | 11.3 | 11.3 | 0.013 | 1.409 | 0.829 | 0.182 |
| K | 1 | 13.9 | 13.9 | 0.016 | 1.737 | 1.251 | 0.097 |
| **sand** | **1** | **22.4** | **22.4** | **0.025** | **2.798** | **2.380** | **0.008** |
| clay | 1 | 11.1 | 11.1 | 0.013 | 1.392 | 0.838 | 0.194 |
| carbon | 1 | 6.9 | 6.9 | 0.008 | 0.857 | -0.141 | 0.558 |
| nitrogen | 1 | 8.4 | 8.4 | 0.009 | 1.044 | 0.260 | 0.402 |
| MBC | 1 | 8.7 | 8.7 | 0.010 | 1.093 | 0.387 | 0.342 |
| plant basal area | 1 | 9.3 | 9.3 | 0.010 | 1.158 | 0.436 | 0.325 |
| plant height | 1 | 12.4 | 12.4 | 0.014 | 1.552 | 1.069 | 0.135 |
| Residuals | 86 | 688.1 | 8.0 | 0.772 | | | |
| Total | 99 | 891.0 | | | | | |

**Root**

**H. Diffusely interacting fungi as dependent variable**

| | df | SS | MS | $R^2$ | *F* | *Z* | *P* |
|---|---|---|---|---|---|---|---|
| MAT | 1 | 3.5 | 3.5 | 0.011 | 1.154 | 0.402 | 0.350 |
| MAP | 1 | 0.7 | 0.7 | 0.002 | 0.219 | -1.240 | 0.891 |
| moisture | 1 | 0.9 | 0.9 | 0.003 | 0.304 | -0.871 | 0.797 |
| pH | 1 | 1.0 | 0.9 | 0.003 | 0.315 | -0.875 | 0.807 |
| P | 1 | 3.4 | 3.4 | 0.010 | 1.122 | 0.435 | 0.328 |
| K | 1 | 1.2 | 1.2 | 0.004 | 0.405 | -0.644 | 0.726 |
| sand | 1 | 0.6 | 0.6 | 0.002 | 0.190 | -1.402 | 0.923 |
| clay | 1 | 4.5 | 4.4 | 0.014 | 1.474 | 0.755 | 0.222 |
| carbon | 1 | 1.0 | 1.0 | 0.003 | 0.336 | -0.946 | 0.820 |
| nitrogen | 1 | 1.7 | 1.7 | 0.005 | 0.565 | -0.342 | 0.640 |
| MBC | 1 | 2.7 | 2.7 | 0.008 | 0.890 | 0.121 | 0.458 |
| plant basal area | 1 | 1.7 | 1.6 | 0.005 | 0.546 | -0.319 | 0.632 |
| **plant height** | **1** | **8.7** | **8.7** | **0.027** | **2.880** | **1.755** | **0.037** |
| Residuals | 96 | 289.7 | 3.0 | 0.886 | | | |
| Total | 109 | 327.0 | | | | | |

**Table S19.** Results from environmental gradient analysis (TITAN2) showing ASVs that significantly track the identified variables from PERMANOVA (Table S18). Tables describe the following statistics for ASVs in (A-F) bacterial and (G-I) fungal leaf communities: ienv.cp: environmental change point for each taxon based on IndVal maximum, zenv.cp: environmental change point for each taxon based on z maximum (default), freq: number of non-zero abundance values per taxon, maxgrp: 1 if z-(negative response) or 2 if z+ (positive response) indicated by filter column which indicates whether each taxa met purity and reliability criteria and direction, IndVal: indicator value statistic scaled 0-100%, obsiv.prob: probability of an equal or larger IndVal from random permutation, z score: IndVal z score, 5%-95%: change point quantiles among bootstrap replicates, purity: proportion of replicates matching observed maxgrp assignment, reliability: proportion of replicate obsiv.prob values <= 0.05, z.median: median score magnitude across all bootstrap replicates.

**A. Bacteria direct interactors associated with pH**

| ASV | ienv.cp | zenv.cp | freq | maxgrp | IndVal | obsiv.prob | zscore | 5% | 10% | 50% | 90% | 95% | purity | reliability | z.median |
|---|---|---|---|---|---|---|---|---|---|---|---|---|---|---|---|
| ASV_842 | 4.328 | 5.258 | 20 | 1 | 44.44 | 0.004 | 6.39 | 4.293 | 4.310 | 5.213 | 5.288 | 5.323 | 1.000 | 1.000 | 7.472 |
| ASV_993 | 5.285 | 5.285 | 30 | 1 | 53.97 | 0.004 | 7.72 | 5.203 | 5.213 | 5.288 | 5.508 | 6.245 | 1.000 | 1.000 | 7.783 |
| ASV_4017 | 4.328 | 4.513 | 23 | 1 | 89.95 | 0.004 | 11.00 | 4.364 | 4.425 | 5.200 | 5.323 | 5.365 | 1.000 | 1.000 | 10.481 |
| ASV_443 | 4.383 | 4.328 | 13 | 1 | 90.07 | 0.004 | 11.17 | 4.320 | 4.328 | 4.439 | 5.058 | 5.110 | 1.000 | 0.994 | 11.679 |
| ASV_1729 | 4.383 | 5.693 | 37 | 1 | 34.88 | 0.004 | 3.86 | 4.383 | 4.660 | 5.745 | 6.501 | 6.723 | 0.990 | 0.990 | 4.843 |
| ASV_3885 | 4.328 | 5.038 | 13 | 1 | 39.78 | 0.004 | 6.06 | 4.283 | 4.293 | 5.063 | 5.390 | 5.855 | 0.980 | 0.988 | 7.944 |
| ASV_13 | 4.513 | 5.870 | 93 | 2 | 71.15 | 0.004 | 4.66 | 4.465 | 4.513 | 5.730 | 6.200 | 6.230 | 1.000 | 1.000 | 5.517 |
| ASV_45 | 4.383 | 6.350 | 94 | 2 | 74.67 | 0.004 | 4.18 | 5.427 | 5.489 | 5.834 | 6.350 | 6.353 | 1.000 | 1.000 | 4.576 |
| ASV_189 | 4.513 | 5.575 | 86 | 2 | 67.36 | 0.004 | 4.99 | 4.530 | 4.615 | 5.570 | 6.196 | 6.350 | 1.000 | 1.000 | 5.809 |
| ASV_1151 | 5.395 | 5.808 | 25 | 2 | 34.60 | 0.004 | 4.68 | 5.380 | 5.390 | 5.575 | 5.856 | 5.889 | 1.000 | 1.000 | 5.237 |
| ASV_1246 | 6.723 | 5.745 | 38 | 2 | 47.20 | 0.004 | 4.89 | 5.523 | 5.550 | 5.808 | 6.738 | 6.813 | 1.000 | 0.998 | 6.168 |
| ASV_3132 | 5.638 | 5.693 | 20 | 2 | 32.25 | 0.004 | 5.53 | 5.592 | 5.620 | 5.680 | 5.835 | 5.889 | 1.000 | 0.998 | 6.152 |
| ASV_121 | 6.350 | 5.923 | 45 | 2 | 53.71 | 0.004 | 3.91 | 5.395 | 5.630 | 5.985 | 6.350 | 6.353 | 1.000 | 0.996 | 4.540 |
| ASV_166 | 6.640 | 5.540 | 44 | 2 | 49.39 | 0.008 | 4.07 | 5.395 | 5.395 | 5.720 | 6.580 | 6.640 | 1.000 | 0.996 | 5.164 |
| ASV_9 | 4.513 | 5.480 | 100 | 2 | 65.11 | 0.004 | 3.73 | 4.465 | 4.513 | 5.460 | 5.771 | 6.069 | 0.996 | 0.998 | 4.348 |
| ASV_116 | 4.513 | 5.563 | 67 | 2 | 64.45 | 0.004 | 3.85 | 5.355 | 5.460 | 5.638 | 6.305 | 6.613 | 0.996 | 0.994 | 4.782 |
| ASV_11551 | 6.795 | 5.808 | 15 | 2 | 23.54 | 0.004 | 4.44 | 5.395 | 5.570 | 5.820 | 6.795 | 6.961 | 0.996 | 0.986 | 5.262 |
| ASV_3230 | 6.075 | 5.888 | 22 | 2 | 30.44 | 0.004 | 4.39 | 5.670 | 5.777 | 5.950 | 6.723 | 6.833 | 0.996 | 0.980 | 5.293 |
| ASV_12 | 4.513 | 5.708 | 76 | 2 | 68.77 | 0.004 | 5.64 | 5.038 | 5.185 | 5.710 | 5.795 | 5.823 | 0.992 | 1.000 | 5.694 |
| ASV_6 | 4.513 | 4.513 | 100 | 2 | 75.13 | 0.008 | 4.29 | 4.458 | 4.465 | 5.154 | 5.693 | 6.820 | 0.984 | 0.974 | 4.303 |
| ASV_320 | 4.513 | 5.575 | 68 | 2 | 52.93 | 0.016 | 3.49 | 4.465 | 4.513 | 5.568 | 5.866 | 6.621 | 0.974 | 0.976 | 4.210 |

**Table S19** (continued)

**B. Bacteria direct interactors associated with plant basal area**

| ASV | ienv.cp | zenv.cp | freq | maxgrp | IndVal | obsiv.prob | zscore | 5% | 10% | 50% | 90% | 95% | purity | reliability | z.median |
|---|---|---|---|---|---|---|---|---|---|---|---|---|---|---|---|
| ASV_166 | 0.037 | 0.037 | 44 | 1 | 61.55 | 0.008 | 3.92 | 0.020 | 0.020 | 0.041 | 0.225 | 0.267 | 0.996 | 0.990 | 4.811 |
| ASV_3885 | 0.074 | 0.074 | 13 | 1 | 23.86 | 0.004 | 4.44 | 0.065 | 0.067 | 0.074 | 0.113 | 0.116 | 0.972 | 0.954 | 4.905 |
| ASV_9 | 0.090 | 0.090 | 100 | 2 | 65.13 | 0.004 | 3.76 | 0.070 | 0.074 | 0.113 | 0.186 | 0.202 | 0.998 | 0.992 | 4.431 |
| ASV_6310 | 0.530 | 0.176 | 16 | 2 | 38.56 | 0.004 | 8.53 | 0.168 | 0.171 | 0.178 | 0.507 | 0.530 | 0.994 | 1.000 | 8.753 |
| ASV_474 | 0.024 | 0.170 | 59 | 2 | 52.27 | 0.008 | 3.74 | 0.031 | 0.067 | 0.170 | 0.460 | 0.473 | 0.994 | 0.990 | 4.904 |
| ASV_747 | 0.537 | 0.457 | 45 | 2 | 63.68 | 0.004 | 5.02 | 0.124 | 0.128 | 0.207 | 0.467 | 0.505 | 0.984 | 0.994 | 6.331 |
| ASV_2248 | 0.470 | 0.470 | 20 | 2 | 66.54 | 0.004 | 8.14 | 0.135 | 0.202 | 0.470 | 0.474 | 0.497 | 0.982 | 1.000 | 8.702 |
| ASV_1751 | 0.530 | 0.537 | 26 | 2 | 72.33 | 0.004 | 5.85 | 0.106 | 0.115 | 0.503 | 0.570 | 0.628 | 0.968 | 0.990 | 6.750 |
| ASV_3230 | 0.530 | 0.166 | 22 | 2 | 40.23 | 0.004 | 7.00 | 0.137 | 0.144 | 0.170 | 0.304 | 0.386 | 0.960 | 1.000 | 7.875 |

**C. Bacteria direct interactors associated with percent sand**

| ASV | ienv.cp | zenv.cp | freq | maxgrp | IndVal | obsiv.prob | zscore | 5% | 10% | 50% | 90% | 95% | purity | reliability | z.median |
|---|---|---|---|---|---|---|---|---|---|---|---|---|---|---|---|
| ASV_1454 | 36.004 | 38.911 | 10 | 1 | 72.48 | 0.004 | 14.10 | 33.396 | 35.623 | 41.086 | 52.053 | 52.966 | 1.000 | 1.000 | 14.846 |
| ASV_4693 | 33.017 | 33.017 | 10 | 1 | 60.51 | 0.004 | 9.74 | 32.508 | 33.017 | 48.494 | 52.867 | 53.496 | 0.996 | 1.000 | 10.011 |
| ASV_1306 | 36.004 | 39.934 | 21 | 1 | 83.52 | 0.004 | 9.86 | 33.017 | 35.623 | 39.934 | 48.292 | 51.072 | 0.994 | 1.000 | 10.804 |
| ASV_842 | 73.742 | 73.742 | 20 | 1 | 27.69 | 0.008 | 3.12 | 63.571 | 63.732 | 71.540 | 73.932 | 74.520 | 0.956 | 0.968 | 3.254 |
| ASV_9 | 44.666 | 56.975 | 100 | 2 | 73.61 | 0.004 | 6.50 | 46.007 | 54.250 | 62.578 | 64.639 | 64.902 | 1.000 | 1.000 | 6.535 |
| ASV_12 | 58.593 | 64.639 | 76 | 2 | 87.94 | 0.004 | 9.10 | 59.769 | 62.214 | 64.741 | 67.580 | 68.104 | 1.000 | 1.000 | 9.582 |
| ASV_3230 | 83.433 | 63.732 | 22 | 2 | 29.32 | 0.004 | 4.09 | 63.445 | 63.675 | 79.619 | 83.261 | 83.433 | 1.000 | 0.996 | 5.641 |
| ASV_6310 | 83.433 | 66.155 | 16 | 2 | 24.91 | 0.004 | 3.95 | 64.899 | 65.004 | 67.701 | 83.242 | 83.433 | 1.000 | 0.994 | 5.199 |
| ASV_626 | 79.895 | 79.895 | 37 | 2 | 46.15 | 0.004 | 3.84 | 53.719 | 54.250 | 74.603 | 80.261 | 81.908 | 0.994 | 0.992 | 4.928 |
| ASV_320 | 46.007 | 46.007 | 68 | 2 | 61.53 | 0.012 | 3.24 | 45.345 | 45.924 | 60.982 | 70.323 | 75.253 | 0.994 | 0.974 | 4.510 |
| ASV_4860 | 69.098 | 69.098 | 27 | 2 | 30.91 | 0.008 | 4.04 | 63.764 | 64.173 | 69.098 | 78.426 | 80.261 | 0.980 | 0.952 | 4.676 |
| ASV_474 | 47.500 | 59.807 | 59 | 2 | 53.47 | 0.008 | 3.55 | 46.445 | 47.702 | 59.807 | 75.253 | 80.276 | 0.952 | 0.980 | 4.284 |
| ASV_2996 | 83.433 | 83.433 | 21 | 2 | 62.36 | 0.004 | 5.70 | 58.447 | 62.214 | 83.242 | 84.526 | 84.526 | 0.950 | 0.972 | 5.983 |

**D. Bacteria diffuse interactors associated with pH**

| ASV | ienv.cp | zenv.cp | freq | maxgrp | IndVal | obsiv.prob | zscore | 5% | 10% | 50% | 90% | 95% | purity | reliability | z.median |
|---|---|---|---|---|---|---|---|---|---|---|---|---|---|---|---|
| ASV_136 | 4.513 | 4.513 | 81 | 2 | 82.72 | 0.004 | 4.44 | 4.465 | 4.513 | 5.305 | 6.010 | 6.230 | 0.998 | 0.996 | 5.101 |
| ASV_571 | 5.888 | 5.888 | 20 | 2 | 30.82 | 0.004 | 3.88 | 5.395 | 5.655 | 5.888 | 6.630 | 6.813 | 0.998 | 0.970 | 4.459 |
| ASV_743 | 6.280 | 6.245 | 18 | 2 | 37.01 | 0.004 | 6.97 | 5.808 | 5.959 | 6.128 | 6.345 | 6.350 | 0.988 | 1.000 | 6.832 |
| ASV_199 | 6.205 | 6.245 | 51 | 2 | 51.65 | 0.004 | 3.57 | 5.462 | 5.620 | 6.045 | 6.276 | 6.640 | 0.964 | 0.972 | 4.155 |
| ASV_4059 | 6.245 | 5.835 | 16 | 2 | 23.98 | 0.004 | 4.34 | 5.539 | 5.804 | 6.120 | 6.350 | 6.353 | 0.960 | 0.966 | 5.000 |

**Table S19** (continued)

**E. Bacteria diffuse interactors associated with plant basal area**

| ASV | ienv.cp | zenv.cp | freq | maxgrp | IndVal | obsiv.prob | zscore | 5% | 10% | 50% | 90% | 95% | purity | reliability | z.median |
|---|---|---|---|---|---|---|---|---|---|---|---|---|---|---|---|
| ASV_3422 | 0.457 | 0.423 | 15 | 2 | 50.03 | 0.004 | 8.90 | 0.290 | 0.304 | 0.434 | 0.470 | 0.540 | 0.994 | 0.984 | 8.580 |
| ASV_597 | 0.505 | 0.505 | 22 | 2 | 75.78 | 0.004 | 7.70 | 0.168 | 0.335 | 0.459 | 0.512 | 0.537 | 0.976 | 0.998 | 8.469 |
| ASV_689 | 0.470 | 0.470 | 24 | 2 | 57.96 | 0.004 | 6.10 | 0.207 | 0.267 | 0.460 | 0.505 | 0.505 | 0.968 | 0.976 | 6.521 |
| ASV_2009 | 0.505 | 0.470 | 20 | 2 | 50.24 | 0.004 | 5.87 | 0.107 | 0.208 | 0.470 | 0.512 | 0.567 | 0.956 | 0.972 | 6.549 |
| ASV_4016 | 0.537 | 0.537 | 16 | 2 | 45.65 | 0.028 | 3.26 | 0.088 | 0.110 | 0.225 | 0.635 | 0.665 | 0.954 | 0.956 | 5.101 |

**F. Bacteria diffuse interactors associated with percent sand**

| ASV | ienv.cp | zenv.cp | freq | maxgrp | IndVal | obsiv.prob | zscore | 5% | 10% | 50% | 90% | 95% | purity | reliability | z.median |
|---|---|---|---|---|---|---|---|---|---|---|---|---|---|---|---|
| ASV_571 | 52.053 | 52.053 | 20 | 1 | 36.76 | 0.004 | 4.64 | 49.398 | 51.886 | 53.080 | 70.323 | 71.495 | 1.000 | 0.966 | 4.724 |
| ASV_597 | 36.004 | 33.017 | 22 | 1 | 96.71 | 0.004 | 11.10 | 32.636 | 33.364 | 38.761 | 44.004 | 48.849 | 0.998 | 1.000 | 11.261 |
| ASV_2009 | 36.004 | 36.004 | 20 | 1 | 95.37 | 0.004 | 11.19 | 32.937 | 33.364 | 38.611 | 41.818 | 43.426 | 0.998 | 0.996 | 11.362 |
| ASV_470 | 32.572 | 33.017 | 25 | 1 | 97.68 | 0.004 | 10.69 | 32.636 | 33.017 | 38.180 | 41.897 | 43.772 | 0.996 | 1.000 | 10.421 |
| ASV_689 | 36.004 | 36.004 | 24 | 1 | 88.58 | 0.004 | 9.75 | 32.636 | 33.017 | 36.304 | 41.810 | 44.670 | 0.992 | 0.992 | 9.427 |
| ASV_5911 | 33.017 | 44.004 | 13 | 1 | 37.92 | 0.004 | 5.41 | 32.508 | 33.017 | 44.004 | 58.142 | 71.453 | 0.954 | 0.956 | 6.534 |
| ASV_4016 | 64.058 | 64.058 | 16 | 2 | 24.62 | 0.008 | 3.94 | 63.554 | 63.686 | 64.173 | 78.387 | 78.678 | 0.996 | 0.990 | 4.363 |
| ASV_9928 | 71.448 | 71.251 | 13 | 2 | 24.96 | 0.004 | 5.61 | 70.566 | 70.853 | 71.343 | 71.633 | 72.800 | 0.994 | 0.978 | 5.922 |

**G. Fungi direct interactors associated with MAT**

| ASV | ienv.cp | zenv.cp | freq | maxgrp | IndVal | obsiv.prob | zscore | 5% | 10% | 50% | 90% | 95% | purity | reliability | z.median |
|---|---|---|---|---|---|---|---|---|---|---|---|---|---|---|---|
| ASV_256 | 13.900 | 14.900 | 18 | 1 | 77.70 | 0.004 | 10.53 | 12.900 | 13.900 | 15.000 | 15.500 | 15.500 | 1.000 | 1.000 | 10.615 |
| ASV_458 | 13.900 | 13.900 | 23 | 1 | 96.19 | 0.004 | 10.70 | 12.900 | 12.900 | 14.900 | 15.500 | 15.500 | 1.000 | 1.000 | 10.189 |
| ASV_406 | 12.900 | 15.500 | 39 | 1 | 42.09 | 0.004 | 4.95 | 14.850 | 14.900 | 14.950 | 15.550 | 15.550 | 1.000 | 0.996 | 6.229 |
| ASV_53 | 15.000 | 15.000 | 91 | 1 | 71.06 | 0.004 | 5.92 | 15.000 | 15.000 | 15.500 | 15.500 | 15.553 | 0.994 | 0.996 | 5.614 |
| ASV_2 | 12.900 | 15.000 | 85 | 1 | 66.31 | 0.004 | 4.37 | 14.900 | 14.900 | 15.550 | 16.500 | 16.500 | 0.994 | 0.994 | 4.943 |
| ASV_7 | 15.700 | 15.800 | 45 | 1 | 47.69 | 0.004 | 5.04 | 15.600 | 15.700 | 15.750 | 15.800 | 15.800 | 0.982 | 0.996 | 5.004 |
| ASV_102 | 14.950 | 15.000 | 40 | 2 | 45.70 | 0.004 | 4.90 | 14.950 | 14.950 | 15.000 | 15.600 | 16.500 | 1.000 | 1.000 | 5.548 |
| ASV_115 | 15.800 | 15.800 | 32 | 2 | 65.68 | 0.004 | 11.01 | 15.600 | 15.600 | 15.800 | 15.800 | 16.500 | 1.000 | 1.000 | 11.592 |
| ASV_138 | 15.500 | 15.500 | 51 | 2 | 58.05 | 0.004 | 6.97 | 15.000 | 15.000 | 15.500 | 15.550 | 15.550 | 1.000 | 1.000 | 6.493 |
| ASV_130 | 16.150 | 15.600 | 42 | 2 | 55.35 | 0.004 | 7.07 | 15.500 | 15.500 | 15.600 | 16.500 | 16.500 | 1.000 | 1.000 | 7.395 |
| ASV_2659 | 16.500 | 16.500 | 17 | 2 | 66.68 | 0.004 | 6.40 | 15.488 | 15.500 | 15.800 | 16.500 | 16.500 | 1.000 | 1.000 | 7.706 |
| ASV_66 | 15.250 | 15.000 | 42 | 2 | 48.50 | 0.004 | 5.33 | 15.000 | 15.000 | 15.250 | 15.500 | 15.500 | 0.990 | 1.000 | 5.731 |

**Table S19** (continued)

**H. Fungi direct interactors associated with percent sand**

| ASV | ienv.cp | zenv.cp | freq | maxgrp | IndVal | obsiv.prob | zscore | 5% | 10% | 50% | 90% | 95% | purity | reliability | z.median |
|---|---|---|---|---|---|---|---|---|---|---|---|---|---|---|---|
| ASV_53 | 83.051 | 70.566 | 91 | 1 | 69.97 | 0.004 | 5.78 | 46.435 | 47.702 | 69.639 | 72.295 | 77.803 | 1.000 | 1.000 | 5.789 |
| ASV_256 | 36.004 | 38.911 | 18 | 1 | 53.30 | 0.004 | 6.97 | 35.623 | 36.004 | 52.152 | 60.616 | 63.686 | 1.000 | 0.994 | 7.931 |
| ASV_458 | 33.017 | 52.053 | 23 | 1 | 42.81 | 0.004 | 5.14 | 33.017 | 35.623 | 52.053 | 63.686 | 67.203 | 0.994 | 0.964 | 5.614 |
| ASV_7 | 83.433 | 62.214 | 45 | 2 | 60.10 | 0.004 | 7.69 | 58.168 | 59.769 | 63.445 | 64.232 | 66.907 | 1.000 | 1.000 | 7.660 |
| ASV_4 | 58.593 | 62.214 | 78 | 2 | 84.60 | 0.004 | 7.99 | 57.382 | 58.593 | 63.732 | 66.334 | 67.603 | 1.000 | 1.000 | 8.211 |
| ASV_138 | 81.570 | 71.251 | 51 | 2 | 73.06 | 0.004 | 10.68 | 67.777 | 68.143 | 71.308 | 76.041 | 77.362 | 1.000 | 1.000 | 11.160 |
| ASV_18 | 47.500 | 63.686 | 49 | 2 | 54.97 | 0.004 | 5.67 | 47.213 | 51.886 | 63.686 | 67.143 | 67.603 | 1.000 | 1.000 | 5.871 |
| ASV_130 | 71.540 | 71.540 | 42 | 2 | 56.14 | 0.004 | 7.00 | 64.790 | 64.902 | 71.466 | 75.446 | 75.849 | 1.000 | 1.000 | 7.148 |
| ASV_2659 | 70.566 | 70.905 | 17 | 2 | 29.33 | 0.004 | 5.48 | 64.953 | 69.788 | 70.566 | 71.540 | 76.334 | 1.000 | 0.998 | 5.844 |
| ASV_66 | 58.593 | 58.593 | 42 | 2 | 50.32 | 0.004 | 5.51 | 56.975 | 58.131 | 64.876 | 77.561 | 79.364 | 0.996 | 0.998 | 5.904 |

**I. Fungi diffuse interactors associated with percent sand**

| ASV | ienv.cp | zenv.cp | freq | maxgrp | IndVal | obsiv.prob | zscore | 5% | 10% | 50% | 90% | 95% | purity | reliability | z.median |
|---|---|---|---|---|---|---|---|---|---|---|---|---|---|---|---|
| ASV_135 | 64.640 | 71.250 | 28 | 1 | 42.34 | 0.004 | 6.06 | 50.894 | 60.385 | 65.535 | 71.250 | 71.445 | 1.000 | 1.000 | 6.928 |
| ASV_122 | 67.145 | 67.145 | 10 | 1 | 19.54 | 0.004 | 4.23 | 66.155 | 66.690 | 67.385 | 69.910 | 70.210 | 0.998 | 0.978 | 4.384 |
| ASV_278 | 36.005 | 36.005 | 16 | 1 | 84.02 | 0.004 | 12.32 | 33.020 | 33.400 | 36.305 | 44.665 | 45.920 | 0.996 | 0.994 | 10.521 |
| ASV_1519 | 79.630 | 79.630 | 19 | 2 | 54.91 | 0.004 | 9.05 | 66.155 | 76.240 | 79.630 | 80.390 | 81.910 | 1.000 | 1.000 | 9.387 |
| ASV_272 | 64.950 | 66.155 | 22 | 2 | 26.66 | 0.004 | 3.92 | 64.383 | 64.800 | 66.335 | 71.725 | 74.531 | 0.954 | 0.954 | 4.238 |

**Table S20.** Results from environmental gradient analysis (TITAN2) showing ASVs that significantly track the identified variables from PERMANOVA (Table S18). Tables describe the following statistics for ASVs in (A-F) bacterial and (G-I) fungal root communities: ienv.cp: environmental change point for each taxon based on IndVal maximum, zenv.cp: environmental change point for each taxon based on z maximum (default), freq: number of non-zero abundance values per taxon, maxgrp: 1 if z- (negative response) or 2 if z+ (positive response) indicated by filter column which indicates whether each taxa met purity and reliability criteria and direction, IndVal: indicator value statistic scaled 0-100%, obsiv.prob: probability of an equal or larger IndVal from random permutation, z score: IndVal z score, 5%-95%: change point quantiles among bootstrap replicates, purity: proportion of replicates matching observed maxgrp assignment, reliability: proportion of replicate obsiv.prob values <= 0.05, z.median: median score magnitude across all bootstrap replicates.

**A. Bacteria direct interactors associated with pH**

| ASV | ienv.cp | zenv.cp | freq | maxgrp | IndVal | obsiv.prob | zscore | 5% | 10% | 50% | 90% | 95% | purity | reliability | z.median |
|---|---|---|---|---|---|---|---|---|---|---|---|---|---|---|---|
| ASV_21 | 5.035 | 5.663 | 94 | 1 | 76.45 | 0.004 | 6.93 | 5.380 | 5.480 | 5.695 | 5.900 | 5.915 | 1.000 | 1.000 | 6.968 |
| ASV_84 | 5.680 | 5.663 | 53 | 1 | 81.47 | 0.004 | 13.75 | 5.424 | 5.523 | 5.680 | 5.725 | 5.760 | 1.000 | 1.000 | 13.875 |
| ASV_625 | 5.725 | 5.725 | 28 | 1 | 42.43 | 0.004 | 8.56 | 5.570 | 5.625 | 5.730 | 5.783 | 5.811 | 1.000 | 1.000 | 8.549 |
| ASV_1334 | 5.368 | 5.663 | 29 | 1 | 34.01 | 0.004 | 5.25 | 5.315 | 5.355 | 5.655 | 6.350 | 6.398 | 1.000 | 1.000 | 5.584 |
| ASV_322 | 6.075 | 5.950 | 33 | 1 | 37.05 | 0.004 | 5.54 | 5.725 | 5.869 | 6.025 | 6.120 | 6.145 | 0.992 | 1.000 | 5.805 |
| ASV_24 | 4.515 | 4.980 | 88 | 2 | 83.69 | 0.004 | 7.25 | 4.465 | 4.515 | 4.980 | 5.248 | 5.255 | 1.000 | 1.000 | 7.672 |
| ASV_38 | 6.840 | 6.103 | 83 | 2 | 84.13 | 0.004 | 6.07 | 5.187 | 5.276 | 5.783 | 6.280 | 6.400 | 1.000 | 1.000 | 6.742 |
| ASV_41 | 5.443 | 5.395 | 75 | 2 | 85.92 | 0.004 | 11.10 | 5.390 | 5.415 | 5.460 | 5.620 | 5.705 | 1.000 | 1.000 | 11.699 |
| ASV_61 | 5.390 | 5.395 | 64 | 2 | 77.43 | 0.004 | 9.29 | 5.370 | 5.380 | 5.428 | 5.548 | 5.680 | 1.000 | 1.000 | 9.620 |
| ASV_69 | 5.315 | 5.505 | 49 | 2 | 53.53 | 0.004 | 5.58 | 5.285 | 5.295 | 5.505 | 5.973 | 6.120 | 1.000 | 1.000 | 6.274 |
| ASV_89 | 5.625 | 5.625 | 71 | 2 | 85.03 | 0.004 | 10.61 | 5.443 | 5.505 | 5.650 | 5.715 | 5.725 | 1.000 | 1.000 | 11.512 |
| ASV_143 | 5.315 | 5.315 | 47 | 2 | 54.20 | 0.004 | 5.53 | 5.258 | 5.315 | 5.693 | 6.245 | 6.280 | 1.000 | 1.000 | 6.070 |
| ASV_165 | 5.443 | 5.443 | 52 | 2 | 59.16 | 0.004 | 6.19 | 5.295 | 5.323 | 5.443 | 5.835 | 5.983 | 1.000 | 1.000 | 6.512 |
| ASV_264 | 6.910 | 5.835 | 55 | 2 | 69.12 | 0.004 | 11.03 | 5.440 | 5.497 | 5.878 | 6.250 | 6.429 | 1.000 | 1.000 | 11.974 |
| ASV_595 | 6.008 | 5.993 | 33 | 2 | 38.46 | 0.004 | 5.90 | 5.248 | 5.322 | 5.783 | 6.023 | 6.073 | 1.000 | 1.000 | 6.639 |
| ASV_28 | 4.433 | 5.200 | 88 | 2 | 77.42 | 0.004 | 7.22 | 4.433 | 5.008 | 5.203 | 5.258 | 5.305 | 0.994 | 1.000 | 7.808 |
| ASV_512 | 4.980 | 5.248 | 46 | 2 | 45.85 | 0.008 | 3.84 | 4.980 | 5.187 | 5.336 | 5.937 | 6.445 | 0.990 | 0.986 | 4.157 |
| ASV_96 | 5.315 | 5.258 | 27 | 2 | 32.53 | 0.008 | 4.18 | 5.245 | 5.258 | 5.315 | 5.490 | 5.656 | 0.986 | 1.000 | 4.365 |
| ASV_215 | 5.428 | 5.395 | 28 | 2 | 34.36 | 0.004 | 5.40 | 5.258 | 5.380 | 5.425 | 5.443 | 5.473 | 0.984 | 1.000 | 5.242 |
| ASV_31 | 4.328 | 4.433 | 96 | 2 | 86.77 | 0.004 | 6.65 | 4.335 | 4.378 | 4.660 | 5.200 | 5.233 | 0.966 | 0.994 | 7.430 |

**Table S20** (continued)

**B. Bacteria direct interactors associated with percent carbon**

| ASV | ienv.cp | zenv.cp | freq | maxgrp | IndVal | obsiv.prob | zscore | 5% | 10% | 50% | 90% | 95% | purity | reliability | z.median |
|---|---|---|---|---|---|---|---|---|---|---|---|---|---|---|---|
| ASV_31 | 3.900 | 3.460 | 96 | 1 | 83.09 | 0.004 | 7.18 | 1.199 | 2.475 | 3.460 | 3.960 | 3.995 | 1.000 | 1.000 | 7.449 |
| ASV_61 | 2.370 | 1.680 | 64 | 1 | 67.44 | 0.004 | 7.25 | 0.790 | 0.990 | 1.470 | 1.720 | 2.090 | 1.000 | 1.000 | 7.583 |
| ASV_397 | 0.895 | 0.895 | 38 | 1 | 50.15 | 0.004 | 5.19 | 0.770 | 0.780 | 0.895 | 1.720 | 1.945 | 1.000 | 1.000 | 5.570 |
| ASV_402 | 2.305 | 2.370 | 68 | 1 | 66.97 | 0.004 | 5.69 | 0.895 | 0.915 | 1.800 | 2.385 | 2.435 | 1.000 | 1.000 | 6.888 |
| ASV_442 | 0.990 | 1.035 | 43 | 1 | 53.55 | 0.004 | 7.87 | 0.925 | 0.945 | 0.990 | 1.040 | 1.050 | 1.000 | 1.000 | 8.542 |
| ASV_503 | 0.695 | 0.705 | 28 | 1 | 71.92 | 0.004 | 8.73 | 0.635 | 0.640 | 0.695 | 1.395 | 1.520 | 1.000 | 1.000 | 8.517 |
| ASV_663 | 0.800 | 0.790 | 35 | 1 | 52.49 | 0.004 | 7.61 | 0.780 | 0.790 | 0.810 | 1.180 | 1.860 | 1.000 | 1.000 | 7.835 |
| ASV_625 | 2.325 | 0.955 | 28 | 1 | 29.23 | 0.004 | 4.23 | 0.855 | 0.870 | 0.960 | 2.008 | 2.325 | 1.000 | 0.978 | 5.091 |
| ASV_28 | 3.900 | 3.900 | 88 | 1 | 89.80 | 0.004 | 7.10 | 2.469 | 3.060 | 3.900 | 3.995 | 4.060 | 0.996 | 1.000 | 6.859 |
| ASV_103 | 2.840 | 2.370 | 57 | 1 | 56.48 | 0.004 | 4.69 | 0.910 | 0.930 | 2.370 | 2.840 | 3.242 | 0.996 | 0.996 | 4.655 |
| ASV_2 | 3.460 | 3.460 | 108 | 1 | 77.12 | 0.004 | 4.79 | 1.084 | 1.167 | 3.460 | 3.900 | 3.900 | 0.996 | 0.994 | 5.162 |
| ASV_89 | 3.900 | 2.325 | 71 | 1 | 64.17 | 0.004 | 4.34 | 0.730 | 0.879 | 1.680 | 2.813 | 3.900 | 0.992 | 0.998 | 4.693 |
| ASV_322 | 0.880 | 0.880 | 33 | 1 | 35.97 | 0.004 | 5.20 | 0.870 | 0.880 | 0.925 | 1.045 | 1.190 | 0.992 | 0.986 | 5.847 |
| ASV_69 | 1.265 | 1.265 | 49 | 2 | 59.62 | 0.004 | 6.43 | 0.895 | 1.200 | 1.265 | 1.485 | 1.500 | 0.998 | 1.000 | 7.106 |

**C. Bacteria diffuse interactors associated with pH**

| ASV | ienv.cp | zenv.cp | freq | maxgrp | IndVal | obsiv.prob | zscore | 5% | 10% | 50% | 90% | 95% | purity | reliability | z.median |
|---|---|---|---|---|---|---|---|---|---|---|---|---|---|---|---|
| ASV_77 | 5.380 | 5.693 | 33 | 1 | 45.98 | 0.004 | 9.05 | 5.340 | 5.365 | 5.575 | 5.713 | 5.760 | 1.000 | 1.000 | 9.536 |
| ASV_92 | 4.433 | 5.808 | 46 | 1 | 54.23 | 0.004 | 7.89 | 4.435 | 5.239 | 5.783 | 5.928 | 5.958 | 1.000 | 1.000 | 8.542 |
| ASV_149 | 4.433 | 5.380 | 34 | 1 | 78.57 | 0.004 | 15.09 | 5.340 | 5.365 | 5.480 | 5.573 | 5.620 | 1.000 | 1.000 | 15.417 |
| ASV_171 | 5.663 | 5.715 | 39 | 1 | 50.64 | 0.004 | 8.19 | 5.540 | 5.563 | 5.680 | 5.740 | 5.778 | 1.000 | 1.000 | 8.467 |
| ASV_191 | 5.333 | 5.315 | 44 | 1 | 59.18 | 0.004 | 7.40 | 5.285 | 5.303 | 5.354 | 5.765 | 6.120 | 1.000 | 1.000 | 7.924 |
| ASV_245 | 5.950 | 5.928 | 51 | 1 | 60.37 | 0.004 | 8.39 | 5.840 | 5.870 | 5.950 | 5.983 | 6.008 | 1.000 | 1.000 | 8.828 |
| ASV_249 | 5.950 | 5.808 | 37 | 1 | 47.00 | 0.004 | 9.04 | 5.725 | 5.750 | 5.828 | 5.973 | 5.995 | 1.000 | 1.000 | 8.703 |
| ASV_269 | 5.745 | 5.663 | 40 | 1 | 52.40 | 0.004 | 8.04 | 5.630 | 5.650 | 5.730 | 5.900 | 5.938 | 1.000 | 1.000 | 8.652 |
| ASV_307 | 4.433 | 5.380 | 33 | 1 | 45.78 | 0.004 | 7.55 | 4.430 | 4.433 | 5.380 | 5.708 | 5.765 | 1.000 | 1.000 | 8.024 |
| ASV_314 | 5.638 | 5.625 | 33 | 1 | 58.20 | 0.004 | 11.35 | 5.380 | 5.508 | 5.638 | 5.741 | 5.760 | 1.000 | 1.000 | 12.165 |
| ASV_414 | 4.515 | 5.395 | 35 | 1 | 52.99 | 0.004 | 8.20 | 5.035 | 5.048 | 5.353 | 5.460 | 5.655 | 1.000 | 1.000 | 8.877 |
| ASV_469 | 4.515 | 4.515 | 31 | 1 | 91.55 | 0.004 | 10.06 | 4.513 | 4.515 | 4.660 | 5.568 | 5.625 | 1.000 | 1.000 | 10.254 |
| ASV_800 | 5.333 | 5.715 | 30 | 1 | 44.30 | 0.004 | 9.71 | 5.325 | 5.345 | 5.565 | 5.745 | 5.765 | 1.000 | 1.000 | 10.744 |
| ASV_1042 | 4.433 | 5.048 | 32 | 1 | 74.00 | 0.004 | 11.02 | 4.838 | 5.035 | 5.213 | 5.680 | 5.760 | 1.000 | 1.000 | 11.824 |
| ASV_1084 | 6.448 | 5.725 | 31 | 1 | 33.66 | 0.004 | 4.49 | 5.480 | 5.523 | 5.740 | 6.206 | 6.448 | 1.000 | 1.000 | 5.344 |

**Table S20** (continued)

| ASV | ienv.cp | zenv.cp | freq | maxgrp | IndVal | obsiv.prob | zscore | 5% | 10% | 50% | 90% | 95% | purity | reliability | z.median |
|---|---|---|---|---|---|---|---|---|---|---|---|---|---|---|---|
| ASV_1115 | 5.380 | 5.380 | 28 | 1 | 53.15 | 0.004 | 9.44 | 5.267 | 5.325 | 5.523 | 5.655 | 5.708 | 1.000 | 1.000 | 10.340 |
| ASV_1436 | 5.915 | 5.855 | 33 | 1 | 43.52 | 0.004 | 7.93 | 5.693 | 5.715 | 5.860 | 5.935 | 5.960 | 1.000 | 1.000 | 8.009 |
| ASV_331 | 4.433 | 5.368 | 48 | 1 | 55.17 | 0.004 | 7.12 | 5.190 | 5.230 | 5.333 | 5.538 | 5.949 | 1.000 | 0.998 | 7.451 |
| ASV_570 | 5.563 | 5.523 | 41 | 1 | 46.99 | 0.004 | 5.71 | 5.463 | 5.498 | 5.663 | 6.163 | 6.185 | 0.998 | 1.000 | 6.808 |
| ASV_78 | 5.540 | 5.540 | 28 | 1 | 40.56 | 0.004 | 5.33 | 5.333 | 5.355 | 5.540 | 5.783 | 5.928 | 0.996 | 1.000 | 6.206 |
| ASV_306 | 5.783 | 5.765 | 42 | 1 | 49.54 | 0.004 | 7.27 | 5.680 | 5.710 | 5.810 | 5.990 | 6.073 | 0.994 | 1.000 | 8.050 |
| ASV_677 | 6.448 | 5.888 | 37 | 1 | 39.56 | 0.004 | 6.00 | 5.650 | 5.693 | 5.888 | 6.398 | 6.448 | 0.994 | 1.000 | 6.412 |
| ASV_193 | 5.523 | 5.600 | 38 | 1 | 45.81 | 0.004 | 6.44 | 5.395 | 5.505 | 5.580 | 5.885 | 6.103 | 0.994 | 0.998 | 6.972 |
| ASV_848 | 4.515 | 5.018 | 28 | 1 | 62.78 | 0.004 | 8.10 | 4.435 | 4.515 | 4.998 | 5.341 | 5.390 | 0.992 | 1.000 | 9.209 |
| ASV_944 | 6.205 | 5.888 | 30 | 1 | 33.49 | 0.004 | 4.32 | 5.490 | 5.568 | 5.890 | 6.205 | 6.245 | 0.988 | 0.994 | 4.996 |
| ASV_26 | 4.328 | 5.480 | 42 | 1 | 38.90 | 0.004 | 4.60 | 5.184 | 5.303 | 5.568 | 6.045 | 6.140 | 0.982 | 0.980 | 5.307 |
| ASV_963 | 5.315 | 5.315 | 29 | 1 | 34.69 | 0.012 | 4.37 | 5.285 | 5.295 | 5.428 | 6.045 | 6.075 | 0.980 | 0.986 | 5.344 |
| ASV_1139 | 5.680 | 5.680 | 39 | 1 | 46.15 | 0.004 | 6.22 | 5.600 | 5.620 | 5.723 | 5.930 | 5.983 | 0.976 | 1.000 | 6.664 |
| ASV_70 | 6.533 | 6.448 | 39 | 2 | 77.01 | 0.004 | 11.86 | 5.925 | 5.960 | 6.163 | 6.418 | 6.500 | 1.000 | 1.000 | 13.436 |
| ASV_86 | 5.315 | 5.315 | 44 | 2 | 50.91 | 0.008 | 6.14 | 5.275 | 5.295 | 5.656 | 6.208 | 6.251 | 1.000 | 1.000 | 7.530 |
| ASV_124 | 4.980 | 5.638 | 54 | 2 | 53.32 | 0.004 | 6.85 | 5.213 | 5.472 | 5.638 | 5.720 | 5.753 | 1.000 | 1.000 | 7.163 |
| ASV_200 | 6.868 | 5.915 | 32 | 2 | 46.03 | 0.004 | 7.68 | 5.443 | 5.710 | 5.915 | 6.245 | 6.445 | 1.000 | 1.000 | 8.445 |
| ASV_241 | 6.398 | 6.418 | 47 | 2 | 79.79 | 0.004 | 10.51 | 5.693 | 5.807 | 6.350 | 6.428 | 6.533 | 1.000 | 1.000 | 10.423 |
| ASV_254 | 6.580 | 6.008 | 37 | 2 | 57.65 | 0.004 | 8.70 | 5.900 | 5.940 | 6.008 | 6.580 | 6.595 | 1.000 | 1.000 | 10.007 |
| ASV_330 | 5.248 | 5.248 | 45 | 2 | 51.10 | 0.004 | 5.88 | 5.212 | 5.230 | 5.505 | 6.190 | 6.280 | 1.000 | 1.000 | 6.663 |
| ASV_394 | 4.980 | 5.070 | 51 | 2 | 51.40 | 0.004 | 4.53 | 4.980 | 5.070 | 5.325 | 6.008 | 6.008 | 1.000 | 1.000 | 5.217 |
| ASV_513 | 5.333 | 5.725 | 32 | 2 | 37.09 | 0.004 | 5.79 | 5.315 | 5.333 | 5.725 | 6.103 | 6.280 | 1.000 | 1.000 | 7.317 |
| ASV_644 | 6.580 | 5.938 | 29 | 2 | 42.63 | 0.004 | 8.42 | 5.783 | 5.940 | 6.121 | 6.580 | 6.585 | 1.000 | 1.000 | 9.393 |
| ASV_650 | 6.823 | 6.580 | 28 | 2 | 64.38 | 0.004 | 8.75 | 6.333 | 6.415 | 6.580 | 6.723 | 6.795 | 1.000 | 1.000 | 9.984 |
| ASV_1292 | 6.533 | 6.025 | 27 | 2 | 42.90 | 0.004 | 7.16 | 5.595 | 5.693 | 6.025 | 6.533 | 6.548 | 1.000 | 1.000 | 8.932 |
| ASV_949 | 6.580 | 6.498 | 29 | 2 | 46.75 | 0.004 | 6.95 | 5.708 | 5.865 | 6.498 | 6.585 | 6.615 | 1.000 | 0.998 | 6.853 |
| ASV_1078 | 6.840 | 5.855 | 29 | 2 | 34.00 | 0.004 | 5.57 | 5.395 | 5.408 | 5.783 | 6.398 | 6.833 | 1.000 | 0.998 | 6.644 |
| ASV_1369 | 5.443 | 5.443 | 28 | 2 | 36.00 | 0.004 | 5.61 | 5.413 | 5.428 | 5.523 | 5.681 | 5.715 | 0.992 | 1.000 | 5.514 |
| ASV_666 | 5.333 | 5.315 | 31 | 2 | 34.90 | 0.008 | 4.14 | 5.187 | 5.230 | 5.366 | 5.443 | 5.961 | 0.986 | 1.000 | 4.623 |
| ASV_1587 | 5.213 | 5.258 | 46 | 2 | 47.95 | 0.004 | 5.12 | 5.185 | 5.203 | 5.325 | 5.625 | 5.680 | 0.982 | 1.000 | 5.470 |
| ASV_109 | 4.980 | 5.213 | 63 | 2 | 57.88 | 0.004 | 5.06 | 4.868 | 4.908 | 5.198 | 5.248 | 5.255 | 0.974 | 1.000 | 5.426 |

**Table S20** (continued)

**D. Fungi direct interactors associated with MAP**

| ASV | ienv.cp | zenv.cp | freq | maxgrp | IndVal | obsiv.prob | zscore | 5% | 10% | 50% | 90% | 95% | purity | reliability | z.median |
|---|---|---|---|---|---|---|---|---|---|---|---|---|---|---|---|
| ASV_179 | 1146.96 | 1166.37 | 15 | 1 | 44.12 | 0.004 | 10.52 | 1127.55 | 1127.55 | 1146.96 | 1166.37 | 1166.37 | 1.000 | 1.000 | 10.979 |
| ASV_291 | 1116.87 | 1122.21 | 20 | 1 | 69.48 | 0.004 | 12.25 | 1116.87 | 1116.87 | 1122.21 | 1127.55 | 1127.55 | 1.000 | 1.000 | 12.385 |
| ASV_134 | 1127.55 | 1192.21 | 36 | 1 | 42.85 | 0.004 | 4.88 | 1116.87 | 1122.21 | 1146.96 | 1200.03 | 1200.03 | 1.000 | 1.000 | 5.783 |
| ASV_370 | 1116.87 | 1127.55 | 15 | 1 | 35.17 | 0.004 | 6.18 | 1116.87 | 1116.87 | 1127.55 | 1169.34 | 1176.24 | 0.996 | 0.988 | 6.432 |
| ASV_443 | 1127.55 | 1166.37 | 14 | 1 | 27.80 | 0.004 | 6.19 | 1116.87 | 1122.21 | 1166.37 | 1167.86 | 1169.34 | 0.996 | 0.970 | 6.471 |
| ASV_26 | 1116.87 | 1166.37 | 18 | 1 | 29.60 | 0.004 | 3.91 | 1116.87 | 1116.87 | 1127.55 | 1167.86 | 1169.34 | 0.994 | 0.962 | 5.176 |
| ASV_240 | 1116.87 | 1116.87 | 11 | 1 | 55.17 | 0.004 | 9.83 | 1116.87 | 1116.87 | 1116.87 | 1127.55 | 1127.55 | 0.954 | 0.992 | 8.443 |
| ASV_113 | 1279.75 | 1279.75 | 24 | 2 | 68.76 | 0.004 | 6.58 | 1169.34 | 1176.24 | 1192.21 | 1279.75 | 1300.15 | 1.000 | 1.000 | 7.037 |
| ASV_107 | 1279.75 | 1279.75 | 11 | 2 | 45.72 | 0.004 | 7.49 | 1177.02 | 1177.02 | 1192.21 | 1300.15 | 1300.15 | 1.000 | 0.988 | 6.045 |
| ASV_169 | 1184.62 | 1184.62 | 11 | 2 | 19.39 | 0.004 | 4.04 | 1167.86 | 1169.34 | 1192.21 | 1192.21 | 1192.21 | 1.000 | 0.974 | 4.373 |
| ASV_248 | 1184.62 | 1184.62 | 12 | 2 | 23.67 | 0.004 | 5.31 | 1177.02 | 1177.02 | 1192.21 | 1192.21 | 1192.21 | 0.998 | 0.990 | 6.063 |

**E. Fungi direct interactors associated with percent nitrogen**

| ASV | ienv.cp | zenv.cp | freq | maxgrp | IndVal | obsiv.prob | zscore | 5% | 10% | 50% | 90% | 95% | purity | reliability | z.median |
|---|---|---|---|---|---|---|---|---|---|---|---|---|---|---|---|
| ASV_179 | 0.040 | 0.060 | 15 | 1 | 27.78 | 0.004 | 5.70 | 0.040 | 0.040 | 0.050 | 0.060 | 0.060 | 1.000 | 1.000 | 6.529 |
| ASV_5 | 0.160 | 0.120 | 66 | 1 | 67.87 | 0.004 | 6.31 | 0.060 | 0.060 | 0.110 | 0.160 | 0.160 | 1.000 | 1.000 | 6.877 |
| ASV_254 | 0.025 | 0.040 | 14 | 1 | 27.09 | 0.004 | 6.02 | 0.025 | 0.030 | 0.048 | 0.050 | 0.060 | 1.000 | 1.000 | 6.977 |
| ASV_38 | 0.025 | 0.030 | 41 | 1 | 57.26 | 0.004 | 4.98 | 0.025 | 0.030 | 0.035 | 0.045 | 0.050 | 1.000 | 0.998 | 6.352 |
| ASV_231 | 0.030 | 0.030 | 17 | 1 | 30.28 | 0.004 | 4.29 | 0.030 | 0.030 | 0.040 | 0.070 | 0.090 | 1.000 | 0.998 | 6.102 |
| ASV_209 | 0.030 | 0.030 | 15 | 1 | 47.48 | 0.004 | 7.30 | 0.025 | 0.030 | 0.045 | 0.050 | 0.060 | 1.000 | 0.994 | 6.964 |
| ASV_248 | 0.030 | 0.030 | 12 | 1 | 38.88 | 0.004 | 7.89 | 0.030 | 0.030 | 0.030 | 0.040 | 0.045 | 0.998 | 0.976 | 7.180 |
| ASV_107 | 0.030 | 0.030 | 11 | 1 | 30.31 | 0.004 | 6.38 | 0.020 | 0.025 | 0.030 | 0.056 | 0.070 | 0.996 | 0.974 | 6.958 |
| ASV_240 | 0.180 | 0.160 | 11 | 2 | 37.77 | 0.004 | 8.89 | 0.150 | 0.160 | 0.180 | 0.240 | 0.260 | 0.994 | 0.994 | 9.363 |
| ASV_319 | 0.260 | 0.160 | 16 | 2 | 38.19 | 0.004 | 7.22 | 0.090 | 0.090 | 0.123 | 0.195 | 0.240 | 0.992 | 0.998 | 8.296 |
| ASV_291 | 0.180 | 0.180 | 20 | 2 | 53.36 | 0.004 | 7.83 | 0.080 | 0.090 | 0.170 | 0.185 | 0.190 | 0.986 | 0.992 | 7.455 |
| ASV_56 | 0.220 | 0.110 | 40 | 2 | 54.48 | 0.004 | 5.24 | 0.040 | 0.085 | 0.110 | 0.240 | 0.260 | 0.952 | 0.960 | 5.937 |

**F. Fungi diffuse interactors associated with the max height of host plant**

| ASV | ienv.cp | zenv.cp | freq | maxgrp | IndVal | obsiv.prob | zscore | 5% | 10% | 50% | 90% | 95% | purity | reliability | z.median |
|---|---|---|---|---|---|---|---|---|---|---|---|---|---|---|---|
| ASV_324 | 2.605 | 2.3 | 14 | 2 | 21.61 | 0.004 | 3.61 | 2.195 | 2.2 | 2.3 | 2.605 | 2.605 | 0.982 | 0.972 | 4.918 |
